# DNA origami vaccine (DoriVac) nanoparticles improve both humoral and cellular immune responses to infectious diseases

**DOI:** 10.1101/2023.12.29.573647

**Authors:** Yang C. Zeng, Olivia J. Young, Qiancheng Xiong, Longlong Si, Min Wen Ku, Sylvie G. Bernier, Hawa Dembele, Giorgia Isinelli, Tal Gilboa, Zoe Newell Swank, Su Hyun Seok, Anjali Rajwar, Amanda Jiang, Yunhao Zhai, LaTonya D. Williams, Caleb A. Hellman, Chris M. Wintersinger, Amanda R. Graveline, Andyna Vernet, Melinda Sanchez, Sarai Bardales, Georgia D. Tomaras, Ju Hee Ryu, Ick Chan Kwon, Girija Goyal, Donald E. Ingber, William M. Shih

**Author notes:** These authors contributed equally to this work (Yang C. Zeng, Olivia J. Young, Qiancheng Xiong, Longlong Si, Min Wen Ku).

## Abstract

Current SARS-CoV-2 vaccines have demonstrated robust induction of neutralizing antibodies and CD4^+^ T cell activation, however CD8^+^ responses are variable, and the duration of immunity and protection against variants are limited. Here we repurposed our DNA origami vaccine nanotechnology, DoriVac, for targeting infectious viruses, namely SARS-CoV-2, HIV, and Ebola. The DNA origami nanoparticle, conjugated with infectious-disease-specific heptad repeat 2 (HR2) peptides, which act as highly conserved antigens, and CpG adjuvant at precise nanoscale spacing, induced neutralizing antibodies, Th1 CD4^+^ T cells, and CD8^+^ T cells in naïve mice, with significant improvement over a bolus control. Pre-clinical studies using lymph-node-on-a-chip systems validated that DoriVac, when conjugated with antigenic peptides or proteins, induced promising cellular and humoral immune responses in human cells. Moreover, DoriVac bearing full-length SARS-CoV-2 spike protein achieved immune responses comparable to current mRNA vaccine platforms while potentially reducing storage constraints. These results suggest that DoriVac holds potential as a versatile, modular vaccine platform, capable of inducing both humoral and cellular immunities, underscoring its potential utility in addressing future pandemics.

## Introduction

The coronavirus disease 2019 pandemic highlighted the need for swift vaccine development. Initial focus on rapid vaccine design for pandemics^1-5^ led to the novel mRNA vaccines, mRNA-1273 (manufactured by Moderna) and BNT162b2 (manufactured by Pfizer), which rely on lipid nanoparticle delivery of mRNA encoding an early variant of the spike protein. Despite their success, the emergence of SARS-CoV-2 variants like B.1.351 (Beta)^6^, B.1.617.2 (Delta)^7^, and B.1.529 (Omicron)^8^ raised concerns about the vaccine efficacy as variants demonstrated the ability to evade immunity^9-15^. The immune evasion observed with current vaccines necessitates interventions effective against mutations.

Current SARS-CoV-2 vaccines focus on the receptor binding domain (RBD) of the spike protein. Viruses rely on RBD to bind to cells and initiate infection, and then heptad repeat 1 (HR1) and heptad repeat 2 (HR2) to fuse the viral and cell membranes. HR1 and HR2, conserved across various viruses, self-assemble into α-helical coils, and then assemble into superhelical structures to facilitate fusion^16-21^. While the RBD region and other viral regions are subject to viral evolution, HR1 and HR2 sequences remain highly conserved, providing a conserved antigen for vaccines^22^. Only three amino acids differ between the original SARS-CoV-2 HR1 sequence and the Omicron variant (**Supplementary Table 1**). HR1 and HR2 also harbor T cell epitopes and may induce neutralizing antibodies, serving as viable antigens for vaccine design^23,24^. HR2, with a simpler structure than HR1, has been successfully targeted by vaccines,^22,25^ and was selected as the antigen for delivery via our vaccine nanotechnology for SARS-CoV-2, HIV, and Ebola (**Supplementary Table 2**).

While the vaccine community traditionally prioritized neutralizing antibody responses^26^, there is now growing acknowledgement of the essential role of cellular immune responses (dendritic cells, CD4^+^ and CD8^+^ T cells) for broad viral protection^9,24,27-33^. Functional T cells prevent immune escape of mutated strains^9^. SARS-CoV-2 mutated strains have been demonstrated to escape neutralizing antibody responses, but not T cell responses^34^. CD4^+^ T cells support antibody generation^35^, and studies show that CD4^+^ T cell transfer can protect against viral challenge^36^. Mild SARS-CoV-2 infections exhibit robust CD8^+^ T cell reactivity^37,38^ contributing to rapid viral clearance^29^. Depleting CD8^+^ T cells in non-human primates increases susceptibility to SARS-CoV-2 re-infection^39^. In HIV and Ebola, CD8^+^ T cells are crucial for long-term control and vaccine-induced protection. CD8^+^ depletion led to failure in controlling simian immunodeficiency virus in non-human primates^40,41^. In Ebola, CD8^+^ cells were essential for immune protection in non-human primates, while antibody transfer failed to protect^42^. An ideal vaccine should induce both humoral and cellular immune responses, including neutralizing antibodies and long-term memory T cells^9^. mRNA vaccines demonstrate robust CD4^+^ responses, but variable CD8^+^ responses; both influence long-term immunity^43-47^.

Multiple vaccine candidates have been developed to induce neutralizing antibodies and cellular responses against the SARS-CoV-2 spike protein, including over 60 different nanoparticle formulations^48,49^. Despite the success of lipid nanoparticle-based mRNA vaccines, these vaccines face challenges like manufacturing complexity, cold-chain requirements, limited stability, high cost, poor cargo loading efficiency, limited control over cargo stoichiometry, and off-target effects^48,50^. Here, we introduce DoriVac, a DNA origami vaccine nanoparticle, as a versatile nanotechnology for infectious disease. While previous studies have demonstrated vaccine delivery with DNA origami for cancer, this study aims to demonstrate its broad applicability for infectious diseases. DoriVac induced robust humoral and T cell immune responses against SARS-CoV-2, HIV, and Ebola viruses in mouse models, demonstrating the nanotechnology’s programmability for various infectious-disease HR2 antigens. This approach may broadly apply to pathogen vaccine development by conjugating the respective antigens to the DNA origami. Our current study also includes a full-length spike protein-conjugation strategy that elicited immune responses comparable to state-of-the-art mRNA-based SARS-CoV-2 vaccines, generating rapid early antibody responses after one dose, further highlighting DoriVac’s promise as a next-generation, broadly applicable vaccine platform.

### Fabrication of modular DoriVac nanoparticles

We previously developed a DNA origami nanoparticle, termed square block (SQB), for its square-lattice architecture for precise spatial presentation of CpG adjuvants^52^. Formed through self-assembly of a long scaffold strand with corresponding short ‘staple’ strands, DoriVac is easy to manufacture and highly stable without cold-chain requirements, exhibits high occupancy of designed cargo-binding sites due to the robustness of DNA hybridization, and offers precise control over cargo attachment. This nanotechnology facilitates optimized spatial arrangement of immune-activating adjuvants, resulting in robust cellular immune responses in various cancers, as previously published^52^. The SQB flat face, modified with 18 CpG strands at 3.5 nm spacing, induces type I (Th1) skewed immune activation (**Fig. 1a**)^52^.

**Fig. 1.**
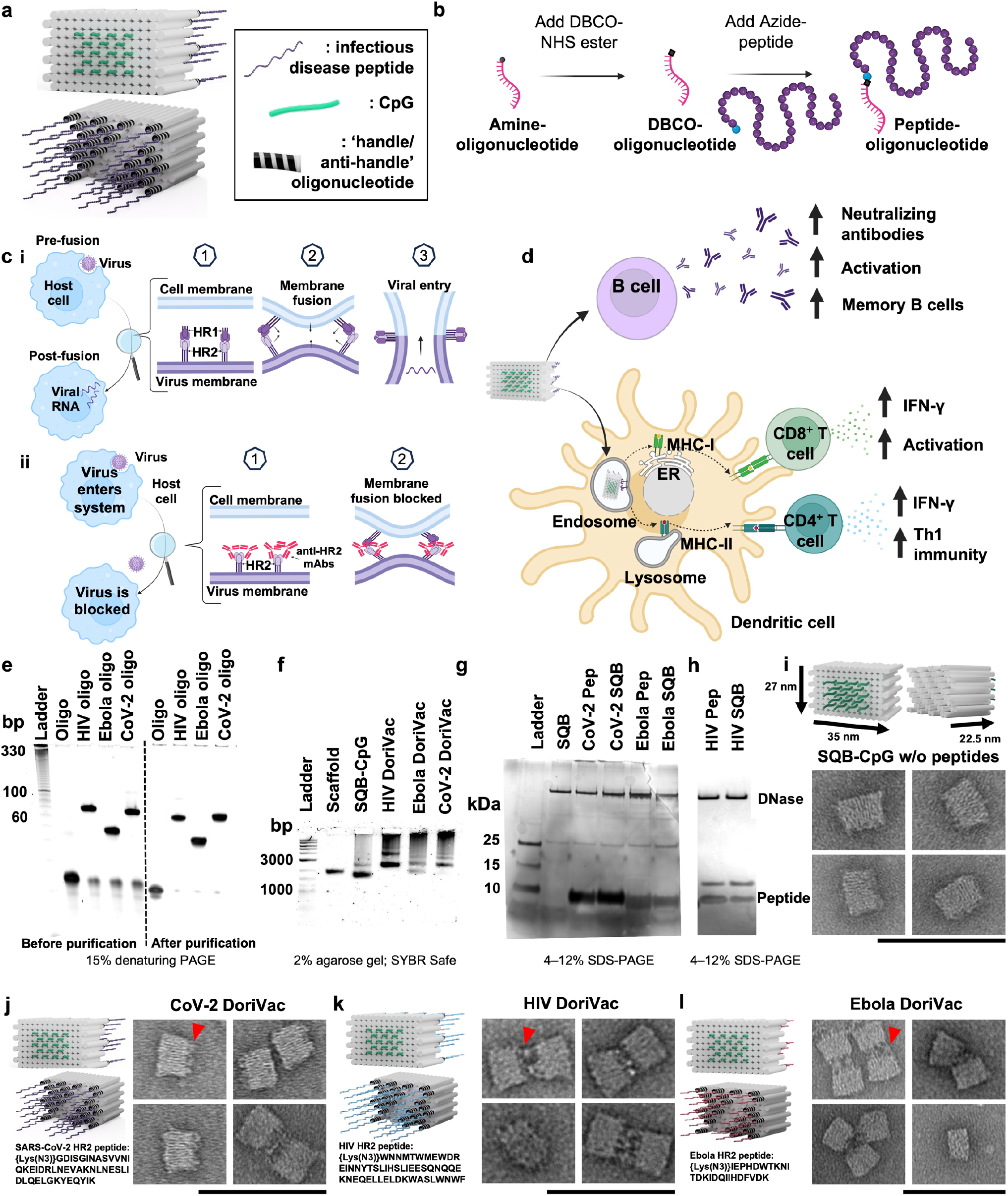
DNA origami vaccines (DoriVac) were fabricated with infectious-disease-specific peptides. **a**, Schematic of DoriVac, consisting of a DNA origami square block (SQB) nanoparticle conjugated with CpG at precise spacing of 3.5 nm and with infectious disease-specific peptides. **b**, Schematic demonstrated conjugation of DBCO**-**modified-oligonucleotide to an azide-modified peptide via copper-free click chemistry. **c.i**, A schematic showing how HR2 protein mediates virus-host fusion; HR2 can serve as a conserved target for infectious-disease vaccines. **ii**, Schematic showing how production of anti-HR2 antibodies (mAbs) prevents virus-host cell-membrane fusion and thereby inhibits viral infection. **d**, Schematic of DNA origami SQB nanoparticles delivering antigen and adjuvant at a precise spacing to antigen presenting cells, eliciting both humoral and cellular immune responses. **e**, Denaturing PAGE gel demonstrating successful conjugation and purification of infectious-disease**-**specific HR2 peptides to anti-handle oligonucleotides (“oligo”). **f**, Agarose gel demonstrating successful conjugation of oligo-HR2 peptides to the SQB DNA origami nanoparticles after removing the unconjugated cargos through PEG purification. **g–h**, SDS-PAGE gel demonstrating the results after DNase I digestion of infectious disease-specific HR2 peptides alone or conjugated with SQBs. DNase I digestion of the SQBs, followed by the analysis of the conjugated peptides using silver staining, confirms the successful peptide conjugation on the SQBs. DNase I sometimes produced a fainter secondary band with higher mobility. Gels with only DNase I and HIV-HR2 peptide are shown in Supplementary Fig. 1. **i-l**, Proposed schematics of the SQBs conjugated with CpGs (SQB-CpG) and SQBs conjugated with CpGs (18 copies) and HR2 peptides (24 copies; CoV-2-HR2 DoriVac, HIV-HR2 DoriVac, Ebola-HR2 DoriVac), respectively, and their representative TEM images. In TEM side views, the SQB exhibits two distinct faces: the flat interface is designated for CpG attachment, whereas the opposite face is reserved for peptide conjugation. The peptide-conjugated side is indicated by a red arrow in the first inset of each image set. The specific HR2 peptide sequences associated with each infectious disease are listed. Scale bar: 100 nm.

We applied DoriVac technology to create vaccines for SARS-CoV-2, HIV, and Ebola viruses by linking highly conserved viral HR2 peptides to the extruding face of the SQB nanoparticles (**Fig. 1a**). HR2 peptides contain MHC-I and MHC-II epitopes, which are crucial for broadly activating cellular immunity. To this end, we designed peptide-oligonucleotide conjugates with the appropriate “anti-handle” strand through DBCO-Azide click chemistry for specific attachment onto 24 specific “handle” sites of the extruding face of the SQB (**Fig. 1a,b**). The SQB nanoparticle co-delivers CpG adjuvant and disease-specific HR2 antigens to antigen presenting cells. Notably, the CpG sequence used in DoriVac remains unchanged from our previous studies, where it was shown to induce robust Th1-skewed immune activation in mouse models^52^. B cells produce neutralizing antibodies, which can block the membrane fusion of the virus with the host cell (**Fig. 1c**). Dendritic cells (DCs) present and cross-present the antigens to activate both CD4^+^ and CD8^+^ T cells (**Fig. 1d**). The oligonucleotide-HR2-peptide conjugates were purified via PAGE purification (**Fig. 1e**). The agarose-gel electrophoresis band shift demonstrates successful fabrication of peptide-functionalized SQB (**Fig. 1f**). To confirm peptide conjugation efficiency to the SQB, we digested the DNA origami via DNase I and estimated peptide occupancy of greater than 95% of the conjugation sites via silver stain (**Fig. 1g–h, Supplementary Table 3**). Fabrication of SARS-CoV-2-HR2, HIV-HR2, and Ebola-HR2 DoriVac was verified via TEM (**Fig. 1i-l, Supplementary Fig. 1**). Aggregation was observed via agarose gel, especially in the case of the HIV and Ebola SQBs of which the majority are dimers, possibly due to hydrophobic peptide interactions.

### DoriVac induces robust humoral immune responses

Having fabricated the vaccine, we evaluated DoriVac’s efficacy for induction of both humoral and cellular immune responses *in vivo*. Naïve mice were administered 20 pmol of HR2-fabricated DoriVac, comprising 0.36 nmol (2.2 μg) of CpG and 0.48 nmol of antigen (1.5 – 3.2 μg) (**Fig. 2a**). Two subcutaneous doses of DoriVac were given on day 0 and day 20, compared to bolus vaccine consisting of free CpG adjuvant and HR2 peptide. Blood samples were collected on day 14 and 28 for peripheral blood mononuclear cells (PBMCs) and plasma processing. On day 21, half of the mice from each group were sacrificed for immune cell profiling (**Supplementary Table 4-6**). On day 35, the remaining mice were sacrificed for immune cell profiling.

**Fig. 2.**
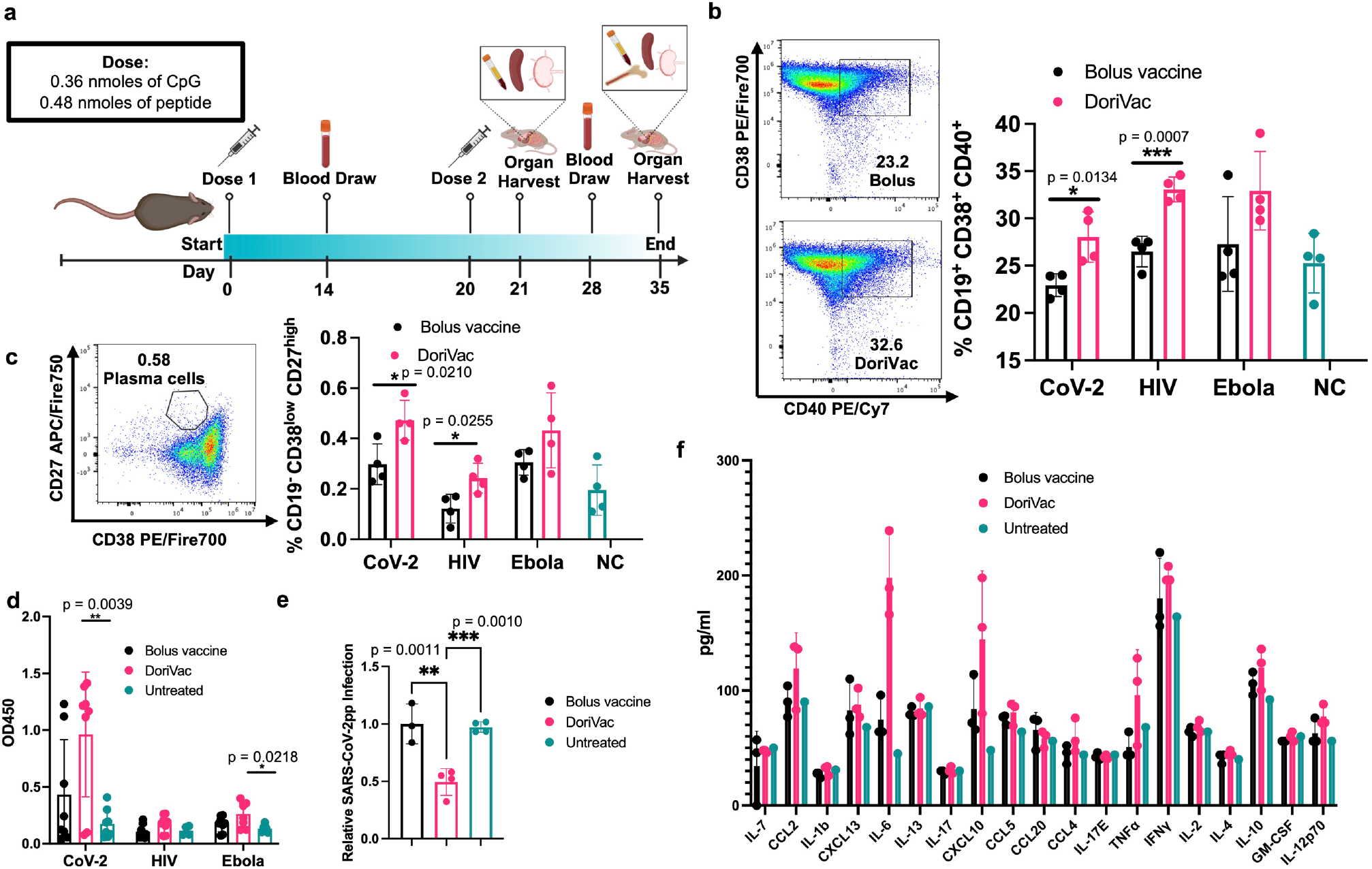
Immune profiling reveals the DoriVac elicits improved neutralizing antibody responses compared to a bolus vaccine. **a**, Schematic delineating the vaccine administration protocol for naïve C57BL/6 mice and the data collection timeline. Lymph nodes (LNs) and spleens were collected on day 21 and day 35 after sacrificing the mice for flow cytometry and ELISpot analysis. Plasma was collected on Days 14 and 28 for anti-HR2 antibody quantification and pseudovirus neutralization assays. Bone marrow and heart blood was collected at the conclusion of the study, day 35, to analyze B cell markers. **b**, B cells collected from the blood demonstrated increased markers of activation and antigen-presentation capabilities (CD40) and memory capacity (CD38) after two doses of DoriVac treatment (n=4) as determined by flow cytometry on day 21. NC refers to negative control. **c**, B cells collected from the blood on Day 21 demonstrated increased plasma-memory-cell population as evidenced by the increased CD19^-^ CD38^low^ CD27^high^ subpopulation as determined by flow cytometry (n=4). **d**, DoriVac treatment enhanced HR2-specific IgG antibody production as evidenced via ELISA assay, after two doses of DoriVac (on Day 35) compared to a bolus vaccine of free peptide and free CpG. Samples were diluted 1:100 before quantification. Data has been normalized (n=8). **e**, SARS-CoV-2 pseudovirus (SARS-CoV-2pp) neutralization assay (n=3-4, 1:100 dilution, Day 28) in model cell line ACE2-293T (n=4). **f**, Plasma was collected four hours after the first treatment dose on Day 0. The inflammatory cytokine response was quantified via Luminex ELISA assay (Bio-Plex Pro Mouse Cytokine 20-plex Assay (Bio-Rad)) (n=3 for treated groups; n=1 for negative (i.e. untreated) control). Data are represented as mean ± SD. The pseudovirus and ELISA data were analyzed by one-way ANOVA (with correction for multiple comparisons using a Tukey test) and significance was defined as a multiplicity-adjusted p value less than 0.05. The flow data were analyzed by multiple unpaired t-tests and significance was defined as a two-tailed p value less than 0.05. ‘*’ refers to p≤ 0.05; ‘**’ refers to p≤ 0.01; ‘***’ refers to p ≤ 0.001; Bars without asterisks indicate no statistically significant difference except for f (p > 0.05).

On day 21, B cells from PBMCs exhibited increased CD40 expression, a marker of activation and antigen-presentation capacity, in all three DoriVac treatment groups (**Fig. 2b**), surpassing the bolus vaccine, suggesting that DoriVac is superior in inducing B cell activation. Day 35 revealed a heightened plasma memory B cell population in the bone marrow, as evidenced by an increased CD19^low^ CD38^low^ CD27^high^ subpopulation after DoriVac treatment (**Fig. 2c**), despite unchanged overall B cell numbers (**Supplementary Figs. 2–3**). SARS-CoV-2-HR2-DoriVac treatment induced elevated HR2 peptide-specific IgG1 antibody responses, as quantified via ELISA, compared to the bolus vaccine (**Fig. 2d, Supplementary Fig. 4**). Neutralizing antibodies harvested from SARS-CoV-2-HR2-DoriVac groups significantly reduced infection in a SARS-CoV-2 pseudovirus (SARS-CoV-2pp) assay (**Fig. 2e**). In contrast, we did not observe neutralization of the pseudovirus for HIV and Ebola in our assay, possibly due to the weak immunogenicity of the antigens associated with these viruses (**Supplementary Tables 7–10**). We did observe modest antigen-specific IgG1 responses for HIV and Ebola after HIV-HR-DoriVac and Ebola-HR2-DoriVac treatment, respectively (**Fig. 2d, Supplementary Fig. 4**). We examined initial cytokine responses four hours post the first vaccine dose to naïve mice (**Fig. 2f**). Type 1 cytokines (TNFα, IL-2, IFNγ, IL-12) were slightly elevated, while type 2 cytokines (IL-4, IL-10) exhibited no obvious elevation after DoriVac treatment compared to the bolus vaccine group and the untreated mice^57^. Overall, these findings affirm DoriVac’s superior induction of humoral immune responses compared to those induced by a bolus vaccine, demonstrating its effectiveness in reducing the infection rate of SARS-CoV-2 pseudovirus.

### DoriVac induces DC activation

To ensure enduring immune protection against viral variants, a vaccine should stimulate both humoral and cellular immune responses. We first checked the DCs, which serve as a link between innate and adaptive immune responses. On day 21 — one day after the second vaccine dose— half of the mice were sacrificed, and the draining lymph nodes near the injection site were collected for flow (**Supplementary Fig. 5**). DoriVac increased the overall DC population (**Fig. 3a**) and activated DCs (CD11c^+^ CD86^+^) compared to the bolus vaccine (**Fig. 3b**). Plasmacytoid DCs (pDCs) are crucial in the anti-viral response in humans, secreting abundant type-1 interferon, fostering T cell activation and recruiting other immune cells^58^. The human pDC-like population (CD11c^+^ Gr-1^+^) significantly increased after DoriVac treatment (**Fig. 3c**), suggesting an increased anti-viral response. Activation markers MHC-II, PD-L1 and CD40 also increased after DoriVac administration (**Fig. 3d–f**). A notable rise was observed in the CD40^+^ DEC205^+^ DC population, indicating an increase in the activated, endocytic DC population (**Fig. 3g**). These results demonstrated DoriVac induced robust activation of DCs in healthy mice (**Supplementary Fig. 6**). Furthermore, co-delivery of SQB and HR2 peptide did not elicit the same level of DC activation as observed with the delivery of HR2 peptide-conjugated SQBs (**Supplementary Fig. 7**). This observation is consistent with the outcomes previously noted in DoriVac studies involving tumor-bearing mice^52^, further supporting the notion that the enhanced efficacy of the conjugated delivery system in activating DCs.

**Fig. 3.**
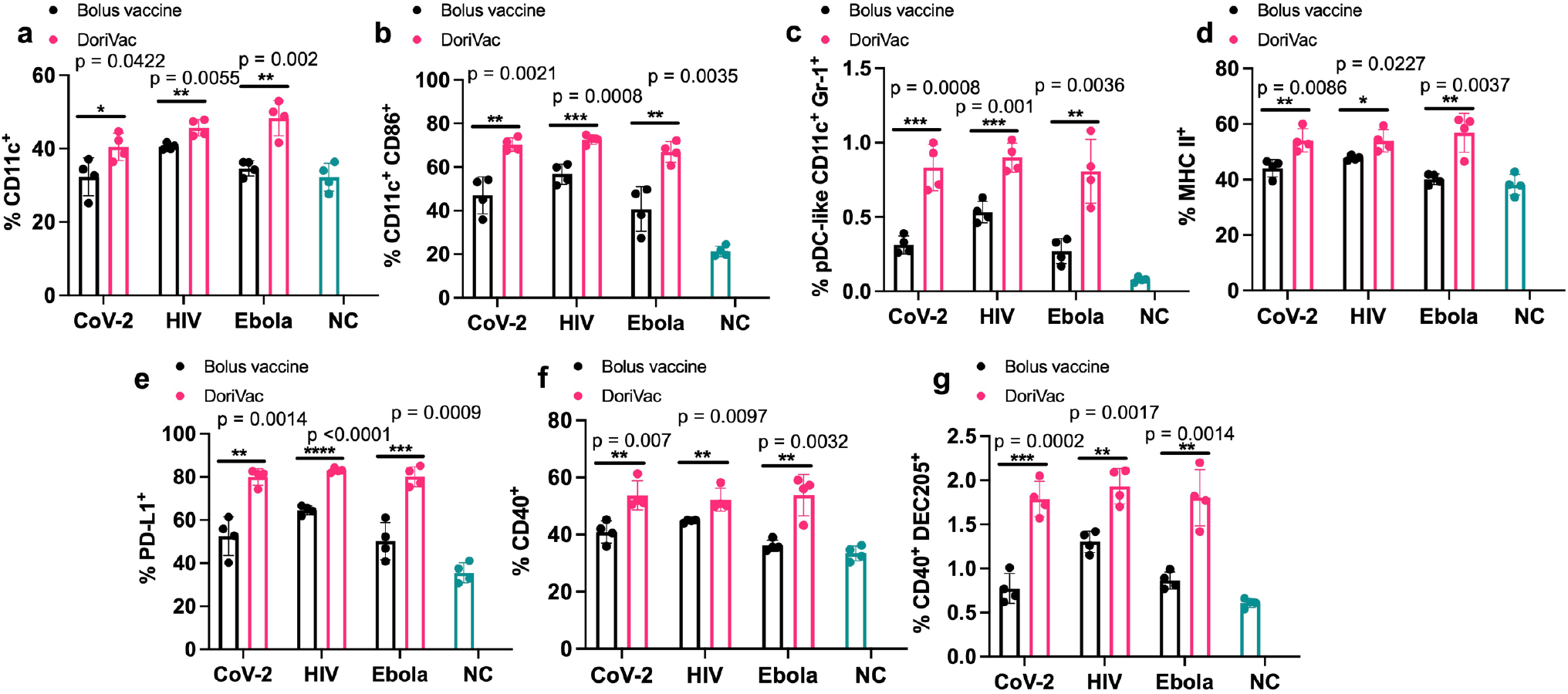
Immune profiling reveals DoriVac elicits superior antigen presenting cell responses compared to a bolus vaccine. C57BL/6 mice were treated with DoriVac (20 pmol) on Day 0 and Day 20. The mice were sacrificed on Day 21 and the draining lymph nodes (LNs) were processed into single-cell suspensions and analyzed by flow cytometry. **a**, Percentages of CD11c^+^ cells in the draining LNs (n=4) were quantified. NC means negative (i.e. untreated) control. **b**, Percentages of CD11c^+^ CD86^+^ DCs in the draining LNs (n=4) as determined by flow cytometry. **c**, Percentages of human plasmacytoid DC (pDC)-like (CD11c^+^ Gr-1^+^) DCs in the draining LNs (n=4) as determined by flow cytometry. **d**, Percentages of MHC-II^+^ DCs in the draining LNs (n=4) as determined by flow cytometry. **e**, Percentages of PD-L1^+^ population in the draining LNs (n=4) as determined by flow cytometry. **f**, Percentages of CD40^+^ population in the draining LNs (n=4) as determined by flow cytometry. **g**, Percentages of CD40^+^ DEC205^+^ population in DCs in the draining LNs (n=4) as determined by flow cytometry. DoriVac demonstrated a significant increase in DC activation compared to bolus-vaccine treatment. Data are represented as mean ± SD. The flow data were analyzed by multiple unpaired t-tests and significance was defined as a two-tailed p value less than 0.05. ‘*’ refers to p ≤ 0.05; ‘**’ refers to p ≤ 0.01; ‘***’ refers to p ≤ 0.001; ‘****’ refers to p ≤ 0.0001.

### DoriVac demonstrates activation of CD4^+^ T cells

In our previous study involving DoriVac in mouse cancer models^52^, we showed that DC activation by DoriVac leads to broad T cell activation. We aimed to confirm T cell induction by DoriVac in the context of viral antigens. Antigen-specific T cell activation was assessed via IFNγ ELISpot assay on splenocytes; our results demonstrated a significant increase in antigen-specific T cells after SARS-CoV-2-HR2 DoriVac administration (**Fig. 4a,b**). In contrast, HIV-HR2 DoriVac administration led to only a modest increase, and Ebola-HR2 DoriVac showed no apparent effect (**Fig. 4a,b; Supplementary Fig. 8**). We attribute this to the limited immunogenicity of the HR2 peptides used for HIV and Ebola, validated by their weaker predicted binding to MHC-I and MHC-II in both mice and humans via NetMHCpan-4.1^59^. In contrast, the SARS-CoV-2 HR2 peptide exhibited stronger binding predictions with six epitopes classified as strong binders, compared to zero to two epitopes for HIV and Ebola (**Supplementary Tables 7–10**). These findings underscore the importance of antigen selection in achieving effective antigen-specific T cell activation. Notably, these results confirmed that SARS-CoV-2-HR2 DoriVac induces significantly more antigen-specific T cells compared to the bolus vaccine.

**Fig. 4.**
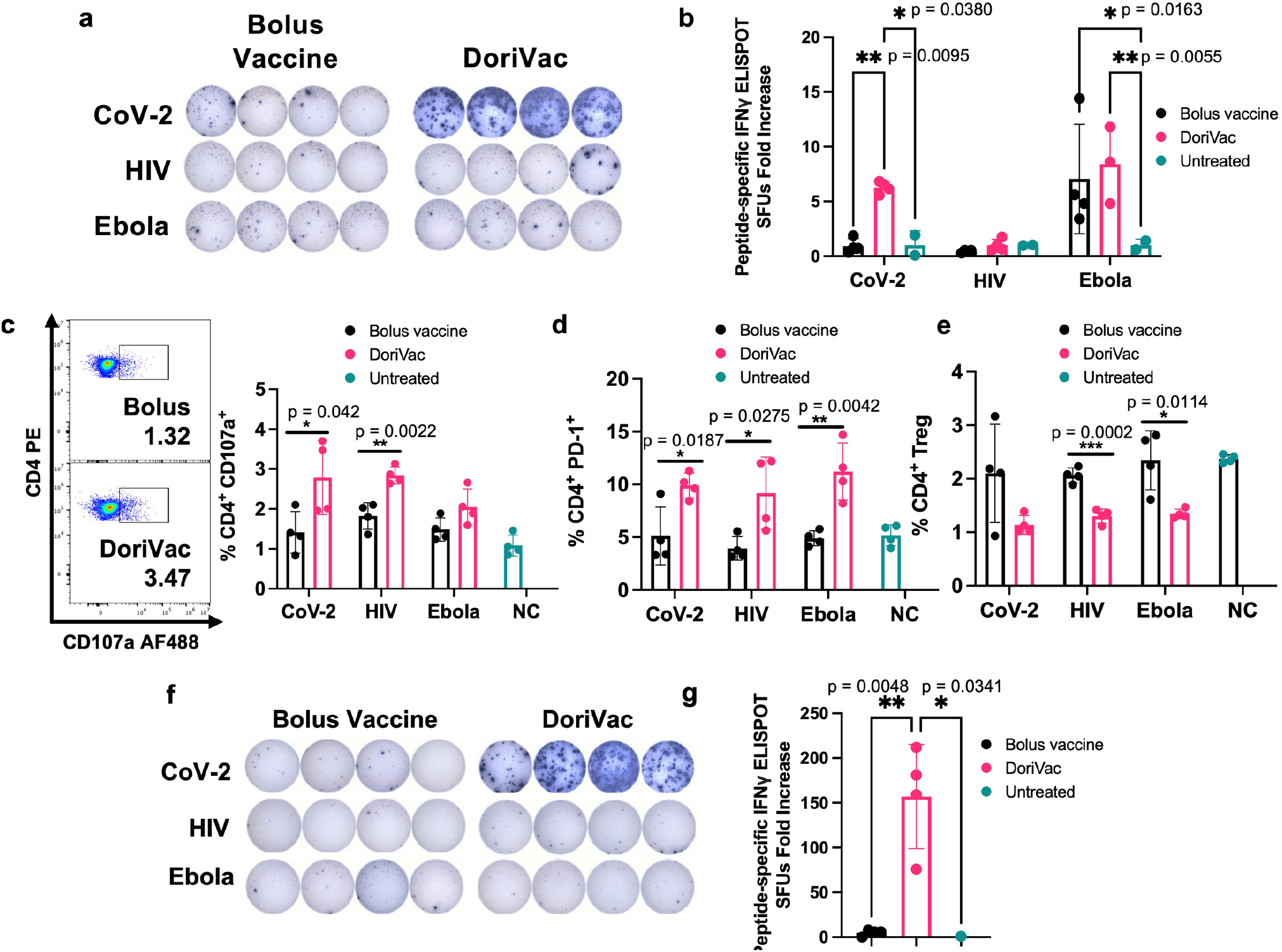
DoriVac induces enhanced Th1 CD4^+^ T cell activation in mice. **a**, IFNγ ELISpot demonstrating frequency of antigen-specific T cells in splenocytes on day 35 (n=4) after treatment with DoriVac compared with the bolus vaccine. **b**, Quantification of IFNγ ELISpot spot forming units (SFUs) demonstrates significant increase in SARS-COV-2 antigen-specific T cell frequency after treatment with DoriVac compared with the bolus vaccine and negative (i.e. untreated) control. **c**, Percentages of CD4^+^ CD107a^+^ T cells in the LN (n=4) as determined by flow cytometry and representative flow plots on day 35. NC means negative (i.e. untreated) control. **d**, Percentages of the LN CD4^+^ PD-1^+^ population (n=4) as determined by flow cytometry on day 21. **e**, Percentages of LN CD4^+^ T regulatory cell (Treg) population (n=4) as determined by flow cytometry on day 21. **f**, IFNγ ELISpot demonstrating frequency of CD4^+^ enriched antigen-specific splenocytes (n=4, day 35) after treatment with SARS-CoV-2-HR2 DoriVac. **g**, Corresponding quantification of IFNγ ELISpot spot forming units (SFUs). Data are represented as mean ± SD. The ELIspot data in **Fig. 4b** were analyzed by two-way ANOVA (with correction for multiple comparisons using Tukey’s test). The ELISpot data in **Fig. 4g** were analyzed by one-way ANOVA (with correction for multiple comparisons using Tukey’s test). In both analyses, statistical significance was defined as a multiplicity-adjusted p value less than 0.05. The flow data were analyzed by multiple unpaired t-tests and significance was defined as a two-tailed p value less than 0.05. ‘*’ refers to p ≤ 0.05; ‘**’ refers to p ≤ 0.01; Bars without asterisks indicate no statistically significant difference (p > 0.05).

Beyond overall T cell responses, we confirmed the presence of activated Th1 CD4^+^ T cells via flow cytometry (**Supplementary Fig. 9** for gating strategy). CD107a notably increased in the CD4^+^ T cell population, indicating enhanced CD4^+^ T cell activation and increased cytotoxic potential (**Fig. 4c**)^60^. PD-1 was also upregulated in the CD4^+^ T cell population, demonstrating increased activation (**Fig. 4d**). The T regulatory cell (Treg) population significantly decreased after treatment with DoriVac, suggesting reduced immunosuppression (**Fig. 4e**). Antigen-specific CD4^+^ T cell activation (CD8^+^ T cells depleted by positive sorting) was quantified via IFNγ ELISpot, revealing a significant increase in antigen-specific activation after SARS-CoV-2 vaccination (**Fig. 4f,g**). These results verified activation of CD4^+^ T cells, demonstrating an antigen-specific immune response critical for immune memory.

### DoriVac induces an antigen-specific CD8^+^ T cell activation

Furthermore, after two vaccine doses, we confirmed activation of cytotoxic CD8^+^ T cells. DoriVac increased the population of IFNγ secreting cytotoxic CD8^+^ T cells (**Fig. 5a**) and degranulating CD107a^+^ CD8^+^ T cells (**Fig. 5b**) in the LNs. PD-1 and CD69 were upregulated in the CD8^+^ T cell population (**Fig. 5c,d**), indicating increased activation. On Day 35, after the second dose, we quantified antigen-specific CD8^+^ enriched T cells (CD4^+^ T cell depleted by positive sorting) in the spleen via IFNγ ELISpot, showing increased antigen-specific CD8^+^ T cells after SARS-CoV-2 DoriVac (**Fig. 5e,f**). Co-delivery of SQB, free HIV-HR2 peptide, and free CpG did not induce similar level of T cell activation as observed with the delivery of DoriVac (**Supplementary Fig. 10**).

**Fig. 5.**
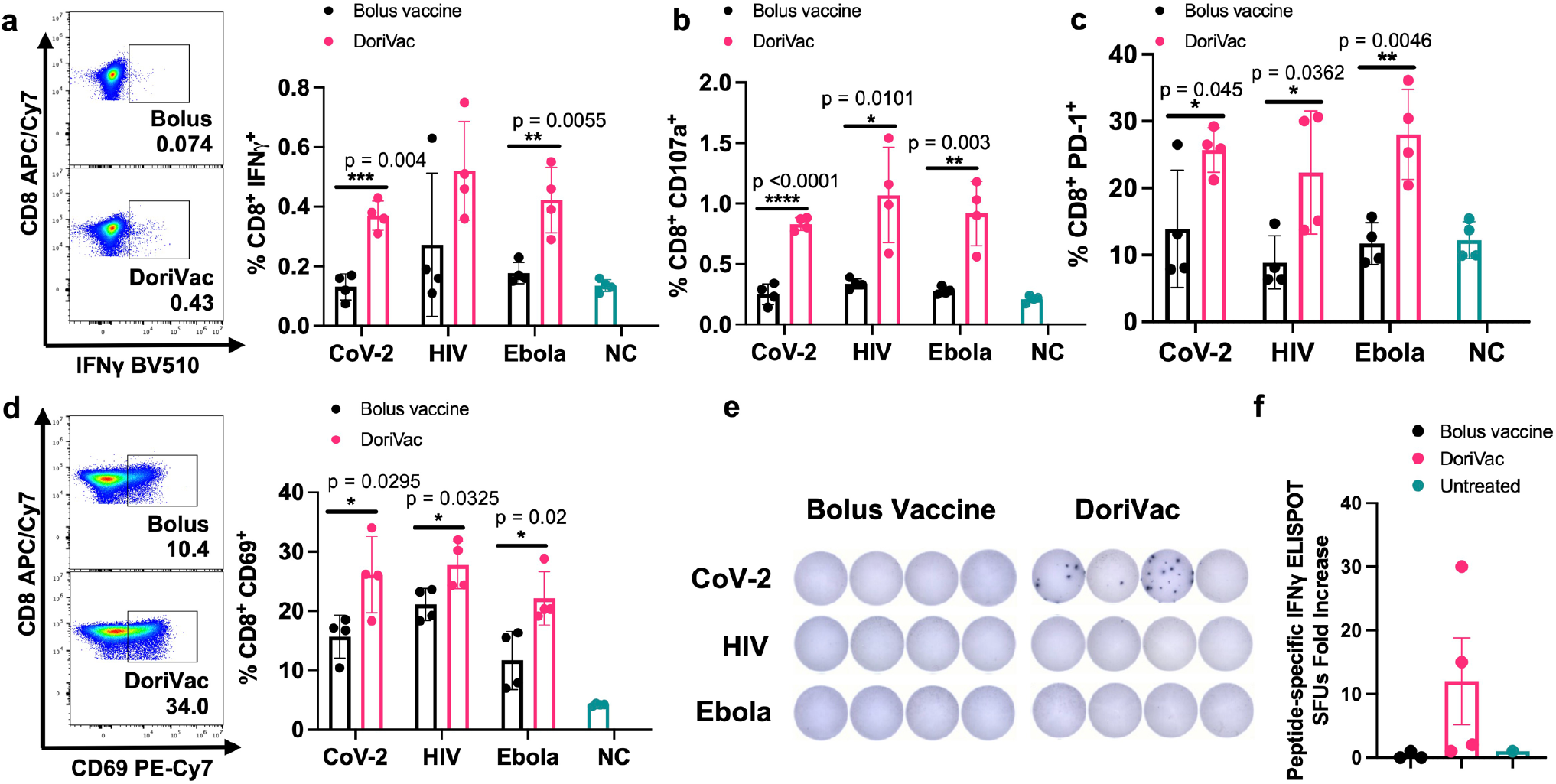
DoriVac induces enhanced antigen-specific CD8^+^ T cell activation in mice compared to bolus vaccine. **a**, Percentages of CD8^+^ IFNγ^+^ T cells in the lymph node (LN; n=4) on Day 21 as determined by flow cytometry and representative flow plots from the SARS-CoV-2 data. NC means negative (i.e. untreated) control. **b**, Percentages of CD8^+^ CD107a^+^ T cells in the LN (n=4) on Day 21 as determined by flow cytometry. **c**, Percentages of CD8^+^ PD-1^+^ T cells in the LN (n=4) on Day 21 as determined by flow cytometry. **d**, Percentages of CD8^+^ CD69^+^ T cells in the LN (n=4) on Day 21 as determined by flow cytometry and representative flow plots. **e–f**, IFNγ ELISpot demonstrating frequency of CD8^+^ enriched antigen-specific splenocytes (n=4, day 35) and accompanying quantification of IFNγ ELISpot SFUs demonstrates an increase significant difference in SARS-COV-2 antigen-specific CD8^+^ T cell frequency after treatment with DoriVac compared with the bolus vaccine. Data are represented as mean ± SD. The flow data were analyzed by multiple unpaired t-tests and significance was defined as a two-tailed p value less than 0.05. The ELISpot data were analyzed by one-way ANOVA (with correction for multiple comparisons using Tukey’s test). Statistical significance was defined as a multiplicity-adjusted p value less than 0.05. ‘*’ refers to p ≤ 0.05; ‘**’ refers to p ≤ 0.01; ‘***’ refers to p≤ 0.001; ‘****’ refers to p ≤ 0.0001; Bars without asterisks indicate no statistically significant difference in tested conditions (p > 0.05).

### Human immune-cell validation of peptide-conjugated and protein-conjugated DoriVac

Beyond murine models, we assessed DoriVac immunogenicity using a human LN organ-on-a-chip model. This model mimics the human LN for rapid prediction of vaccine responses in humans^61^. Analyzing the impact of SARS-CoV-2-HR2 DoriVac on human monocyte-derived DCs, we observed increased CD86, CD40, HLA-DR, and CD83 expression, indicating DoriVac can activate human DCs (**Fig. 6a, Supplementary Table 11, Supplementary Fig. 11**). DoriVac treatment also elevated inflammatory cytokines secreted by DCs compared to the bolus (**Fig. 6b**). Analyzing effector T cell responses on the human LN organ-on-a-chip model after nine days of vaccination, DoriVac displayed a substantial increase in CD4^+^ and CD8^+^ T cell activation in two of the three donors, as evidenced by TNFα^+^ and IL-2^+^ staining (**Fig. 6c, Supplementary Table 12, Supplementary Fig. 12**) Polyfunctionality analysis (examining T cells that express IFNγ, TNFα and IL-2) revealed significantly more CD4^+^ polyfunctional cells induced by DoriVac than the bolus vaccine, and an overall increase CD8^+^ polyfunctional cells (**Fig. 6d**). Inflammatory cytokine analysis indicated similar levels induced by bolus and DoriVac across three donors (**Fig. 6e**). Furthermore, using a Meso Scale Discovery (MSD) assay to measure antibody production on the human LN organ-on-a-chip, we observed robust and broad antibody responses against SARS-CoV-2 spike variants in samples treated with HR2 peptide-conjugated DoriVac (**Fig. 6f, Supplementary Fig. 13**). These findings suggest that DoriVac induces a robust immune response in an *in vitro* human immune system that closely predicts human vaccine response.

**Fig. 6.**
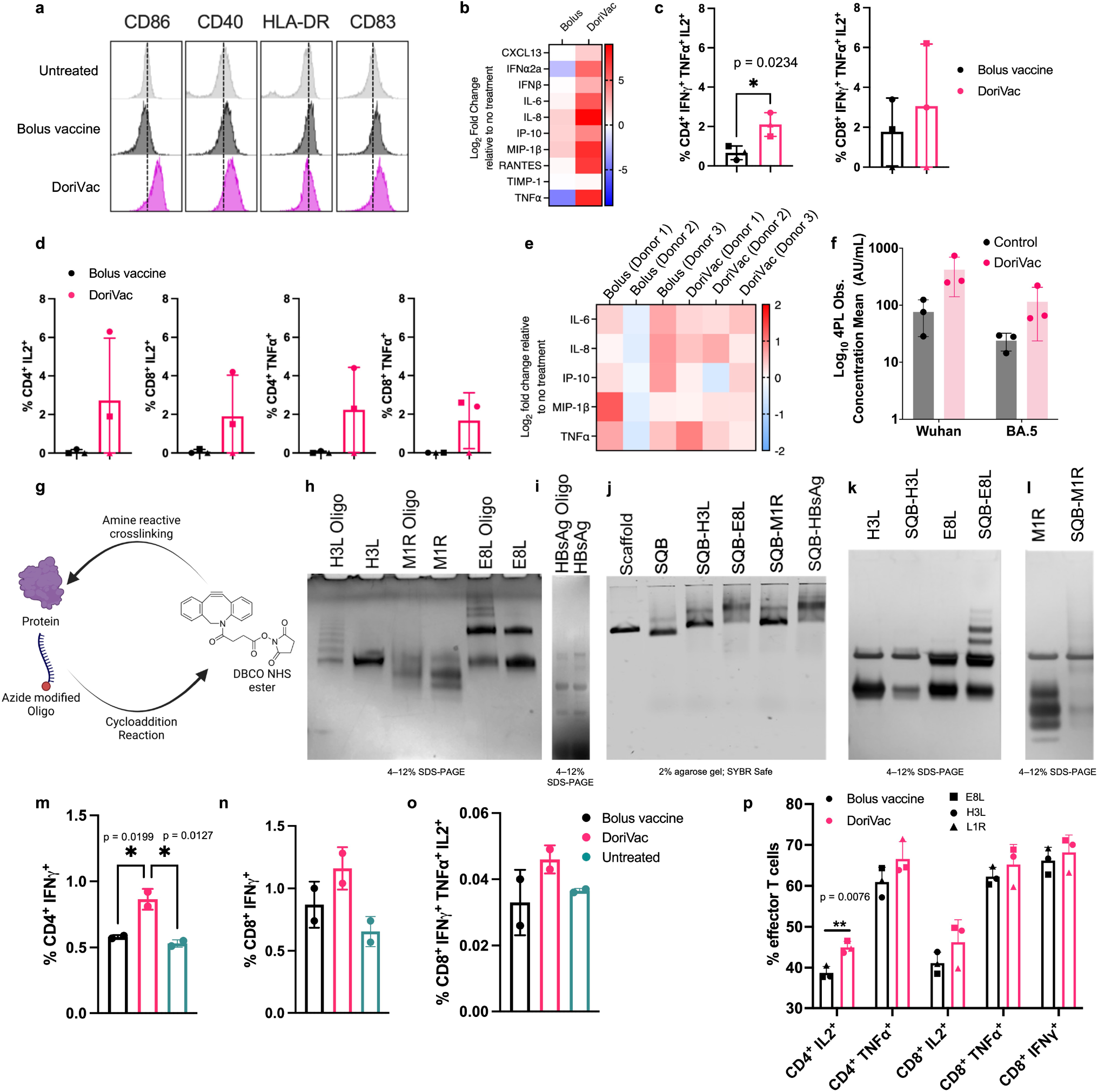
Peptide or protein-conjugated DoriVac effectively stimulates human DCs and induce enhanced immunogenicity compared to bolus vaccine on lymph node organ-on-a-chip model. **a**, Human monocyte-derived DCs were stimulated for 24 hours with bolus or DoriVac, and the DC activation markers were analyzed using flow cytometry. **b**, Relative fold changes in cytokines and chemokines after 24 hours of human monocyte-derived DCs with bolus or DoriVac. Fold change relative to no treatment is shown. **c**, Graph quantifying polyfunctional T cells, as determined by their ability to co-secrete IFNγ, TNFα and IL-2 as determined via intracellular cytokine staining and flow cytometry. **d**, LN organ-on-a-chips (n = 3) were vaccinated with bolus or the SARS-CoV-2-HR2 DoriVac. Nine days after vaccination, T cell responses were assessed via intracellular cytokine staining and flow cytometry after *ex vivo* stimulation with SARS-CoV-2 HR2 peptide and PMA/Ionomycin. Graph quantifying the average cytokine-producing CD4^+^ and CD8^+^ T cell populations at 9 days after vaccination in three different donors. Each symbol represents one donor. **e**, Relative fold changes in IL-6, IL-8, IP-10, MIP-1β and TNFα at 9 days after transduction either of the bolus or DoriVac on the LN organ-on-a-chip. Fold change relative to no treatment is shown. **f**, In a human LN organ-on-a-chip Meso Scale Discovery (MSD) assay, HR2 peptide-conjugated DoriVac induced a robust and broad antibody response against SARS-CoV-2 spike variants (n=3) compared to untreated control. Other tested variants are shown in Supplementary Fig. 13. **g**, Schematic representation of protein-oligonucleotide conjugation, demonstrating the utilization of DBCO-NHS ester crosslinker to attach an azide-modified oligonucleotide to the protein via free amine groups on lysines. **h**, SDS-PAGE gel (Coomassie-stained) confirming successful conjugation of monkeypox-specific proteins to oligonucleotides. Multiple bands in H3L-Oligo and E8L conjugates indicate variable degrees of modification; M1R shifts slightly relative to the unconjugated protein. **i**, SDS-PAGE gel demonstrating the successful conjugation of Hepatitis B surface antigen (HBsAg) proteins to oligonucleotides. **j**, Agarose gel verifying attachment of these oligo–protein complexes to the SQB DNA origami via handle–anti-handle hybridization. Conjugated SQB bands migrate more slowly than unconjugated SQB controls. **k–l**, Confirmation via SDS-PAGE gel of successful protein conjugation after DNase degradation of the DNA origami scaffold and staple strands and analysis of the remaining protein via silver stain. **m–o**, LN chips (n = 2) were vaccinated with bolus or DoriVac harboring CpG and full-length HBsAg. The T cell responses were assessed using intracellular cytokine staining and flow cytometry after *ex vivo* stimulation with autologous DCs pulsed with HBsAg (1:10 effector:target ratio), nine days after vaccination. Graphs quantify (m) IFNγ^+^ CD4^+^ populations, (n) IFNγ^+^ CD8^+^ populations, and (o) CD8^+^ polyfunctionality (IFNγ^+^, TNFα^+^, IL-2^+^) in two different donors. **p**, Tonsil organoids from one donor were vaccinated with bolus or DoriVac harboring CpG and full length monkeypox antigens (E8L, H3L or M1R). T cell responses were assessed using intracellular cytokine staining and flow cytometry after *ex vivo* stimulation with 15-mer monkeypox antigenic peptides (overlapping by 11-mer) and PMA/Ionomycin stimulation during the last 4 hours of incubation. Graph quantifies the cytokine-producing effector CD4^+^ and CD8^+^ populations, nine days after vaccination (n = 3 cell samples obtained from one preparation, treated with DoriVac fabricated with three different monkeypox antigens). The flow data for (c) and (d) were analyzed by unpaired t-tests (two-tailed p < 0.05), for (m–o) by one-way ANOVA with Tukey’s test (multiplicity-adjusted p < 0.05), and for (p) by multiple unpaired t-tests (p < 0.05). In all cases, ‘*’ refers to p ≤ 0.05; ‘**’ refers to p ≤ 0.01; Bars without asterisks indicate no statistically significant difference (p > 0.05). Data are represented as mean ± SD.

We further expanded our investigation to showcase the versatility of DoriVac, enabling the conjugation of full-length viral protein antigens. Protein vaccines historically faced limitations in presenting antigens on MHC-I and inducing CD8^+^ T cell responses^62^. As a proof-of-concept, we selected hepatitis B surface antigen (HBsAg) and three monkeypox antigens (E8L, H3L and M1R), validated for prior immunogenicity in their respective diseases (**Supplementary Table 13**)^63,64^.

The protein-’anti-handle’-oligonucleotide conjugate was synthesized using DBCO-azide click chemistry, where azide-modified oligonucleotide was conjugated to a protein via a DBCO-NHS ester linker (**Fig. 6g**). Successful conjugation was confirmed by SDS-PAGE gel (**Fig. 6h**,**i**). The protein-’anti-handle’-oligonucleotide conjugate was hybridized to the corresponding ‘handle’ strands on the SQB via Watson-Crick base-pairing. Protein-conjugated DoriVac exhibited reduced mobility in agarose gel electrophoresis compared to unconjugated DNA origami (**Fig. 6j**). Successful protein conjugation was observed via SDS-PAGE gel analysis after DNase I digestion (relative to a protein-only control) (**Fig. 6k**,**l**).

Initially, we assessed T cell responses to HbsAg-conjugated DoriVac and bolus vaccine in the human LN organ-on-a-chip system nine days post-vaccination. DoriVac-stimulated DCs, pulsed with HBsAg, induced IFN-γ secretion in both CD4^+^ and CD8^+^ subsets, demonstrating antigen-specific T cell activation (**Fig. 6m-n, Supplementary Table 12, Supplementary Fig. 12**). Polyfunctionality of T cells, measured by CD8^+^ T cells secreting IFNγ, TNFα and IL-2, was higher for DoriVac than the bolus (**Fig. 6o**). To confirm successful T cell responses to DoriVac, we evaluated the immunogenicity of monkeypox antigens in a tonsil organoid model^65^ from an independent donor, observing increased CD4^+^ IL-2^+^ and TNFα^+^ effector T cells, as well as a higher percentage of IL-2^+^, TNFα^+^ and IFNγ^+^ CD8^+^ T cells (**Fig. 6p**). These findings illustrate the capability of DoriVac to induce cellular immunity in humans against protein antigens of various infectious diseases.

### DoriVac and mRNA-LNP vaccines elicited comparable immune responses in mice

To assess the DoriVac vaccine platform’s suitability as protein antigen carrier and compare to commercially available mRNA-lipid nanoparticle (mRNA-LNP) vaccines, we fabricated DoriVac conjugated with complete SARS-CoV-2 spike proteins and evaluated its immunogenicity alongside two bivalent mRNA–LNP vaccines (**Fig. 7a,b, Supplementary Fig. 14**). Specifically, we compared two physiologically relevant doses of DoriVac (20 pmol and 100 pmol, see **Supplementary Methods**) to the bivalent vaccines (original and Omicron BA.4/BA.5) of mRNA-1273.222^1,66^ and BNT162b2-Omi.BA.4/BA.5^3,67^ (0.1 μg and 1 μg), hereafter mRNA-1273 and BNT162b2, in C57BL/6 mice (**Fig. 7c**). After the first booster dose on day 21, DoriVac at 100 pmol prompted greater dendritic cell (DC) mobilization in peripheral blood mononuclear cells (PBMCs) than the mRNA vaccines for both tested doses (**Fig. 7d, Supplementary Fig. 15**), suggesting an early, potent induction of key antigen-presenting cells. Notably, on day 50 (one day post-second booster), similar DoriVac-induced enhancements in DC activation and CD11b^+^ cell mobilization were evident in the lymph nodes and comparable to those of mRNA-LNP treatment groups (**Supplementary Fig. 16**), reflecting sustained antigen-presenting cell responses. All three vaccines elicited CD4^+^ and CD8^+^ effector and IFNγ-secreting T cells at similar levels in both PBMC (**Supplementary Fig. 17**) and lymphatic cell (**Fig. 7e,f, Supplementary Fig. 18**) populations. Additionally, DoriVac-treated groups showed elevated CD69 expression on T cells in the lymph nodes on day 50, at levels comparable to the mRNA-LNP treatment groups we tested (**Supplementary Fig. 18**). This observation suggested sustained activation within these primary immune sites, reinforcing that DoriVac’s immune stimulation persisted after each booster. We also noted heightened NK cell activation and increased IFNγ expression for DoriVac treatment groups at both dosage levels on day 50, underscoring the breadth of DoriVac’s immune engagement across multiple effector cell types (**Supplementary Fig. 19**). In addition, DoriVac-treated splenocytes and PBMCs produced high IFNγ levels comparable to those of mRNA-LNP vaccines at different time points upon *ex vivo* restimulation with SARS-CoV-2 peptides (**Fig. 7g, Supplementary Fig. 20**), underscoring a robust, antigen-specific T cell response across multiple tissue compartments. The specificity of these antibody responses were further validated against SARS-CoV-2 S1 domains (**Supplementary Fig. 21**) and spike proteins via ultra-sensitive single molecule array (SiMoA)^68^ (**Fig. 7h, Supplementary Fig. 21**) and pseudovirus neutralization assays (**Fig. 7i, Supplementary Fig. 22**). Notably, DoriVac elicited earlier and, at times, stronger antibody responses than the mRNA–LNP groups by day 22, highlighting a potentially faster humoral response under these dosing conditions. Interestingly, while demonstrating comparable or higher immune activation, DoriVac did not induce the marked PD-1 upregulation observed in mice treated with 1 μg BNT162b2 (**Supplementary Fig. 23**), suggesting that DoriVac may induce reduced T cell exhaustion compared to BNT162b2 under our tested conditions. Finally, we assessed intramuscular administration and found similarly robust T cell responses with DoriVac, indicating route flexibility (**Supplementary Fig. 24**). Comparable to mRNA–LNP vaccines, DoriVac drove strong PBMC IFNγ responses up to 21 weeks post-booster, emphasizing the sustained immunity generated by DoriVac (**Supplementary Fig. 24**). Strikingly, the day 91 (10 weeks) and day 147 (21 weeks) quantification of IFNγ-secreting cells from intramuscular administration showed intergroup comparisons comparable to the results from subcutaneous administration (**Fig. 7g, 7j, Supplementary Figs. 20, 24**), underscoring DoriVac’s robust and reproducible response over time. Despite fewer boosters and lower doses (10 pmol and 50 pmol, respectively), DoriVac consistently maintained IFNγ responses across extended time points, reinforcing its durable immunogenic profile. Consistent with reports of waning immunity in mRNA vaccines^69-73^, the BNT162b2 1 μg vaccine treatment group saw a marked reduction in IFNγ^+^ PBMC between day 91 and day 147, whereas the DoriVac treatment groups maintained fairly stable responses through the same interval. Moreover, to evaluate potential autoimmune or off-target responses, we measured anti-dsDNA and anti-PEG IgG two weeks and ten weeks after the second booster (day 35 and day 91), observing no significant increases compared to controls and thereby confirming DoriVac’s favorable safety profile (**Supplementary Fig. 25**). Collectively, these findings reinforce DoriVac’s promise as an effective DNA–protein vaccine platform capable of matching—and in certain measures surpassing—mRNA–LNP responses under our tested doses.

**Fig. 7.**
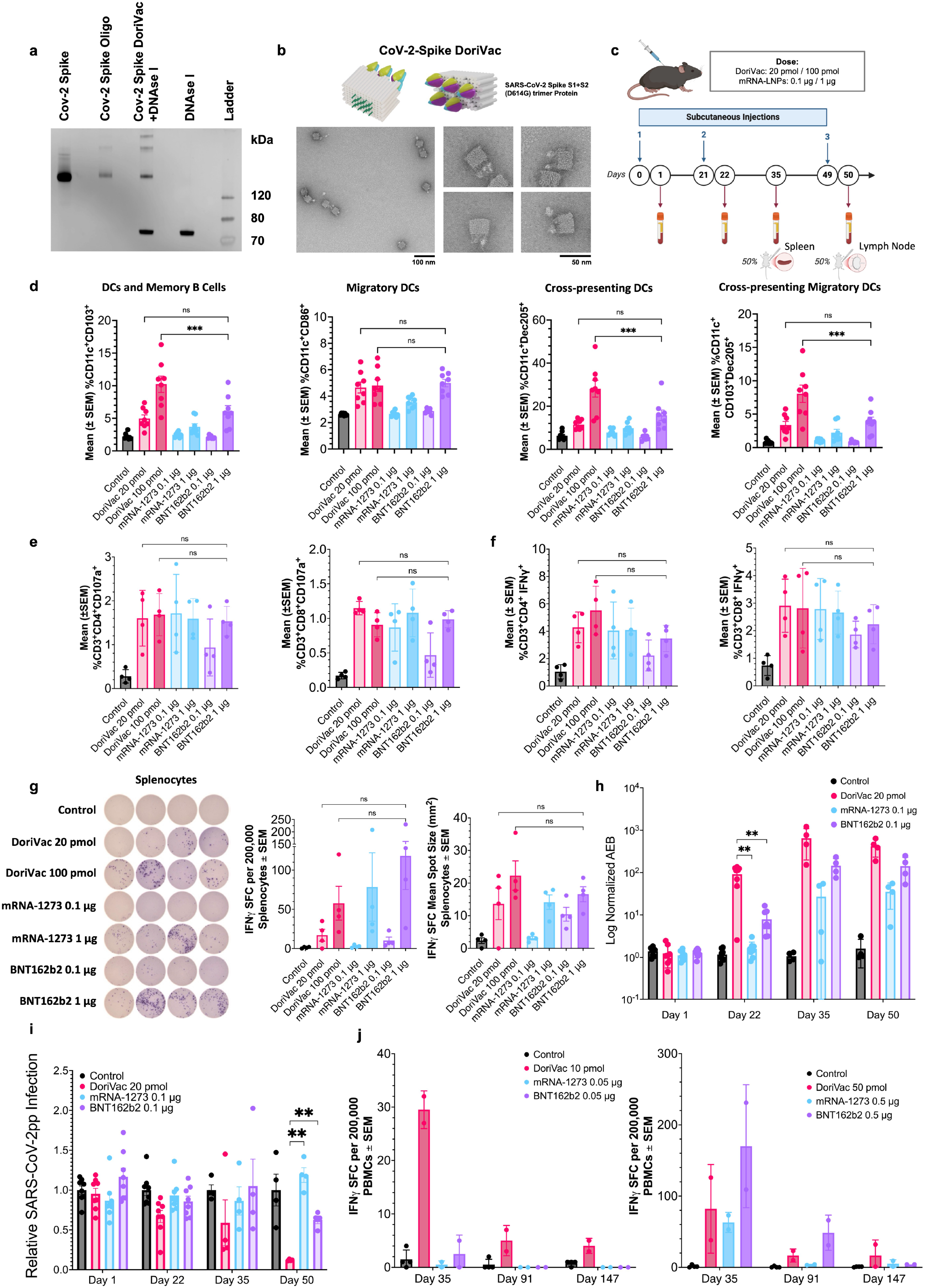
SARS-CoV-2 spike-protein-conjugated DoriVac induces potent cellular and humoral immune activation in mice. **a**, SDS-PAGE gel verifying successful conjugation of SARS-CoV-2 spike proteins to oligonucleotides and the successful hybridization of spike-protein-conjugated oligonucleotides on the DoriVac SQB DNA origami. **b**, Schematic representation of SARS-CoV-2 protein-conjugated DoriVac SQB and its negative-stained TEM images. **c**, Schematic delineating the vaccine administration protocol for naïve C57BL/6 mice and the data collection timeline. Blood samples were collected on day 1, day 22, day 35 and day 50. Plasma was separated for anti-SARS-CoV-2 antibody quantification and pseudovirus neutralization assays, while PBMCs were isolated for flow cytometry and ELISpot assay. Spleens and lymph nodes (LNs) were collected on day 35 and day 50 respectively, after sacrificing the mice. **d**, Percentages of CD103^+^, CD86^+^, Dec205^+^, and CD103^+^ Dec205^+^ double-positive CD11c^+^ PBMCs (n=8) on day 22 as determined by flow cytometry. Representative flow scatter plots are shown in Supplementary Fig. 15. DoriVac induced enhanced immune activation compared to controls, in some cases surpassing that of mRNA-LNPs. The flow data were analyzed by one-way ANOVA. **e–f**, Percentages of CD107a^+^ LN cells (n=4) and IFNγ-secreting cells (n=4) on day 50 as determined by flow cytometry. Representative flow scatter plots are shown in Supplementary Fig. 18. The DoriVac treatment group showed enhanced activation of these cells compared to the control and displayed comparative performance versus mRNA-LNP treatment groups. The flow data were analyzed by one-way ANOVA. **g**, IFNγ ELISpot demonstrating frequency of antigen-specific splenocytes (left, n=4, day 35), with accompanying quantification of IFNγ ELISpot SFUs and mean spot sizes. The results demonstrate an increase in SARS-COV-2 antigen-specific T cell frequency after treatment with DoriVac compared to the control, with larger spots indicating robust cytokine release. **h**, Quantification of relative anti-spike IgG levels in plasma samples of control and DoriVac 100 pmol/mRNA-LNP 1 μg dose treatment groups using SiMoA (n=8 on day 1 and day 22, n=4 on day 35 and day 50). DoriVac treatment group exhibited increased antibody levels compared to the control across all sampling dates. On day 22, DoriVac also showed a significantly higher antibody level than mRNA-LNP treatment groups. Data for spike-specific DoriVac 100 pmol/mRNA-LNP 1 μg dose responses and S1-specific responses for both dosage levels can be viewed in Supplementary Fig. 21. **i**, SARS-CoV-2 pseudovirus (SARS-CoV-2pp) neutralization assay (n=8 on day 1 and day 22, n=4 on day 35 and day 50; 1:20 dilution) for DoriVac 20 pmol/mRNA-LNP 0.1 μg dose treatment groups in model cell line ACE2-293T. DoriVac showed stronger neutralizing capabilities from Day 22 onward, compared to controls and, by Day 50, to the tested mRNA-LNP groups. Additional data for other dilutions and dosages are available in Supplementary Fig. 22. **j**, Longitudinal IFNγ ELISpot of PBMCs following intramuscular administration reveals the lasting T cell responses in DoriVac-treated mice, extending to Day 147, comparable to or exceeding those from mRNA–LNP groups (control n=4, others n=2). The data were analyzed by one-way ANOVA (with correction for multiple comparisons using a Tukey’s test) and significance was defined as a multiplicity-adjusted p value less than 0.05. ‘ns’ refers to p > 0.05; ‘**’ refers to p ≤ 0.01; ‘***’ refers to p ≤ 0.001.

## Discussion

Enduring immune memory and protection against viral variants rely on cellular immune responses, particularly CD8 T cells that target less mutable viral proteins. SARS-CoV-2 vaccines with greater than 90% protection demonstrated induction of Th1-skewed immunity^74-79^, emphasizing the importance of Th1 CD4 and CD8 T cell responses. In recent decades, DNA origami has achieved crucial milestones, indicating its potential as a modular therapeutic nanoparticle. Its programmability makes it a versatile ‘plug-and-play’ vaccine nanotechnology, particularly relevant for emerging infectious diseases. In this proof-of-concept study, HR2-DoriVac, conjugated with infectious-disease-associated peptides and proteins, elicited robust neutralizing antibodies and antigen-specific CD4 and CD8 T cell activation in healthy mice, a notable contrast to some SARS-CoV-2 vaccines with limited T cell responses. DoriVac presents four distinct advantages: (1) precise nanoscale arrangement of antigen and adjuvant for their co-delivery on each nanoparticle; (2) well-established, simple, and scalable fabrication; (3) modular, programmable design adaptable for various antigens via DNA hybridization; (4) stability at 4°C, contrasting with mRNA vaccines needing −20°C to −80°C cold chain storage^48^. Although we have not conducted a detailed cost analysis, DNA origami nanoparticles can potentially be produced at scale using established industrial processes, which may reduce manufacturing complexity and associated costs. Moreover, we have observed that DoriVac formulations remain stable at 4°C for at least several weeks^52^, offering logistical advantages over more stringent cold-chain requirements. Future studies focusing on process optimization, long-term storage conditions, and economies of scale will provide more quantitative data on cost-effectiveness and stability profiles, further informing DoriVac’s potential as a practical and accessible vaccine platform.

While robust antigen-specific T cell activation was observed with SARS-CoV-2-HR2 DoriVac, the same level of activation was not achieved with HIV and Ebola DoriVac formulations. Notably, we observed that HIV-HR2 and Ebola-HR2 peptide-functionalized DoriVac nanoparticles tended to form dimers and aggregates, as evidenced by agarose gel electrophoresis and TEM imaging. This aggregation is likely attributed to the greater hydrophobicity of these particular HR2 sequences, which can promote intermolecular interactions between peptide-conjugated nanoparticles. However, whether such aggregation will reduce vaccine efficacy and accessibility to immune cells needs further investigation. In future implementations, an interior hollow cavity in the origami could be engineered for hosting of peptides^80^, thereby mitigating aggregation. We have also explored the HIV and Ebola antigen epitopes prediction (**Supplementary Tables 7–10**), which showed fewer epitopes than CoV-2. While this may be the direct reason for the limited immune response, we acknowledge that binding affinity alone does not equate to actual peptide immunogenicity. Future *in vitro* and *in vivo* assays will be crucial to validate and refine these computational findings.

However, significant activation of B cells, DCs, CD4 and CD8 T cells for HIV and Ebola, suggested a strong immune response, albeit possibly insufficient for protection from infection. The chosen antigens for HIV and Ebola are also predicted to be weakly immunogenic in humans (**Supplementary Table 14–17**), so future studies could focus on identifying more immunogenic peptides for HIV and Ebola for presentation by human HLA alleles, potentially enhancing antigen-specific activation. Viral rechallenge studies, contingent on availability of BSL-3 facilities, could further assess the effectiveness of DoriVac-induced immune activation against live viruses. Additionally, while these results underscore DoriVac’s architectural versatility, they also highlight that not all antigens yield comparable levels of immune activation. The platform’s modular design ensures that the CpG-modified SQB scaffold and conjugation strategies can be easily adapted to different diseases; however, the ultimate efficacy depends on the intrinsic immunogenicity of the selected antigens. As a result, achieving consistently high efficiency across diverse pathogens may require antigen-specific optimizations and careful selection of epitopes to fully leverage DoriVac’s modular potential.

Our head-to-head comparison of SARS-CoV-2 spike–conjugated DoriVac and mRNA–LNP vaccines underscores DoriVac’s versatility and efficacy at the tested doses. By eliciting strong T cell and antibody responses with full-length spike protein, DoriVac demonstrates adaptability in transitioning from peptide-to protein-based antigens, matching or potentially surpassing mRNA–LNP responses. Moreover, the complete spike protein encompasses a broader array of epitopes than short peptide fragments, which can further enhance overall immunogenicity by engaging multiple T and B cell clones. Notably, we observed a significant induction of PD-1 expression in the higher dose BNT162b2-vaccinated group compared to those vaccinated with DoriVac, which may indicate a potential exhaustion of CD8 T cells in the higher dose BNT162b2 group. Further mechanistic studies are warranted to clarify how this induction impacts long-term immune memory and informs vaccine design. These findings are dose-dependent; however, only two doses were tested. A broader dose range and varied boosting intervals will be needed to identify optimal regimens and further assess DoriVac’s relative performance. Systematic dose-ranging, multiple boosting schedules, and evaluations against other protein-based vaccines will clarify where DoriVac excels and how it complements existing options. Comparisons under similar dosing with commercial protein-based vaccines could be used to evaluate DoriVac’s real-world competitiveness and its benefits, including sustained immune responses and simpler storage. DoriVac’s consistent performance via both subcutaneous and intramuscular routes highlights its flexibility, underscoring its potential to address mRNA–LNP limitations such as storage constraints. Future *in vivo* studies on long-term immunity and protective efficacy will further explore DoriVac’s potential as an alternative, particularly given its capacity to incorporate other full-length proteins and adapt quickly to emerging pathogens.

This proof-of-concept study highlights DoriVac’s capacity to rapidly generate vaccines against emerging infectious diseases and variants. Its programmable modularity supports multiplexed nanoparticles carrying diverse antigens, offering the promise of a versatile vaccine strategy. With a constant CpG-modified SQB scaffold, antigens can be readily exchanged, further supporting rapid adaptability across pathogens. This versatility positions DoriVac as a notable advancement for rapidly evolving infectious threats. Taken together, these attributes make DoriVac a promising next-generation platform that offers robust, sustained immune responses and simpler storage than current mRNA-based approaches.

## Methods

### SQB fabrication

SQB fabrication is detailed in a previous publication, including scaffold and staple sequences^52,81^. Scaffold p8634 was produced in-house, as previously published^82^. DNA staple strands were purchased from IDT. Folding concentrations were 5 mM Tris base, 1 mM ethylenediaminetetraacetic acid (EDTA; pH 8.0), 12 mM MgCl_2,_ 20–100 nM scaffold, 5 times excess of the basic staple strands (relative to scaffold concentration), 10 times excess of handle-conjugated staple strands (for attachment of relevant infectious-disease antigens) and 20 times excess CpG-staple strands. An 18-hour thermocycler program was used to fabricate SQBs: denaturation at 80°C for 15 minutes, then annealing via a temperature ramp from 50°C to 40°C decreasing at −0.1°C every 10 minutes and 48 seconds. Most staple strands include ten thymidine residues at the end of the double helices to minimize aggregation. The CpG-containing strands were appended on the flat face of the SQB. The CpG oligonucleotides with nuclease-resistant phosphorothioate backbones (5’
s – TCCATGACGTTCCTGACGTT-3’, IDT) replaced the corresponding thymine residues in a 3.5 nm nanoscale pattern as determined previously^52^. CpG was appended to the 5’ ends of designated strands.

### HR2 peptide conjugation with ‘anti-handle’ oligonucleotide

An ‘anti-handle’ oligonucleotide, which corresponds to 24 sites of ‘handle’ oligonucleotide on the extruding face of the SQB, was ordered from IDT with a 5’ amine (aminoC6-TTCTAGGGTTAAAAGGGGACG). HR2 peptides were ordered from GenScript with an azide-modified N-terminal lysine (**Fig. 1i–l; Supplementary Table 2**) for DBCO-Azide copper-free click chemistry reaction between the peptide and oligonucleotide. The oligonucleotide was prepared at 1 mM with phosphate buffer (pH 8.0). The dibenzocyclooctyne-N-hydroxysuccinimidyl ester (DBCO-NHS ester; Millipore, #761524) (diluted in DMSO to 2 mM) was incubated with the oligonucleotide (diluted in phosphate buffer pH 8.0 to 100 uM) in 1:1 volume ratio and 1: 20 oligonucleotides to DBCO ratio (with final concentration of DBCO >1 mM) and incubated overnight at ambient temperature. The oligonucleotide-DBCO was purified via Illustra NAP column (GE Healthcare Life Sciences, #17-0852-02), eluted with sterile water, and concentrated via 3K Amicon Ultra Centrifugal Filter Unit (Millipore; #UFC500324). DBCO incorporation was confirmed via OD310 peak. Concentration measured via A260 with the NanoDrop. The Azide-modified HR2 peptides, representing SARS-CoV-2, HIV and Ebola, were dissolved in DMSO to 5 mM. The peptide-Azide was mixed with the oligonucleotide-DBCO at a ratio of 1.5: 1 and incubated overnight at room temperature in 1 × PBS.

### Denaturing PAGE (dPAGE) verification and purification of peptide-conjugated oligonucleotide

15% denaturing PAGE (dPAGE) was used to confirm peptide-oligonucleotide conjugation. dPAGE gel (15%) was prepared using 9 mL urea concentrate (Fisher Scientific #EC-833), 4.5 mL urea dilutant, 1.5 mL urea buffer, 10 μL tetramethylethylenediamine (TEMED) and 150 μL 10 wt% ammonium persulfate in a casette (ThermoFisher Novex #NC2010)^52^. 5 picomoles of the sample were mixed in a 1:1 ratio with formamide loading buffer (FLB) and denatured at 80°C for 10 minutes before loading into wells. Electrophoresis was carried out in 0.5 × TBE buffer at 250V for 45 minutes, followed by staining with SYBR Gold and imaging using the Typhoon Gel Scanner. For purification, the mixture was combined with formamide loading buffer (FLB), loaded into an 8% dPAGE gel in a large well formed using a taped comb, and run at 250V for 50 minutes. The peptide-oligonucleotide was observed through UV shadowing on a thin layer chromatography plate, excised, crushed, and immersed in 1× TE buffer. After overnight shaking at 25°C, purification was performed using Freeze ‘N Squeeze DNA Gel Extraction Spin Columns (Bio-Rad; #7326165) and ethanol precipitation. The resulting peptide-oligonucleotide was resuspended in 1× PBS, and its concentration was determined using NanoDrop. Confirmation of purification was achieved via dPAGE.

### Protein conjugation with ‘anti-handle’ oligonucleotide

We generated protein-oligonucleotides by conjugating azide-modified DNA handles to the protein (obtained from Sino Biologic or Advanced ImmunoChemical, **Supplementary Table 13**) via the amino group present on lysine residues using DBCO-Azide click chemistry. The reaction was incubated in phosphate buffer with a pH of 8.0 overnight 4°C and was subsequently agitated at 37°C for 30 minutes. At higher pH levels (> 8.0), NHS reactivity towards the ε-amino groups of lysines is enhanced compared to the α-amino group^83^. For gel quantification assays, the protein was simultaneously labeled with NHS-Cy3 dye. DTT was also added to the reaction to reduce any disulfide bonds.

For SARS-CoV-2 spike protein conjugations, the purified spike protein (Sino Biological, Cat. #40589-V08H8, in phosphate buffer pH 8.0) was conjugated to azide-modified oligonucleotides similar to above but agitated longer at 37°C for 60 minutes. Conjugation efficiency was assessed by SDS-PAGE and silver staining, and successful SQB attachment was confirmed via agarose gel electrophoresis and TEM.

### Silver stain verification and purification of protein-conjugated oligonucleotide

Silver stain confirmed protein-oligonucleotide conjugation. The sample, mixed with 4× NuPAGE LDS sample buffer (ThermoFisher; #NP0008), underwent incubation at 95°C for 2 minutes before loading onto 4– 12% NuPAGE Bis-Tris gels (ThermoFisher; #NP0322) and electrophoresis at 150V for 45 minutes in 1× MES SDS running buffer (ThermoFisher; #NP0002). Gel analysis followed Pierce’s (#24612) silver staining protocol using Image Lab 6 on a Gel Doc EZ Imager (Bio-Rad). Purification utilized a 10K Amicon filter; the reaction sample, supplemented with phosphate buffer, was centrifuged at 14000rcf for 30 minutes until the flow-through reached a DNA concentration of less than 1ng/μl, indicating removal of all unconjugated DNA. Buffer exchange was carried out to 1xTE 10mM MgCl_2_.

### Peptide-or protein-conjugated oligonucleotide hybridization with SQB

The peptide-oligonucleotides or protein-oligonucleotides were hybridized to the SQB DNA origami in a 2× excess, maintaining 10 mM MgCl2 and 1× TE by adding stock 10× TE and 100 mM MgCl2. SQBs were added last to ensure a consistent buffer environment. The resulting conjugated SQBs were incubated at 37°C for 1–2 hours with shaking, followed by purification through PEG precipitation. Analysis was performed using agarose gel electrophoresis, TEM, and a DNase I degradation assay.

### Agarose gel electrophoresis

SQBs were analyzed via 2% native agarose gel electrophoresis. Gel was prepared with 0.5 × TBE buffer with 11 mM MgCl_2_ and 0.005% v/v SYBR Safe (ThermoFisher #S33102), run at 70V for 2 hours, and imaged via a Typhoon Gel Scanner.

### Transmission electron microscopy (TEM) analysis

Transmission electron microscopy (TEM) was utilized to assess structural integrity and SQB aggregation using negative-stain techniques. Formvar-coated, carbon-stabilized grids, either self-prepared or obtained from Electron Microscopy Services (FCF200-CU-TA), were plasma-discharged for 30 seconds for passivation. Subsequently, 4–10 nM SQBs were deposited on the grids for 45 seconds, followed by blotting with filter paper. Uranyl-formate solution (0.75% w/v in H2O) was applied to the grid, blotted off, and a second application lasted for 2 minutes before blotting. Imaging of the grids occurred using a JEOL JEM-1400 TEM in brightfield mode at 120 kV.

### SQB purification via PEG precipitation

CpG-SQBs or peptide-or protein-conjugated CpG-SQBs were purified via PEG precipitation. 1 × TE buffer (5 mM Tris base, pH 8.0 and 1 mM EDTA acid) containing 15% w/v PEG-8000 (Fisher Scientific, BP2331) and 510 mM NaCl was added to the SQB sample at 1:1 volume and mixed gently via pipetting. MgCl_2_ stock was added to the PEG solution to achieve 10 mM MgCl_2_ final concentration. As described previously, the solution was incubated for 30 min, centrifuged at 16000 g for 25 minutes and the supernatant was removed^52^. This procedure purifies and concentrates the sample, as a high concentration is often required for further studies. The concentration was determined via Nanodrop; the sample purity and integrity were confirmed via agarose gel electrophoresis and TEM.

### DNase I degradation and silver stain analysis of conjugation efficiency

One μg of SQBs was incubated with 1.0 U/μL DNase I (NEB) with 10 × DNase I buffer diluted in water (New England Biolabs #M0303S). Samples were incubated in the thermocycler for 30 minutes at 37°C. The silver stain was performed as described above. ImageJ was used to quantify band intensities and determine peptide or protein loading efficiencies.

### K10-PEG5k coating of SQBs

Before administration, PEG-purified peptide-or protein-conjugated SQBs were mixed with oligolysine-PEG5k (K10-PEG5k; methoxy-poly(ethylene glycol)-block-poly(L-lysine hydrochloride); n=113, x=10; Alamanda Polymers (mPEG20K-b-PLKC10) based on the calculated number of phosphates in the SQB sequence. An appropriate quantity of K10-PEG5k was added to match the number of nitrogens in its amines with the SQB phosphates, following a previously published method^84^. The mixture underwent incubation at 37°C for at least 30 minutes, and concentration was determined based on dilution.

### Animal model and treatment

C57BL/6 mice (6–8 weeks old) were obtained from Jackson Laboratory and housed at the Harvard Medical School (HMS) animal facility. Eight groups of eight mice each underwent the following treatments: (1) SARS-CoV-2-HR2 DoriVac, (2) SARS-CoV-2 HR2 bolus, (3) HIV-HR2 DoriVac, (4) HIV-HR2 bolus, (5) Ebola-HR2 DoriVac, (6) Ebola-HR2 bolus, (7) SQB-CpG + free HIV-HR2 peptide, and (8) untreated. The bolus contained an equivalent dose of CpG and HR2 peptide in PBS. Following one-week acclimation, mice received the first treatment dose (100μL of DoriVac containing 0.36 nmoles of CpG, 0.48 nmoles of HR2 peptide) subcutaneously on days 0 and 20. Blood was drawn via submandibular vein puncture four hours after dosing and also on days 14, 20, and 28. On days 21 and 35, four mice from each group were sacrificed; heart blood, LNs, spleens, and femurs were collected. All procedures were approved by the HMS Institutional Animal Care and Use Committee.

mRNA-LNP vaccines (mRNA-1273.222 and BNT162b2-Omi.BA.4/BA.5) were diluted in sterile DPBS to the indicated doses before administration. Subcutaneous or intramuscular (right hind leg; quadriceps femoris) injections were performed using fresh preparations to maintain vaccine integrity.

### Lymph-node-on-a-chip and tonsil organoid vaccination

Lymph-node-on-a-chip (LN chip) was fabricated as previously described^61^. Tonsil organoids was seeded according to a prior publication^65^. Human patient-derived apheresis collars and tonsil were obtained from the Crimson Biomaterials Collection Core Facility under approval of Harvard University’s Institutional Review Board. Chips and organoid culture were treated with 1nM of vaccine. In LN chip experiments, medium (RPMI supplemented with 10% FBS, 1% antibiotics, IL-2 and IL-4 as previously described^61^) circulated for 4 days of treatment (i.e., effluents were added back to the inlet perfusion reservoir), and at day 4, a 1:1 mix of effluent and fresh medium was used for perfusion to maintain the cytokine milieu. The outflow of the day 14 chip culture was collected to detect the antibody response against SARS-CoV-2 spike variants. In the tonsil organoid experiment, fresh media was added in a 1:1 ratio at day 4. At the study’s conclusion, cells were harvested by blocking one port of the basal channel and manually pipetting Cell Recovery Medium (Corning, 354253, 200 μL per chip) through the other port to extract the ECM and cells. The ECM was incubated in Cell Recovery Medium for 1 hour at 4 °C to depolymerize it and release associated cells. The released cells were centrifuged at 300 × g for 5 minutes and resuspended in PBS.

### Processing blood cells

Blood was collected either via heart extraction or through a submandibular cheek draw into heparin-coated tubes. Plasma and blood cells were separated via centrifugation at 800g and 4°C for 5 minutes. The collected plasma was stored at −80°C until analysis, while the blood cells underwent treatment with red blood cell lysis buffer (10×) from BioLegend (#420301) three times, following the manufacturer’s protocol. Peripheral blood mononuclear cells (PBMCs) were subsequently analysed using ELISpot and/or flow cytometry (Cytoflex LX).

### Luminex Multiplex ELISA analysis

The customized Bio-Plex Pro Mouse Cytokine Standard 23-plex kit from Bio-Rad included the following cytokines: IL-1α, IL-1β, IL-2, IL-3, IL-4, IL-5, IL-6, IL-9, IL-10, IL-12p40, IL-12p70, IL-13, IL-17A, Eotaxin, G-CSF, GM-CSF, IFNγ, MCP-1, MIP-1α, MIP-1β, RANTES, TNFα, following the manufacturer’s protocol. Data collection was performed using the Bio-Plex 3D Suspension Array System (Bio-Rad).

### Processing lymph nodes (LNs)

Following euthanasia, the upper axillary and superficial cervical LNs from the mouse were harvested and stored in cold PBS, as previously outlined^52^. These LNs were processed into single-cell suspensions for flow cytometry analysis (Cytoflex LX) by gently mashing them through a 40 μm cell strainer using a sterile syringe plunger into a petri dish. Cells were collected into 1.5 mL Eppendorf tubes, centrifuged at 400g for 5 minutes at 4°C, and the supernatant was discarded. The cell pellet was resuspended in 700 μL of PBS and distributed into 96-well plates for flow cytometry analysis.

### Processing spleens

Following euthanasia, mouse spleens were harvested, washed with PBS, and mashed through a 40 μm cell strainer into a 60 mm petri dish using a sterile syringe plunger. The resulting single-cell suspension was washed with complete RPMI-1640 media (containing 10% fetal bovine serum and 1% penicillin-streptomycin), collected into a Falcon tube, and subjected to two treatments with red blood cell lysis buffer (BioLegend; #420301) following the manufacturer’s protocol. The cell pellet was resuspended in 2 mL of complete RPMI media, and the cell count was determined. These splenocytes were used for flow cytometry or ELISpot assays.

### Processing bone marrow

Following euthanasia, femurs were repeatedly washed with PBS. Muscle fibers and connective tissues were extracted using forceps. Marrow extraction involved flushing the bone with a syringe into a PBS-filled dish. The collected marrow clot was pipetted, filtered through a 40 μm cell strainer, and gathered into a Falcon tube. The resulting single-cell suspension underwent centrifugation at 300 × g for five minutes. After discarding the supernatant, the pellet was treated with red blood cell lysis buffer (BioLegend #420301) following the manufacturer’s instructions. The suspension was centrifuged at 300 × g for five minutes and subsequently resuspended in culture media for flow cytometry (Cytoflex LX).

### Flow cytometry

Single cell suspensions of LNs, PBMCs, spleens, and bone marrow were obtained. The suspensions were washed with PBS, stained with Zombie UV (BioLegend; #423108) or ViaKrome 808 (Beckman Coulter #C36628) viability dye and washed with cell staining buffer (BioLegend, #420201). The cells were stained with fluorophore-conjugated cell surface antibodies (**Supplementary Table 4-6, 11-12, 18**). Intracellular staining was performed using permeabilization and fixation reagents (BioLegend; #424401). Antibodies were arranged into appropriate panels, compensations were set up to minimize fluorescent emission overlap, and the cells were analyzed on a Cytoflex LX flow cytometer. Storage events were gated on the population of interest, based on protocols published previously^52^, and according to the gating in **Supplementary Figures 2, 5, 9, 11, 12, 16, 18, and 19**. Flow data was analyzed using FlowJo V10.

### CD8 and CD4 enrichment of splenocytes

Splenocytes were depleted for CD4 or CD8 T cells using CD8 Dynabeads™ (Thermofisher, #11145D) or CD4 Microbeads (Miltenyi Biotec, #130-117-043), according to manufacturer’s instructions. The remaining sample was enriched for CD4 T cells (via CD8 T cell depletion) or enriched for CD8 T cells (via CD4 T cell depletion). Splenocytes were maintained in 4ºC for 36 hours before processing for CD8 T cell enrichment. For CD8 depletion, the Dynabeads™ were washed in isolation buffer and placed in the magnet. At the same time, cells were prepared at a concentration of 1 × 10^7^ cells per mL in isolation buffer. The prewashed beads were added, and the solution was incubated for 30 minutes at 4°C with gentle tilting. After, the tubes were placed in the magnet for 2 minutes and the supernatant was transferred to a new tube for further analysis. The beads and associated cells were discarded. Regarding CD4 depletion via Microbeads, the cells were incubated with the microbeads for 10 minutes at 4°C and then processed through an LD column, yielding the CD8-enriched (CD4-depleted) population in the flow-through for subsequent analysis.

### IFNγ ELISpot

Samples were processed into single-cell suspensions, followed by plating PBMCs or splenocytes into a 96-well round-bottom plate, each containing cells from an individual mouse in 200 μl of media. The cell quantities utilized were: two and a half million cells for splenocytes on day 21, three million cells for splenocytes on day 35, two million cells for PBMCs, eight and a half million cells for CD4 and CD8 enriched samples and two hundred thousand cells for full-length SARS-CoV-2 Spike DoriVac and mRNA-LNP assays. Each well was stimulated with 2 μg/mL of HR2 peptide or 1.67 μg/mL SARS-CoV-2 peptide pools (Miltenyl Biotec, #130-127-951, #130-132-051). For HR2 peptide experiments, after 48 hours of incubation, cells were collected, resuspended in 100 μl of sterile media, and plated onto an ELISpot plate (RND systems, Mouse IFNγ ELISpot kit, #505841) to incubate for 36 hours at 37°C. Cells from full-length SARS-CoV-2 Spike DoriVac and mRNA-LNP animal experiments were directly plated with stimulating peptides on an ELISpot plate and incubated for 20 hours at 37°C. The plate was then processed as per the manufacturer’s guidelines, and analyzed using an ELISpot plate reader at Dana Farber Cancer Institute’s Center for Immuno-oncology Translational Immunogenics Laboratory.

### Enzyme-linked Immunosorbent Assay (ELISA)

Plasma IgG from vaccinated mice was quantified using an ELISA method. Nunc Maxisorp ELISA plates (ThermoFisher, USA #44-2404-21) were coated with HR2 peptide at a concentration of 2–20 μg/mL in 100 μL of coating buffer (100 mM bicarbonate/carbonate buffer, pH 9.5) and incubated overnight at 4°C. After washing three times with washing buffer (PBS containing 0.05% Tween 20), 150 μL of blocking buffer (2% bovine serum albumin (Sigma, USA #9048-46-8) in washing buffer) was applied for 1 h at 37°C. After removing the blocking buffer, 100 μL of plasma samples diluted in blocking buffer (1:100, 1:200, 1:400 dilutions) were added and incubated for 1 h at 37°C. After washing three times with washing buffer, 150 μL of blocking buffer was applied for 1 h at 37°C. After removing the blocking buffer, 100 μL of HRP-conjugated anti-mouse IgG antibody (Cell Signaling Technology, USA #7076) diluted in blocking buffer was applied for 1 h at 37°C. After washing five times with washing buffer, 50 μL of 3,3’,5,5’-tetramethyl benzidine substrate (Sigma #54827-17-7) for detection was added, and the reaction was stopped after 15 min by the addition of 50 μL of 1 M H_2_SO_4_. Absorbance at 450 nm was measured using an automated plate reader (BioTek).

### Pseudovirus assay

Plasma was isolated by collecting the clear supernatant post-centrifugation. Samples were diluted in culture media at varying ratios and cultured with the corresponding pseudovirus and ACE2-293 T cells. Relative pseudovirus infection level was assessed as the ratio of infected cells in each group to those in the bolus or the control group, which was assigned a relative infection level of 1.0.

### Single molecule array (SiMoA)

Multiplexed single molecule array (SiMoA) assays were used to measure anti-SARS-CoV-2 immunoglobulin G (IgG) antibodies against the S1 subunit of the spike protein, the full spike protein, and nucleocapsid protein (control), as previously described^68^. The collected mouse serum samples were diluted 4000 fold and analyzed using an automated three-step assay format onboard an HD-X Analyzer (Quanterix). In the first step, the diluted serum samples are incubated with SARS-CoV-2 antigen conjugated fluorescent beads. Then the beads are washed and incubated in a solution of 1 ng/mL biotinylated anti-mouse IgG antibody (Abcam, ab6788) in the second step. The beads are washed again and incubated in a solution of 30 pM of streptavidin-β-galactosidase in the third step. Afterwards, the beads are resuspended in a solution of resorufin β-d-galactopyranoside and loaded into a microwell array for analysis. Average enzyme per bead (AEB) values were calculated by the HD-X analyzer and normalized between runs using a control solution of a neutralizing anti-spike mouse monoclonal antibody (Sino Biological, 40591-MM43) at a concentration of 5 pg/mL.

### Assay for anti-dsDNA and anti-PEG IgG

Anti-dsDNA and anti-PEG IgG levels were measured using the Mouse Anti-dsDNA IgG ELISA Kit (Cat. No. 5120, Alpha Diagnostic Intl.) and the Mouse Anti-PEG IgG ELISA Kit (Cat. No. PEG-030, Alpha Diagnostic Intl.), respectively. Both assays were performed according to the manufacturers’ protocols. Briefly, all kit-provided calibrators, samples, and positive controls were assayed in duplicate. Serum samples were diluted 1:100 in the supplied sample diluent, consistent with the recommended dilution of ≥1:100. After appropriate incubations and washes, the bound IgGs were detected with an anti-mouse IgG-HRP conjugate followed by TMB substrate development and measurement of optical density at 450 nm with normalization at 630 nm. Data analysis was carried out according to the manufacturers’ guidelines for each kit.

### SARS-CoV-2 Spike-Specific Immunoglobulin Quantitation in Lymph Node Chip Outflows

Anti-SARS-CoV-2 IgG concentrations were determined in day 14 lymph node chip outflows via the 10-PLEX Meso Scale Discovery (MSD) V-PLEX SARS-CoV-2 Spike Panel 27 (IgG) Kit (K15606U). Assays were performed according to the manufacturer’s specifications. Briefly, LN chip outflows were tested in duplicate at 1:10, 1:70, and 1:490 dilutions, with samples incubated on plates pre-coated with spike antigens from multiple SARS-CoV-2 variants (n=10); antibody binding was detected with SULFO-Tag-labeled anti-human IgG antibody. Data were acquired with an MSD Meso Sector S 600MM plate reader and analyzed with MSD Discovery Workbench 4.0 software. Sample antibody concentrations [arbitrary units per milliliter (AU/mL)] were calculated against the kit-provided standard (using four-parameter logistic regression), reporting data from the dilution nearest to the midpoint of the curve. Concentrations are reported for Wuhan and BA.5 spike variants in **Fig. 6f**, with data from additional variants reported in **Supplementary Figure 13**, including B.1.351 (beta variant), B.1.617.2; AY.4 Alt Seq 2 (delta variant), BA.2, BA.2.12.1, BA.2+L452M, BA.2+L452R, BA.3, and BA.4 (omicron sublineage variants). Three positive controls, containing known levels of SARS-CoV-2-specific IgG, were included in the kit and tested to confirm assay performance. CH65 IgG human monoclonal antibody^85^, specific to influenza receptor binding site hemagglutinin glycoprotein, served as a negative control.

### Statistical analyses

One-way or two-way ANOVA or unpaired t-test(s) with appropriate corrections for multiple comparisons as detailed in the figure captions was applied to determine the statistical significance of all flow, ELISpot, and ELISA data in **Figures 2–7**. GraphPad Prism 10 was used to make graphs, analyze statistics, and calculate p values. A p value ≤ 0.05 was considered statistically significant. ‘*’ refers to p ≤ 0.05; ‘**’ refers to p ≤ 0.01; ‘***’ refers to p≤ 0.001; ‘****’ refers to p ≤ 0.0001. Error bars represent standard deviation (SD).

### Reporting Summary

Further information on research design is available in the Nature Research Reporting Summary linked to this article.

## Data availability

Data supporting the findings of this study are presented in the paper and the supplementary materials. Source data for the figures will be provided with this paper.

## Acknowledgements

We appreciate the support and experimental input from Shih Lab members. Special thanks to Kathleen Mulligan, Maurice Perez, Michael Carr, Thomas Ferrante, and Eric Zigon for their expertise with instrumentation. We would also like to thank Jessica Baker Flechtner for her insights during results discussions and her assistance with figure preparation. **Figures 1b-d, Figure 2a, Figure 7c**, and **Supplementary Figure 24a** were generated using BioRender (https://biorender.com/).

The authors would like to thank the Harvard University Health Services for sharing the unused portion of mRNA-LNP vaccines (mRNA-1273.222 and BNT162b2-Omi.BA.4/BA.5) from opened vials to enable the comparative studies of DoriVac and mRNA-LNP vaccines.

## Author contributions

Y.C.Z. and L.S. developed the idea. Y.C.Z., O.J.Y., Q.X., L.S., and M.W.K. planned experiments. O.J.Y. and Q.X. fabricated and tested the vaccine. Y.C.Z., O.J.Y., and Q.X. led the animal studies and immune-cell profiling. L.S. led the HR2 design and neutralization studies with S.G.B., M.W.K., and A.R. fabricated the protein vaccine and tested via organ-on-a-chip with Y.Z., L.D.W., C.A.H, and G.D.T. quantified IgG against SARS-CoV-2 variants in lymph node organ-on-a-chip outflows using MSD. T.G. and Z.N.S. quantified relative anti-spike IgG levels in plasma samples using SiMoA. O.J.Y. and Q.X. drafted the manuscript with the guidance of Y.C.Z. and L.S. and supported by M.W.K. and A.R. in figure and writing preparation. Y.C.Z., O.J.Y., Q.X., L.S., M.W.K., S.G.B., H.D., G.I., T.G., Z.N.S., S.H.S., A.R., A.J., Y.Z., L.W., and C.A.H. performed experiments. O.J.Y. and Q.X. modeled the SQB and peptide/protein-conjugated SQB in Fusion with C.M.W. providing assistance. A.R.G., A.V., M.S., and S.B. assisted animal study design, model set-up, and sampling. J.H.R. and I.C.K. supported the project and manuscript editing. W.M.S., Y.C.Z., D.E.I., G.G., and G.D.T. provided overall guidance for the project, and edited the manuscript.

