## Supplementary Information for "DNA origami vaccine (DoriVac) nanoparticles improve both humoral and cellular immune responses to infectious diseases"

| Variant | Sequence |
|---|---|
| SARS-CoV-2 (Alpha) | GDISGINASVVNIQKEIDRLNEVAKNLNESLIDLQELG KYEQYIK |
| SARS-CoV-2 (Omicron) | GDISGINASVVNIQKEIDRLNEVAKNLNESLIDLQELG KYEQGSG |

**Supplementary Table 1 | Differences between HR2 sequences in variants of SARS-CoV-2.** The amino acids that differ between the two variant sequences are colored in red. HR2 peptide is highly conserved, compared to many epitopes within the spike region, which renders the effectiveness of HR2-based SARS-CoV-2 vaccines less susceptible to antigenic drift.

| Peptide Name | Sequence | Molecular Weight | Number of Residues | Net Charge at pH 7 | Hydrophobicity (GRAVY) |
|---|---|---|---|---|---|
| SARS-CoV-2 HR2 | [Lys(N3)]GDIS GINASVVNIQK EIDRLNEVAKN LNESLIDLQEL GKYEQYIK | 5203.81 g/mol | 46 | -2.0 | -0.52 |
| HIV HR2 | [Lys(N3)]WNN MTWMEWDRE INNYTSLIHSLI EESQNQQEKN EQELLELDKW ASLWNWF | 6600.19 g/mol | 52 | -5.9 | -1.14 |
| Ebola HR2 | [Lys(N3)]IEPHD WTKNITDKIDQ IIHDFVDK | 3049.38 g/mol | 25 | -1.8 | -0.98 |

**Supplementary Table 2 | Peptide sequences and relevant properties.** Peptides were purchased with N-terminal azide modifications from GenScript. The azide modifications enable a click-chemistry reaction between the azide-modified peptide and a DBCO-modified oligonucleotide. A negative value in the Grand AVerage of hydropathY (GRAVY) score indicates peptides that are net hydrophilic at pH 7.

| Sample | Intensity of DoriVac Band | Intensity of Peptide Band (Control) | Percent Conjugation Efficiency |
| --- | --- | --- | --- |
| SARS-CoV-2 DoriVac | 5885.468 | 5770.296 | 102.0% |
| Ebola DoriVac | 2660.154 | 2573.125 | 103.4% |
| HIV DoriVac | 3330.874 | 3436.288 | 96.9% |

**Supplementary Table 3 | Peptide conjugation efficiency.** Peptide conjugation efficiency was determined by using the band intensity of a DNase I-digested infectious disease SQB, as calculated by ImageJ, and comparing it with the band intensity of the theoretical amount of peptide.

| Antibody | Fluorophore | BioLegend Catalog Number |
|---|---|---|
| CD11c | BV510 | 117338 |
| CD40 | PE | 124610 |
| CD80 | APC/fire 750 | 104740 |
| CD86 | PE/Cy7 | 105014 |
| MHC-II | BV421 | 107632 |
| PD-L1 | BV785 | 124331 |
| DEC205 | PerCp/Cy 5.5 | 138207 |
| CD11b | BV605 | 101257 |
| CD103 | AF700 | 121442 |
| Gr-1 | FITC | 108406 |
| Viability | Zombie UV | 423108 |

**Supplementary Table 4 | Antibodies and fluorophores used for flow cytometric characterization of dendritic cell (DC) phenotypes in our DoriVac studies.** This table lists the antibodies employed to delineate DC lineage, activation status, and antigen-presentation capacity. Markers include DC lineage proteins (CD11c, CD11b, CD103, Gr-1), co-stimulatory molecules (CD40, CD80, CD86), antigen presentation molecules (MHC-II, DEC205), and the immune checkpoint regulator PD-L1, alongside the Zombie UV viability dye. These antibodies were assembled into panels according to specific experimental objectives to evaluate the DC responses associated with DoriVac immunogenicity.

| Antibody | Fluorophore | BioLegend Catalog Number |
|---|---|---|
| B220 | BV510 | 103247 |
| CD19 | AF700 | 115528 |
| CD138 | BV650 | 142518 |
| CD38 | PE/Fire 700 | 102747 |
| IgG1 | PerCP/Cyanine5.5 | 406612 |
| IgG2a | BV421 | 407117 |
| CD40 | PE/CY7 | 124622 |
| CD27 | APC/fire750 | 124237 |
| PD-L2 | PE594 | 107216 |
| Viability | Zombie UV | 423108 |

**Supplementary Table 5 | Antibodies and fluorophores used for flow cytometric analysis of B cell subsets in our DoriVac studies.** This table details the antibodies used to characterize B cell populations and monitor their activation and differentiation following DoriVac immunization. Markers such as B220 and CD19 identify the overall B cell compartment, while CD138 marks plasma cells. CD38 and CD27 help assess activation and memory status, whereas IgG1 and IgG2a detect class-switched immunoglobulins. Additionally, CD40 and PD-L2 indicate co-stimulatory functions. The Zombie UV viability dye is used to exclude dead cells. Antibodies were organized into panels tailored to specific experimental objectives, with each entry listing the corresponding BioLegend catalog number.

| Antibody | Fluorophore | BioLegend Catalog Number |
|---|---|---|
| CD3 | BV785 | 100232 |
| CD8 | APC/Cy7 | 100713 |
| CD4 | PE | 100408 |
| CD69 | PE-Cy7 | 104512 |
| CD62L | PerCp | 104430 |
| CD44 | AF-700 | 103026 |
| CD127 | APC | 135012 |
| CD25 | BV650 | 102037 |
| PD-1 | BV605 | 135220 |
| CD107a | FITC-AF488 | 121608 |
| Viability | Zombie UV | 423108 |
| FoxP3 (Intra) | BV421 | 126419 |
| TNFα (Intra) | PE594 | 506346 |
| IL-2 (Intra) | PE - Cy5 | 503824 |
| IFN-γ (Intra) | BV510 | 505841 |

**Supplementary Table 6 | Antibodies and fluorophores for multiparametric T cell profiling in DoriVac studies.** This table lists the antibodies used to characterize T cell subsets and assess activation, differentiation, and effector functions following DoriVac immunization. Markers such as CD3, CD4, and CD8 define the T cell compartment, while CD69, CD62L, and CD44 provide insight into activation and memory status. CD127, CD25, and PD-1 further refine the phenotypic profile, and CD107a indicates degranulation. Intracellular staining for FoxP3 identifies regulatory T cells, and TNFα, IL-2, and IFN-γ are measured to evaluate cytokine production. The Zombie UV viability dye is used to exclude dead cells. Antibodies were organized into specific panels according to experimental goals, and each entry includes the corresponding BioLegend catalog number.

| MHC-I or MHC-II | Allele | Start | End | Length | Peptide | Core peptide region | Rank |
|---|---|---|---|---|---|---|---|
| MHC-I | H-2-Db | 8 | 17 | 10 | ASVVNIQKEI | ASVVNIQEI | 0.04 |
| MHC-I | H-2-Db | 3 | 11 | 9 | ISGINASVV | ISGINASVV | 0.09 |
| MHC-I | H-2-Kb | 6 | 13 | 8 | INASVVNI | INAS-VVNI | 0.37 |
| MHC-I | H-2-Kb | 30 | 37 | 8 | SLIDLQEL | SLID-LQEL | 0.37 |
| MHC-I | H-2-Db | 8 | 20 | 13 | ASVVNIQKEIDRL | ASVVNIDRL | 0.39 |
| MHC-I | H-2-Db | 29 | 37 | 9 | ESLIDLQEL | ESLIDLQEL | 0.44 |
| MHC-I | H-2-Kb | 13 | 20 | 8 | IQKEIDRL | IQKE-IDRL | 0.99 |
| MHC-I | H-2-Db | 4 | 11 | 8 | SGINASVV | SGI-NASVV | 0.68 |
| MHC-II | H2-IAb | 2 | 16 | 15 | DISGINASVVNIQKE | INASVVNIQ | 2 |
| MHC-II | H2-IAb | 3 | 17 | 15 | ISGINASVVNIQKEI | INASVVNIQ | 2.4 |
| MHC-II | H2-IAb | 1 | 15 | 15 | GDISGINASVVNIQK | INASVVNIQ | 3.4 |
| MHC-II | H2-IAb | 4 | 18 | 15 | SGINASVVNIQKEID | INASVVNIQ | 4.4 |
| MHC-II | H2-IAb | 5 | 19 | 15 | GINASVVNIQKEIDR | INASVVNIQ | 18 |
| MHC-II | H2-IAb | 15 | 29 | 15 | KEIDRLNEVAKNLNE | DRLNEVAKN | 29 |
| MHC-II | H2-IAb | 14 | 28 | 15 | QKEIDRLNEVAKNLN | DRLNEVAKN | 36 |
| MHC-II | H2-IAb | 16 | 30 | 15 | EIDRLNEVAKNLNES | DRLNEVAKN | 40 |

**Supplementary Table 7 | Predicted murine MHC epitopes of the SARS-CoV-2 HR2 peptide and associated MHC binding.** NetMHCpan-4.1 was used to predict the top eight murine MHC-I and MHC-II epitopes of each HR2 peptide. The default cutoff for MHC binders was used as described by NetMHCpan[59]. A lower rank score for the peptide is associated with stronger binding to MHC. 0.5% was defined as the cutoff for 'strong' MHC-I binders while 2% was defined as the cutoff for 'weak' MHC-I binders. Any peptides that had a rank score >2% were assumed to not bind to MHC-I. 2% was defined as the cutoff for 'strong' MHC-II binders while 10% was defined as the cutoff for 'weak' MHC-II binders. Any peptides that had a rank score >10% were assumed to not bind to MHC-II.

| MHC-I or MHC-II | Allele | Start | End | Length | Peptide | Core peptide region | Rank |
|---|---|---|---|---|---|---|---|
| MHC-I | H-2-Db | 11 | 19 | 9 | REINNYTSL | REINNYTSL | 0.09 |
| MHC-I | H-2-Kb | 17 | 24 | 8 | TSLIHSLI | TSL-IHSLI | 0.42 |
| MHC-I | H-2-Kb | 16 | 23 | 8 | YTSLIHSL | YTS-LIHSL | 0.5 |
| MHC-I | H-2-Db | 30 | 38 | 9 | QQEKNEQEL | QQEKNEQEL | 0.39 |
| MHC-I | H-2-Db | 27 | 38 | 12 | SQNQQEKNEQEL | SQNQNEQEL | 0.43 |
| MHC-I | H-2-Kb | 12 | 19 | 8 | EINNYTSL | EINN-YTSL | 0.79 |
| MHC-I | H-2-Db | 11 | 20 | 10 | REINNYTSLI | REINNYTSI | 0.7 |
| MHC-I | H-2-Db | 29 | 38 | 10 | NQQEKNEQEL | NQQENEQEL | 0.85 |
| MHC-II | H2-IAb | 16 | 30 | 15 | YTSLIHSLIEESQNQ | LIHSLIEES | 11 |
| MHC-II | H2-IAb | 15 | 29 | 15 | NYTSLIHSLIEESQN | LIHSLIEES | 14 |
| MHC-II | H2-IAb | 14 | 28 | 15 | NNYTSLIHSLIEESQ | LIHSLIEES | 20 |
| MHC-II | H2-IAb | 17 | 31 | 15 | TSLIHSLIEESQNQQ | LIHSLIEES | 20 |
| MHC-II | H2-IAb | 12 | 26 | 15 | EINNYTSLIHSLIEE | NYTSLIHSL | 21 |
| MHC-II | H2-IAb | 13 | 27 | 15 | INNYTSLIHSLIEES | YTSLIHSLI | 23 |
| MHC-II | H2-IAb | 11 | 25 | 15 | REINNYTSLIHSLIE | NYTSLIHSL | 26 |
| MHC-II | H2-IAb | 21 | 35 | 15 | HSLIEESQNQQEKNE | IEESQNQQE | 30 |

**Supplementary Table 8 | Predicted murine MHC epitopes of the HIV HR2 peptide and associated MHC binding.** NetMHCpan-4.1 was used to predict the top eight murine MHC-I and MHC-II epitopes of each HR2 peptide. The default cut-off for MHC binders was used as described by NetMHCpan[59]. A lower rank score for the peptide is associated with stronger binding to MHC. 0.5% was defined as the cut-off for 'strong' MHC-I binders while 2% was defined as the cut-off for 'weak' MHC-I binders. Any peptides that had a rank score >2% were assumed to not bind to MHC-I. 2% was defined as the cut-off for 'strong' MHC-II binders while 10% was defined as the cut-off for 'weak' MHC-II binders. Any peptides that had a rank score >10% were assumed to not bind to MHC-II.

| MHC-I or MHC-II | Allele | Start | End | Length | Peptide | Core peptide region | Rank |
|---|---|---|---|---|---|---|---|
| MHC-I | H-2-Db | 8 | 17 | 10 | KNITDKIDQI | KNITDIDQI | 1.8 |
| MHC-I | H-2-Kb | 13 | 21 | 9 | KIDQIIHDF | KIDQIIHDF | 4 |
| MHC-I | H-2-Kb | 10 | 17 | 8 | ITDKIDQI | IT-DKIDQI | 4.9 |
| MHC-I | H-2-Db | 10 | 18 | 9 | ITDKIDQII | ITDKIDQII | 4 |
| MHC-I | H-2-Db | 8 | 18 | 11 | KNITDKIDQII | KNITIDQII | 5.3 |
| MHC-I | H-2-Db | 6 | 14 | 9 | WTKNITDKI | WTKNITDKI | 5.6 |
| MHC-I | H-2-Db | 13 | 21 | 9 | KIDQIIHDF | KIDQIIHDF | 6.1 |
| MHC-I | H-2-Kb | 9 | 17 | 9 | NITDKIDQI | NITDKIDQI | 7.4 |
| MHC-II | H2-IAb | 3 | 17 | 15 | PHDWTKNITDKIDQI | WTKNITDKI | 19 |
| MHC-II | H2-IAb | 2 | 16 | 15 | EPHDWTKNITDKIDQ | WTKNITDKI | 19 |
| MHC-II | H2-IAb | 1 | 15 | 15 | IEPHDWTKNITDKID | WTKNITDKI | 23 |
| MHC-II | H2-IAb | 4 | 18 | 15 | HDWTKNITDKIDQII | WTKNITDKI | 31 |
| MHC-II | H2-IAb | 5 | 19 | 15 | DWTKNITDKIDQIIH | WTKNITDKI | 57 |
| MHC-II | H2-IAb | 6 | 20 | 15 | WTKNITDKIDQIIHD | NITDKIDQI | 60 |
| MHC-II | H2-IAb | 7 | 21 | 15 | TKNITDKIDQIIHDF | ITDKIDQII | 62 |
| MHC-II | H2-IAb | 10 | 24 | 15 | ITDKIDQIIHDFVDK | IDQIIHDFV | 67 |

**Supplementary Table 9 | Predicted murine MHC epitopes of the Ebola HR2 peptide and associated MHC binding.** NetMHCpan-4.1 was used to predict the top eight murine MHC-I and MHC-II epitopes of each HR2 peptide. The default cut-off for MHC binders was used as described by NetMHCpan[59]. A lower rank score for the peptide is associated with stronger binding to MHC. 0.5% was defined as the cut-off for 'strong' MHC-I binders while 2% was defined as the cut-off for 'weak' MHC-I binders. Any peptides that had a rank score >2% were assumed to not bind to MHC-I. 2% was defined as the cut-off for 'strong' MHC-II binders while 10% was defined as the cut-off for 'weak' MHC-II binders. Any peptides that had a rank score >10% were assumed to not bind to MHC-II.

| Infectious disease | MHC-I strong binders | MHC-I weak binders | MHC-II strong binders | MHC-II weak binders |
|---|---|---|---|---|
| SARS-CoV-2 | 6 | >8 | 0 | 4 |
| HIV | 2 | >8 | 0 | 0 |
| Ebola | 0 | 1 | 0 | 0 |

**Supplementary Table 10 | Summary Table of 'strong' and 'weak' MHC binding epitopes for various HR2 peptides.** NetMHCpan-4.1 was used to predict the top eight murine MHC-I and MHC-II epitopes of each HR2 peptide. The default cutoff for MHC binders was used as described by NetMHCpan[59]. A lower rank score for the peptide is associated with stronger binding to MHC. 0.5% was defined as the cutoff for 'strong' MHC-I binders while 2% was defined as the cutoff for 'weak' MHC-I binders. Any peptides that had a rank score >2% were assumed to not bind to MHC-I. 2% was defined as the cutoff for 'strong' MHC-II binders while 10% was defined as the cutoff for 'weak' MHC-II binders. Any peptides that had a rank score >10% were assumed to not bind to MHC-II.

| Antibody | Fluorophore | Catalog Number | Supplier |
|---|---|---|---|
| CD1c | BV421 | 331526 | Biolegend |
| CD14 | BUV395 | 563561 | BD |
| CD86 | BV605 | 305430 | Biolegend |
| CD40 | APC/Fire™ 750 | 334345 | Biolegend |
| HLA-DR | PE-Cy7 | 307616 | Biolegend |
| CD83 | FITC | 305306 | Biolegend |
| ViaKrome 808 Fixable Viability Dye | ViaKrome 808 | C36628 | Beckman Coulter |

**Supplementary Table 11 | Antibodies and fluorescent reagents used for multiparametric flow cytometric analysis of human monocyte - derived dendritic cells (DCs).** This panel includes markers for DC lineage (CD1c, CD14), activation and co-stimulation (CD86, CD40), antigen presentation (HLA-DR) and maturation (CD83), along with a fixable viability dye (ViaKrome 808) to exclude dead cells.

| Antibody | Fluorophore | Catalog Number | Supplier |
|---|---|---|---|
| CD3 | PerCP Cy5.5 | 317336 | Biolegend |
| CD4 | BV650 | 317436 | Biolegend |
| CD8 | AF700 | 344724 | Biolegend |
| IFN$\gamma$ | BV421 | 506538 | Biolegend |
| TNFα | FITC | 502906 | Biolegend |
| IL2 | PE-Cy7 | 500326 | Biolegend |
| Mouse IgG1, κ Isotype Ctrl Antibody | BV421 | 400158 | Biolegend |
| Mouse IgG1, κ Isotype Ctrl Antibody | FITC | 400108 | Biolegend |
| Rat IgG2a, κ Isotype Ctrl Antibody | PE-Cy7 | 400522 | Biolegend |
| ViaKrome 808 Fixable Viability Dye | ViaKrome 808 | C36628 | Beckman Coulter |

**Supplementary Table 12 | Antibodies and reagents used for intracellular cytokine staining in human T cells by flow cytometry.** This panel includes T cell lineage markers (CD3, CD4, CD8) to delineate T cell subsets; intracellular cytokines (IFN$\gamma$, TNFα, IL2) to assess functional responses; appropriate isotype control antibodies (mouse IgG1, κ and rat IgG2a, κ) for specificity; and a fixable viability dye (ViaKrome 808) to discriminate live cells.

| Protein Name | Supplier | Catalog Number | Number of Lysines | Molecular Weight |
|---|---|---|---|---|
| HBsAg | Advanced Immunochemical Inc. | 7-HbADW | 4 | 25,394.7 g/mol |
| E8L | Sino Biological | 40890-V08B | 21 | 35,750 g/mol |
| H3L | Sino Biological | 40893-V08H1 | 22 | 33,690 g/mol |
| M1R | Sino Biological | 40904-V07H | 10 | 21,590 g/mol |
| SARS-CoV-2 Spike | Sino Biological | 40589-V08H8 | 61 | 136,450 g/mol |

**Supplementary Table 13 | Protein sources and relevant properties.** The number of lysines for each protein is relevant to the protein-oligonucleotide conjugation reaction, as the oligonucleotide is conjugated to lysine residues via an NHS ester reaction with the amino group on the lysines, followed by a DBCO-azide click chemistry reaction.

| HLA-A or HLA-DRB | Start | End | Length | Peptide | Core peptide region | Rank |
|---|---|---|---|---|---|---|
| HLA-A*02:01 | 19 | 27 | 9 | RLNEVAKNL | RLNEVAKNL | 0.16 |
| HLA-A*02:01 | 26 | 34 | 9 | NLNESLIDL | NLNESLIDL | 0.18 |
| HLA-A*02:01 | 30 | 37 | 8 | SLIDLQEL | SLID-LQEL | 0.37 |
| HLA-A*02:01 | 25 | 34 | 10 | KNLNESLIDL | KLNESLIDL | 0.66 |
| HLA-A*02:01 | 24 | 34 | 11 | AKNLNESLIDL | ALNESLIDL | 1.1 |
| HLA-A*02:01 | 5 | 13 | 9 | GINASVVNI | GINASVVNI | 1.1 |
| HLA-A*02:01 | 15 | 23 | 9 | KEIDRLNEV | KEIDRLNEV | 1.5 |
| HLA-A*02:01 | 13 | 23 | 11 | IQKEIDRLNEV | IQIDRLNEV | 1.5 |
| HLA-DRB1*01:11 | 1 | 15 | 15 | GDISGINASVVNIQK | ISGINASVV | 12 |
| HLA-DRB1*01:11 | 14 | 28 | 15 | QKEIDRLNEVAKNLN | IDRLNEVAK | 26 |
| HLA-DRB1*01:11 | 2 | 16 | 15 | DISGINASVVNIQKE | INASVVNIQ | 28 |
| HLA-DRB1*01:11 | 3 | 17 | 15 | ISGINASVVNIQKEI | INASVVNIQ | 29 |
| HLA-DRB1*01:11 | 13 | 27 | 15 | IQKEIDRLNEVAKNL | IDRLNEVAK | 32 |
| HLA-DRB1*01:11 | 10 | 24 | 15 | VVNIQKEIDRLNEVA | IQKEIDRLN | 34 |
| HLA-DRB1*01:11 | 12 | 26 | 15 | NIQKEIDRLNEVAKN | IDRLNEVAK | 35 |
| HLA-DRB1*01:11 | 8 | 22 | 15 | ASVVNIQKEIDRLNE | VNIQKEIDR | 38 |

**Supplementary Table 14 | Predicted human HLA epitopes of the SARS-CoV-2 HR2 peptide and associated HLA binding.** NetMHCpan-4.1 was used to predict the top eight human HLA-A*02:01 and HLA-DRB1*01:11 epitopes of each HR2 peptide. The default cutoff for HLA binders was used as described by NetMHCpan[59]. A lower rank score for the peptide is associated with stronger HLA binding. 0.5% was defined as the cutoff for 'strong' HLA-A*02:01 binders while 2% was defined as the cutoff for 'weak' HLA-A*02:01 binders. Any peptides that had a rank score >2% were assumed to not bind to HLA-A*02:01. 2% was defined as the cutoff for 'strong' HLA-DRB1*01:11 binders while 10% was defined as the cutoff for 'weak' HLA-DRB1*01:11 binders. Any peptides that had a rank score >10% were assumed to not bind to HLA-DRB1*01:11. HLA-A*02:01 and HLA-DRB1*01:11 were chosen as representative human HLAs, as they are the most well-studied HLAs of each class.

| HLA-A or HLA-DRB | Start | End | Length | Peptide | Core peptide region | Rank |
|---|---|---|---|---|---|---|
| HLA-A*02:01 | 38 | 47 | 10 | LLELDKWASL | LLLDKWASL | 1.2 |
| HLA-A*02:01 | 16 | 23 | 8 | YTSLIHSL | YTS-LIHSL | 2.7 |
| HLA-A*02:01 | 37 | 47 | 11 | ELLELDKWASL | ELLELDASL | 3.2 |
| HLA-A*02:01 | 16 | 24 | 9 | YTSLIHSLI | YTSLIHSLI | 3.9 |
| HLA-A*02:01 | 18 | 26 | 9 | SLIHSLIEE | SLIHSLIEE | 4.1 |
| HLA-A*02:01 | 18 | 27 | 10 | SLIHSLIEES | SLIHSLIES | 4.3 |
| HLA-A*02:01 | 22 | 30 | 9 | SLIEESQNQ | SLIEESQNQ | 4.7 |
| HLA-A*02:01 | 15 | 23 | 9 | NYTSLIHSL | NYTSLIHSL | 4.9 |
| HLA-DRB1*01:11 | 13 | 27 | 15 | INNYTSLIHSLIEES | YTSLIHSLI | 4.2 |
| HLA-DRB1*01:11 | 12 | 26 | 15 | EINNYTSLIHSLIEE | YTSLIHSLI | 5.7 |
| HLA-DRB1*01:11 | 14 | 28 | 15 | NNYTSLIHSLIEESQ | YTSLIHSLI | 9 |
| HLA-DRB1*01:11 | 11 | 25 | 15 | REINNYTSLIHSLIE | YTSLIHSLI | 12 |
| HLA-DRB1*01:11 | 17 | 31 | 15 | TSLIHSLIEESQNQQ | IHSLIEESQ | 29 |
| HLA-DRB1*01:11 | 16 | 30 | 15 | YTSLIHSLIEESQNQ | IHSLIEESQ | 33 |
| HLA-DRB1*01:11 | 15 | 29 | 15 | NYTSLIHSLIEESQN | YTSLIHSLI | 33 |
| HLA-DRB1*01:11 | 10 | 24 | 15 | DREINNYTSLIHSLI | YTSLIHSLI | 34 |

**Supplementary Table 15 | Predicted human HLA epitopes of the HIV HR2 peptide and associated HLA binding.** NetMHCpan-4.1 was used to predict the top eight human HLA-A*02:01 and HLA-DRB1*01:11 epitopes of each HR2 peptide. The default cutoff for HLA binders was used as described by NetMHCpan[59]. A lower rank score for the peptide is associated with stronger HLA binding. 0.5% was defined as the cutoff for ‘strong’ HLA-A*02:01 binders while 2% was defined as the cutoff for ‘weak’ HLA-A*02:01 binders. Any peptides that had a rank score >2% were assumed to not bind to HLA-A*02:01. 2% was defined as the cutoff for ‘strong’ HLA-DRB1*01:11 binders while 10% was defined as the cutoff for ‘weak’ HLA-DRB1*01:11 binders. Any peptides that had a rank score >10% were assumed to not bind to HLA-DRB1*01:11. HLA-A*02:01 and HLA-DRB1*01:11 were chosen as representative human HLAs, as they are the most well-studied HLAs of each class.

| HLA-A or HLA-DRB | Start | End | Lengt h | Peptide | Core peptide region | Rank |
|---|---|---|---|---|---|---|
| HLA-A*02:01 | 13 | 22 | 10 | KIDQIIHDFV | KIDQIIHFV | 0.71 |
| HLA-A*02:01 | 13 | 21 | 9 | KIDQIIHDF | KIDQIIHDF | 1.4 |
| HLA-A*02:01 | 9 | 17 | 9 | NITDKIDQI | NITDKIDQI | 1.6 |
| HLA-A*02:01 | 10 | 18 | 9 | ITDKIDQII | ITDKIDQII | 3.2 |
| HLA-A*02:01 | 8 | 17 | 10 | KNITDKIDQI | KITDKIDQI | 4.5 |
| HLA-A*02:01 | 10 | 17 | 8 | ITDKIDQI | ITD-KIDQI | 9.2 |
| HLA-A*02:01 | 12 | 22 | 11 | DKIDQIIHDFV | KIDQIIHFV | 9.5 |
| HLA-A*02:01 | 13 | 23 | 11 | KIDQIIHDFVD | KIDQIIHFV | 11 |
| HLA-DRB1*01:11 | 3 | 17 | 15 | PHDWTKNITDKIDQI | WTKNITDKI | 8.6 |
| HLA-DRB1*01:11 | 2 | 16 | 15 | EPHDWTKNITDKIDQ | WTKNITDKI | 9.7 |
| HLA-DRB1*01:11 | 1 | 15 | 15 | IEPHDWTKNITDKID | WTKNITDKI | 17 |
| HLA-DRB1*01:11 | 4 | 18 | 15 | HDWTKNITDKIDQII | WTKNITDKI | 20 |
| HLA-DRB1*01:11 | 7 | 21 | 15 | TKNITDKIDQIIHDF | ITDKIDQII | 35 |
| HLA-DRB1*01:11 | 10 | 24 | 15 | ITDKIDQIIHDFVDK | IDQIIHDFV | 36 |
| HLA-DRB1*01:11 | 6 | 20 | 15 | WTKNITDKIDQIIHD | ITDKIDQII | 38 |
| HLA-DRB1*01:11 | 5 | 19 | 15 | DWTKNITDKIDQIIH | WTKNITDKI | 43 |

**Supplementary Table 16 | Predicted human HLA epitopes of the Ebola HR2 peptide and associated HLA binding.** NetMHCpan-4.1 was used to predict the top eight human HLA-A*02:01 and HLA-DRB1*01:11 epitopes of each HR2 peptide. The default cutoff for HLA binders was used as described by NetMHCpan[59]. A lower rank score for the peptide is associated with stronger HLA binding. 0.5% was defined as the cutoff for 'strong' HLA-A*02:01 binders while 2% was defined as the cutoff for 'weak' HLA-A*02:01 binders. Any peptides that had a rank score >2% were assumed to not bind to HLA-A*02:01. 2% was defined as the cutoff for 'strong' HLA-DRB1*01:11 binders while 10% was defined as the cutoff for 'weak' HLA-DRB1*01:11 binders. Any peptides that had a rank score >10% were assumed to not bind to HLA-DRB1*01:11. HLA-A*02:01 and HLA-DRB1*01:11 were chosen as representative human HLAs, as they are the most well-studied HLAs of each class.

| Infectious disease | HLA-A strong binders | HLA-A weak binders | HLA-DRB strong binders | HLA-DRB weak binders |
|---|---|---|---|---|
| SARS-CoV-2 | 3 | >8 | 0 | 0 |
| HIV | 0 | 1 | 0 | 3 |
| Ebola | 0 | 3 | 0 | 2 |

**Supplementary Table 17 | Summary table of 'strong' and 'weak' HLA binding epitopes for various HR2 peptides.** NetMHCpan-4.1 was used to predict the top eight human HLA-A*02:01 and HLA-DRB1*01:11 epitopes of each HR2 peptide. The default cutoff for HLA binders was used as described by NetMHCpan[59]. A lower rank score for the peptide is associated with stronger HLA binding. 0.5% was defined as the cutoff for 'strong' HLA-A*02:01 binders while 2% was defined as the cutoff for 'weak' HLA-A*02:01 binders. Any peptides that had a rank score >2% were assumed to not bind to HLA-A*02:01. 2% was defined as the cutoff for 'strong' HLA-DRB1*01:11 binders while 10% was defined as the cutoff for 'weak' HLA-DRB1*01:11 binders. Any peptides that had a rank score >10% were assumed to not bind to HLA-DRB1*01:11. HLA-A*02:01 and HLA-DRB1*01:11 were chosen as representative human HLAs, as they are the most well-studied HLAs of each class.

| Antibody | Fluorophore | Catalog Number | Supplier |
|---|---|---|---|
| I-A/I-E (MHC II) | Brilliant Violet 421™ | 107632 | Biolegend |
| CD11b | Brilliant Violet 605™ | 101257 | Biolegend |
| CD103 | Alexa Fluor® 700 | 121442 | Biolegend |
| CD86 | PE/Cyanine7 | 105014 | Biolegend |
| CD205 (DEC-205) | PerCP/Cyanine5.5 | 138208 | Biolegend |
| CD11c | Brilliant Violet 510™ | 117338 | Biolegend |
| CD3 | Brilliant Violet 785™ | 100232 | Biolegend |
| CD4 | PE | 100408 | Biolegend |
| CD8a | APC/Cyanine7 | 100714 | Biolegend |
| CD107a (LAMP-1) | Alexa Fluor® 488 | 121608 | Biolegend |
| CD279 (PD-1) | PerCP/Cyanine5.5 | 135208 | Biolegend |
| IFN-γ | Brilliant Violet 510™ | 505841 | Biolegend |
| CD69 | PE/Cyanine7 | 104512 | Biolegend |
| NK-1.1 | PE | 156504 | Biolegend |
| T-bet | Brilliant Violet 421™ | 644832 | Biolegend |
| CD40 | PE | 124610 | Biolegend |
| CD64 | FITC | 139316 | Biolegend |

**Supplementary Table 18 | Antibodies used for flow cytometry in Cov-2-Spike DoriVac immunogenicity studies.** This panel was used to characterize immune cell phenotypes and activation in S-protein-conjugated DoriVac-treated mice and includes markers for antigen-presenting cells (I-A/I-E, CD11b, CD11c, CD40, CD64, CD86, CD205), T cells (CD3, CD4, CD8a, CD69, CD279, IFN-γ, T-bet), NK cells (NK-1.1, CD107a), and additional markers (CD103).

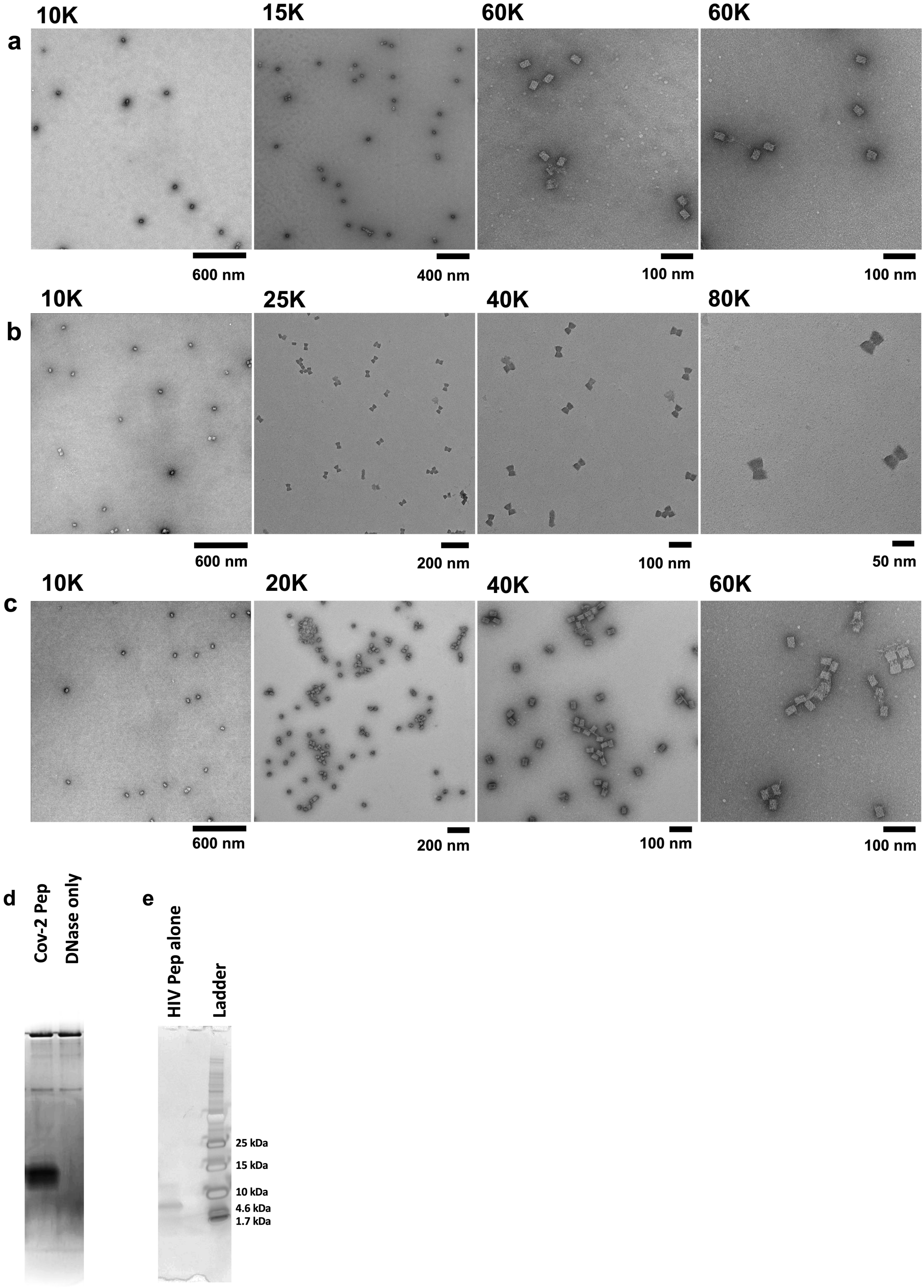


**Supplementary Fig. 1 | Additional TEM images of HR2-functionalized DoriVac nanoparticles and SDS-PAGE analysis of associated peptides. a,** Negative stain TEM image of DoriVac nanoparticles demonstrated that the nanoparticles were monodispersed, further supporting the agarose gel results. **b,**

Negative stain TEM image of HIV nanoparticles demonstrated that the nanoparticles were majority monodispersed, with some dimers observed, further supporting the agarose gel results. **c,** Negative stain TEM image of Ebola nanoparticles demonstrated that the nanoparticles were majority monodispersed, with some dimers observed, further supporting the agarose gel results. **d,** SDS-PAGE gel showing DNase I only alongside DNase I digested Cov-2 peptide. **e,** SDS-PAGE gel showing HIV peptide alone, with two bands located near the 10 kDa ladder marker.

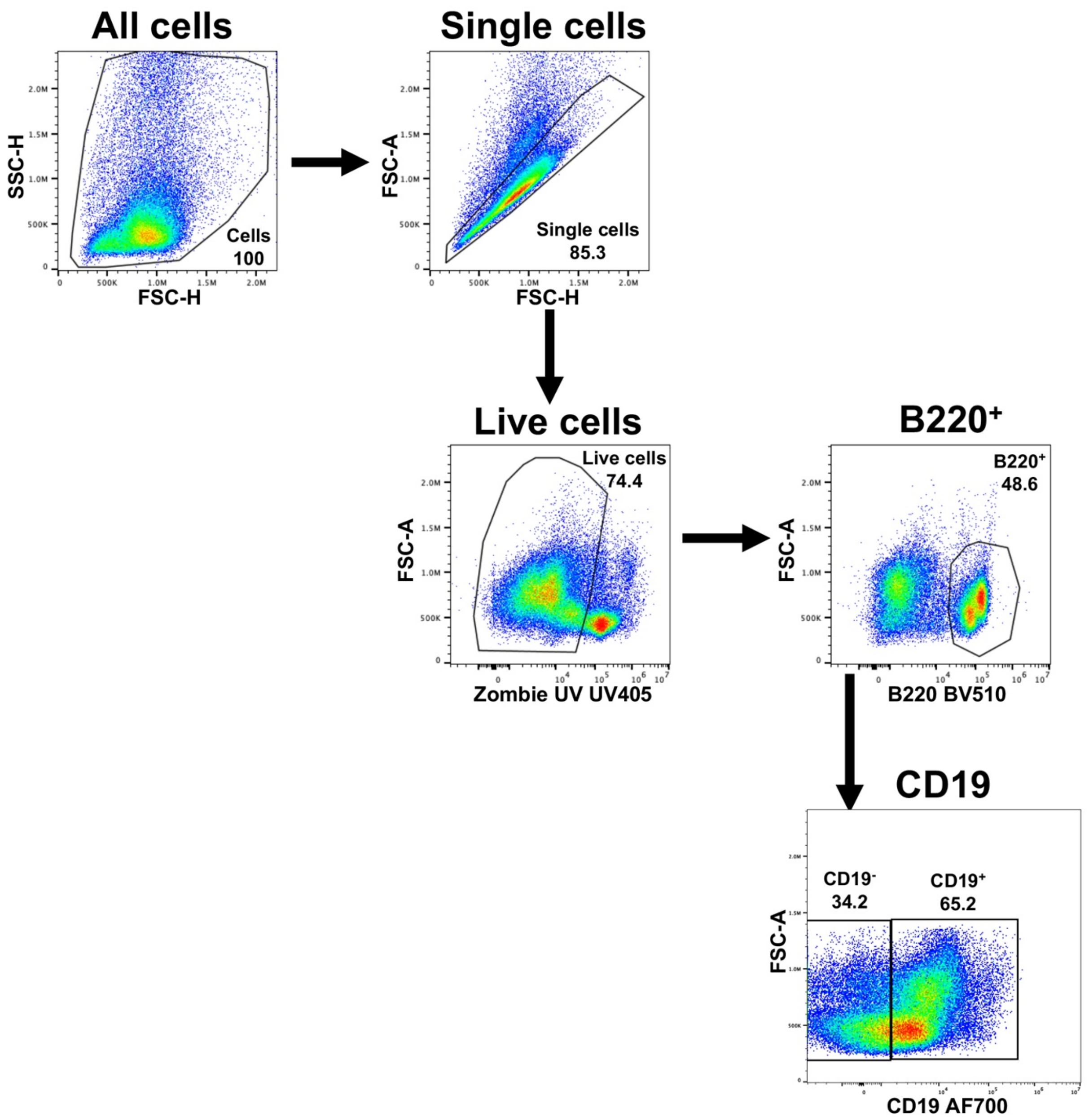


**Supplementary Fig. 2 | B cell gating strategies.** B cells were analysed using flow cytometry. This figure demonstrates the gating strategy to analyse B cells from frozen PBMC samples.

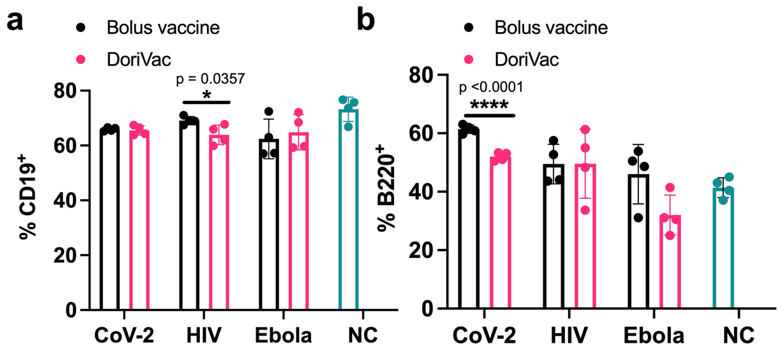


**Supplementary Fig. 3 | Additional data demonstrating robust humoral responses for B cells. a,** The CD19⁺ B cell population in the blood on Day 21, as determined by flow cytometry, showed no significant difference after bolus vaccine treatment compared with DoriVac treatment. **b,** The B220⁺ B cell population in the blood on Day 21, as determined by flow cytometry, showed minimal difference after bolus vaccine treatment compared with DoriVac treatment. Data are represented as mean ± SD. The flow data were analyzed by multiple unpaired t-tests and significance was defined as a two-tailed p value less than 0.05. '*' refers to $p \leq 0.05$; '****' refers to $p \leq 0.0001$.

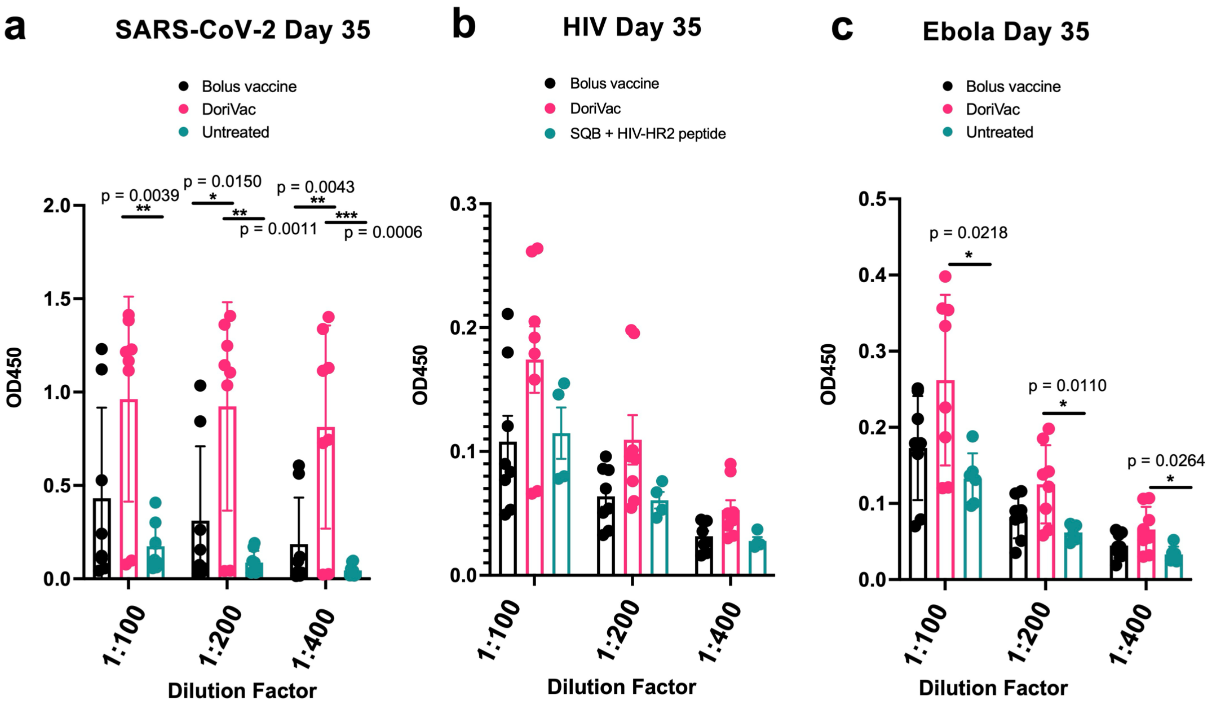


**Supplementary Fig. 4 | Additional data demonstrating robust IgG antibody production. a,** DoriVac treatment enhanced SARS-CoV-2 peptide-specific IgG antibody production in the plasma, as determined by ELISA assay, after two doses of the vaccine (on Day 35) compared to a bolus vaccine of free peptide and free CpG. **b,** DoriVac treatment enhanced HIV-peptide-specific IgG antibody production in the plasma, as determined by ELISA assay, after two doses of the vaccine (on Day 35) compared to a bolus vaccine. **c,** DoriVac treatment enhanced Ebola-peptide-specific IgG antibody production in the plasma, as determined by ELISA assay, after two doses of the vaccine (on Day 35) compared to a bolus vaccine. Data has been normalized. Data are represented as mean ± SD. The ELISA data was analyzed by one-way ANOVA (with correction for multiple comparisons using a Tukey's test) and significance was defined as a multiplicity-adjusted p value less than 0.05 (n=8). '*' refers to $P \leq 0.05$; '**' refers to $P \leq 0.01$. '***' refers to $P \leq 0.001$.

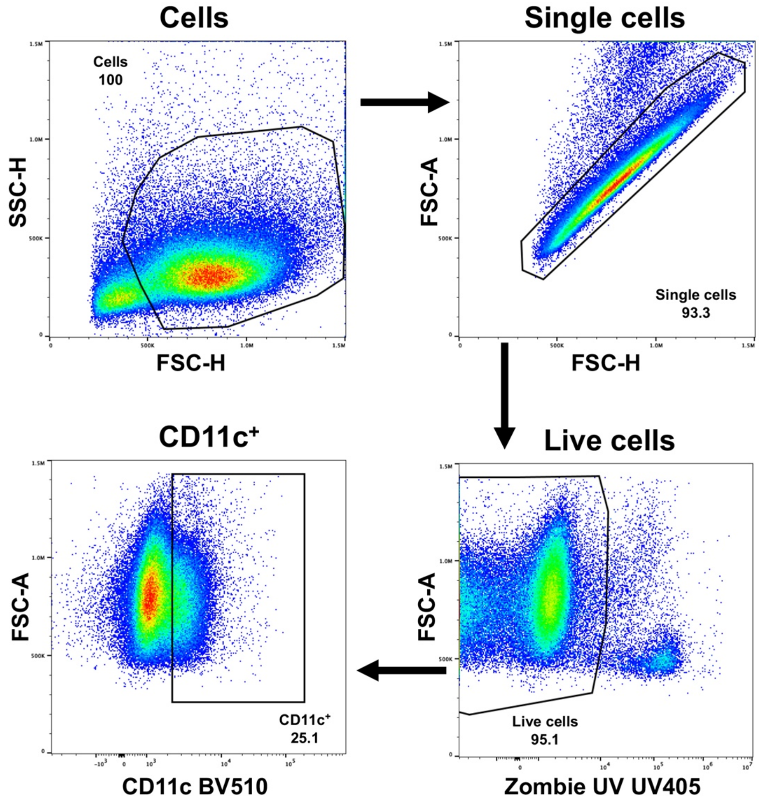


**Supplementary Fig. 5 | DC gating strategies for lymph node samples.** Dendritic cells were analyzed using flow cytometry. This figure demonstrates the gating strategy to analyze dendritic cells collected from the lymph nodes.

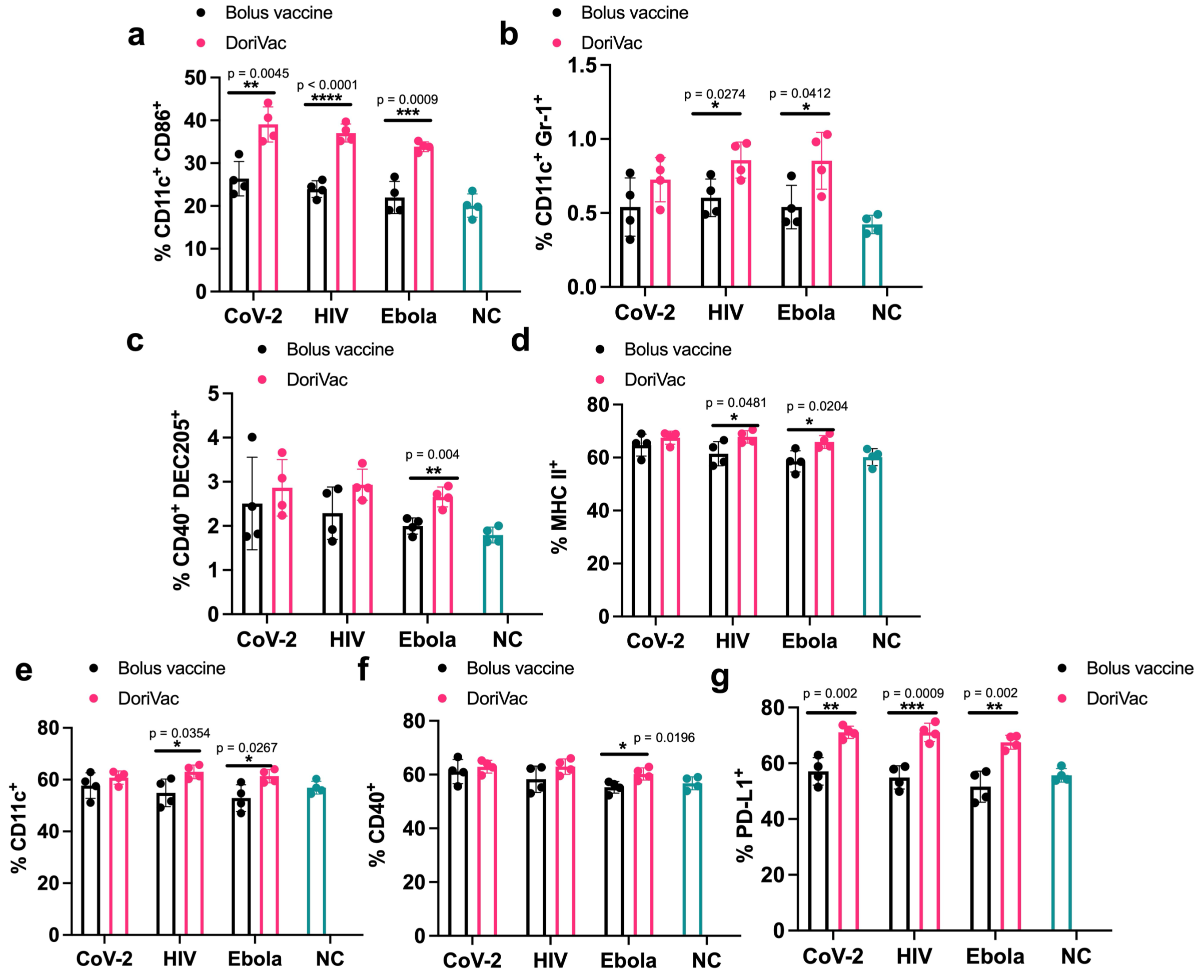


**Supplementary Fig. 6 | Additional data demonstrating DC responses in the splenocytes on Day 21.** Spleens were collected on Day 21 after two doses of treatment and processed into single-cell suspensions for flow cytometry. **a,** Percentages of $CD11c^+$ $CD86^+$, as determined by flow cytometry, in the splenocyte population (n=4). DoriVac demonstrated a significant increase in this activated DC population compared to bolus-vaccine treatment. **b,** Percentages of $CD11c^+$ $Gr\text{-}1^+$ DCs, as determined by flow cytometry, in the splenocyte population (n=4). DoriVac treatment demonstrated an increase in this plasmacytoid DC (pDC) – like subpopulation compared to bolus-vaccine treatment. **c,** Percentages of $CD40^+$ $DEC205^+$ DCs, as determined by flow cytometry, in the splenocytes (n=4). DoriVac treatment demonstrated an increase in this activated, endocytic subpopulation compared to bolus-vaccine treatment. **d,** Percentages of MHC-$II^+$ DCs, as determined by flow cytometry, in the draining lymph node (n=4). DoriVac treatment demonstrated a significant increase in this activated DC population compared to bolus-vaccine treatment. **e,** DoriVac treatment led to a notable increase in the $CD11c^+$ dendritic-cell population, as determined by flow cytometry. **f,** The lymphocytes were more activated (as indicated by upregulation of $CD40^+$) after DoriVac treatment compared to bolus vaccine treatment, as determined by flow cytometry. **g,** An increase in the PD-$L1^+$ subpopulation was observed after DoriVac treatment, compared to bolus-vaccine treatment, as determined by flow cytometry. Data are represented as mean ± SD. The flow data were analyzed by multiple unpaired t-tests and significance was defined as a two-tailed p value less than 0.05. '*' refers to $p \leq 0.05$; '**' refers to $p \leq 0.01$; '***' refers to $p \leq 0.001$; '****' refers to $p \leq 0.0001$.

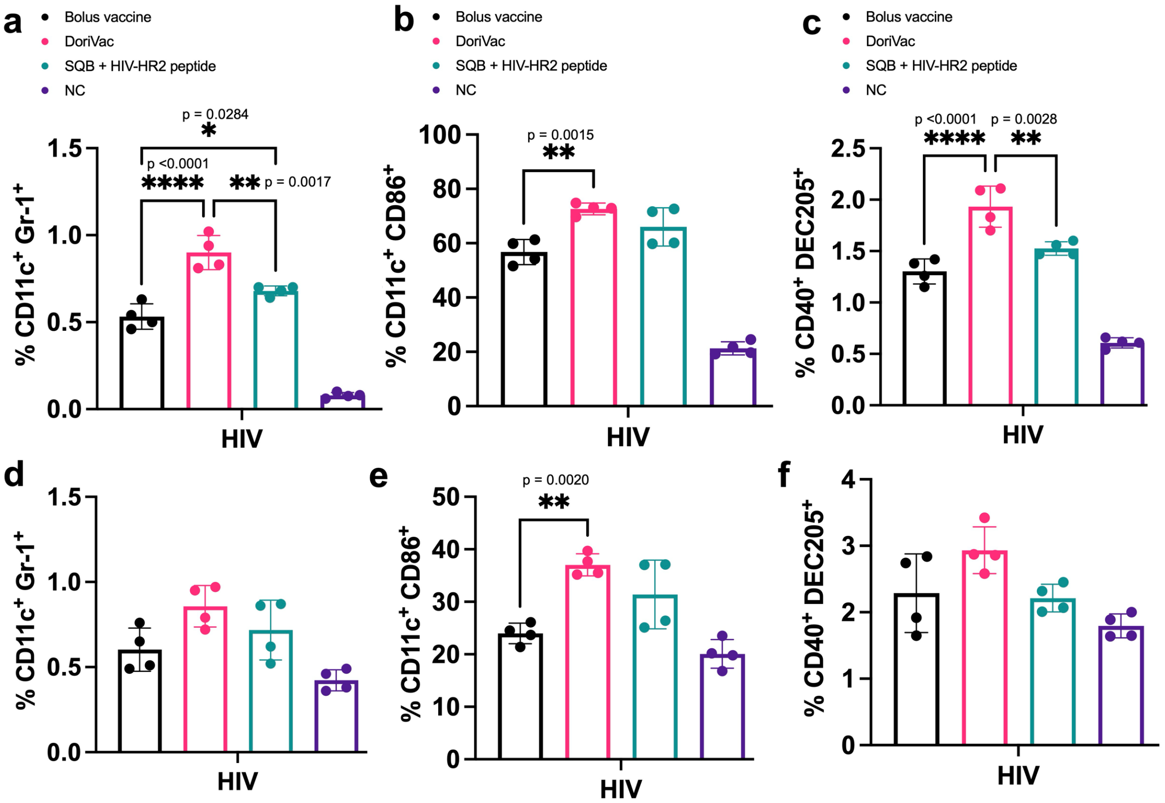


**Supplementary Fig. 7 | Co-delivery of CpG and peptide on SQB is critical to robust DC responses.** Half of the mice were sacrificed on Day 21 and draining lymph nodes were collected. **a,** Percentages of human pDC-like (CD11c$^+$ Gr-1$^+$) DCs in the draining lymph node (n=4), as determined by flow cytometry. DoriVac treatment demonstrated a significant increase in this plasmacytoid DC (pDC) – like subpopulation compared to treatment with the CpG SQB and HIV peptide, administered as a solution. **b,** Percentages of activated cDCs (CD11c$^+$ CD86$^+$) in the draining lymph node, as determined by flow cytometry. DoriVac treatment showed significant activation compared to the bolus vaccine and slightly more than the DNA origami SQB and HIV peptide administered as separate components. **c,** Percentages of CD40$^+$ DEC205$^+$ DCs in the draining lymph node (n=4), as determined by flow cytometry. The DoriVac treatment demonstrated a significant increase in this activated, endocytic subpopulation compared to bolus vaccine treatment and more than the DNA origami SQB and HIV peptide administered as separate components. **d,** Percentages of human pDC-like DCs in the splenocytes on Day 21 (n=4), as determined by flow cytometry. DoriVac treatment shows a significant increase compared to the bolus control. **e,** Percentages of activated cDCs in the splenocyte population, as determined by flow cytometry. DoriVac induced a significant increase in this population compared to the bolus control. **f,** Percentages of activated, endocytic DCs in the splenocytes population, as determined by flow cytometry. DoriVac showed significant increase in this population compared to the DNA origami SQB and HIV peptide delivered separately. Data are represented as mean ± SD. The flow data was analyzed by one-way ANOVA (with correction for multiple comparisons using a Tukey's test) and significance was defined as a multiplicity-adjusted p value less than 0.05 (n=4). '*' refers to P ≤ 0.05; '**' refers to P ≤ 0.01; '***' refers to P ≤ 0.001; '****' refers to P ≤ 0.0001.

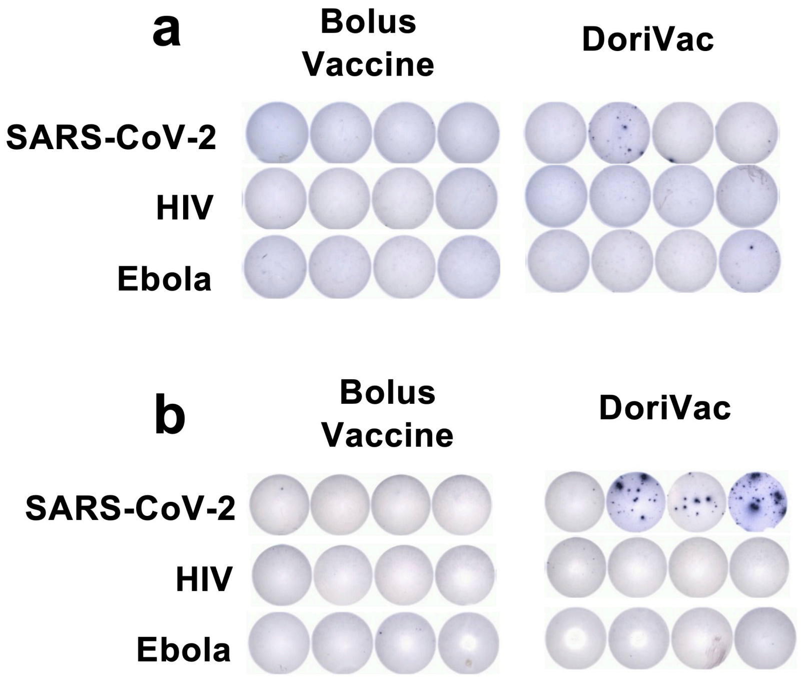


**Supplementary Fig. 8 | Additional data on antigen-specific T cell responses on Day 21 and 28. a,** Interferon-gamma (IFNγ) ELISpot demonstrating frequency of antigen-specific splenocytes on Day 21. Minimal difference is observed between the bolus-vaccine treated group and the DoriVac-treated group. **b,** IFNγ ELISpot demonstrating frequency of antigen-specific PBMCs on Day 28. There is a notable increase in the frequency of SARS-CoV-2 antigen-specific T cells.

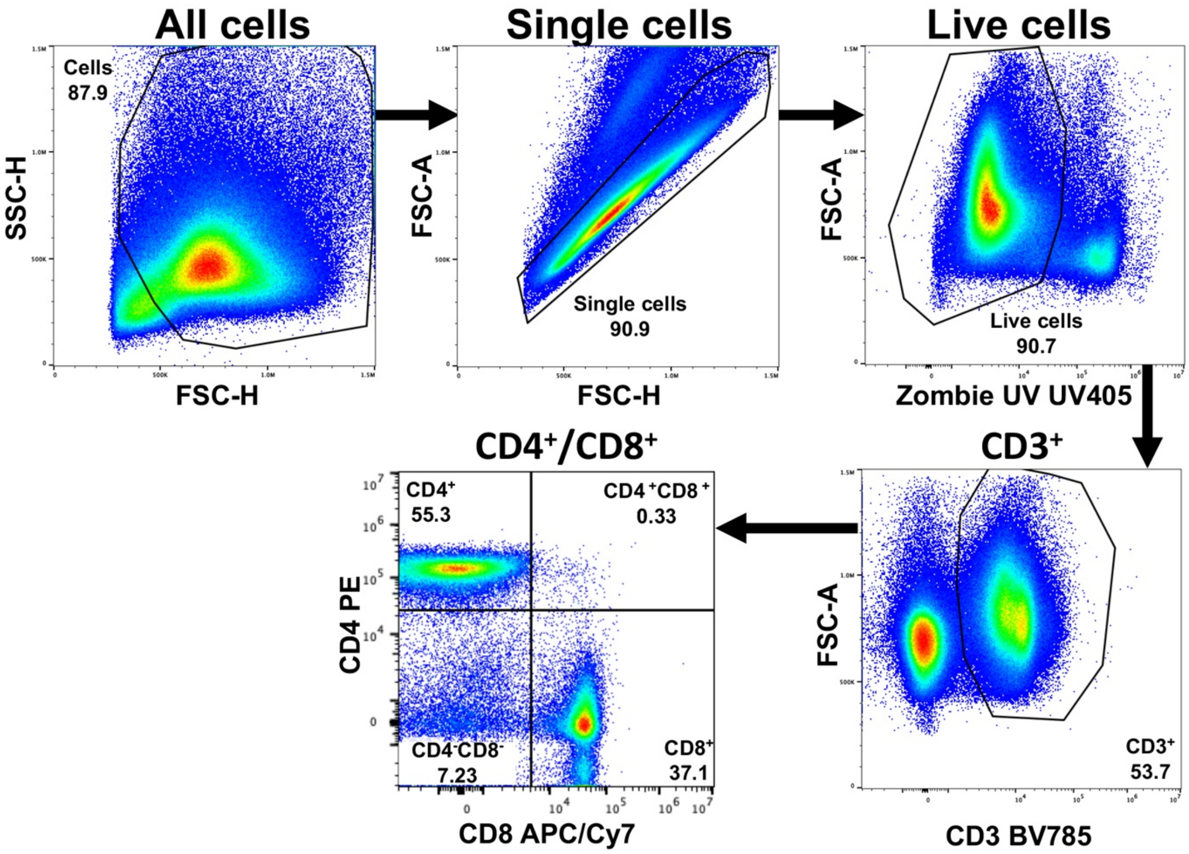


**Supplementary Fig. 9 | T cell gating strategies.** T cells were analyzed using flow cytometry. This figure demonstrates the gating strategy to analyze T cells in lymph nodes and spleen. Images shown are from the lymph nodes.

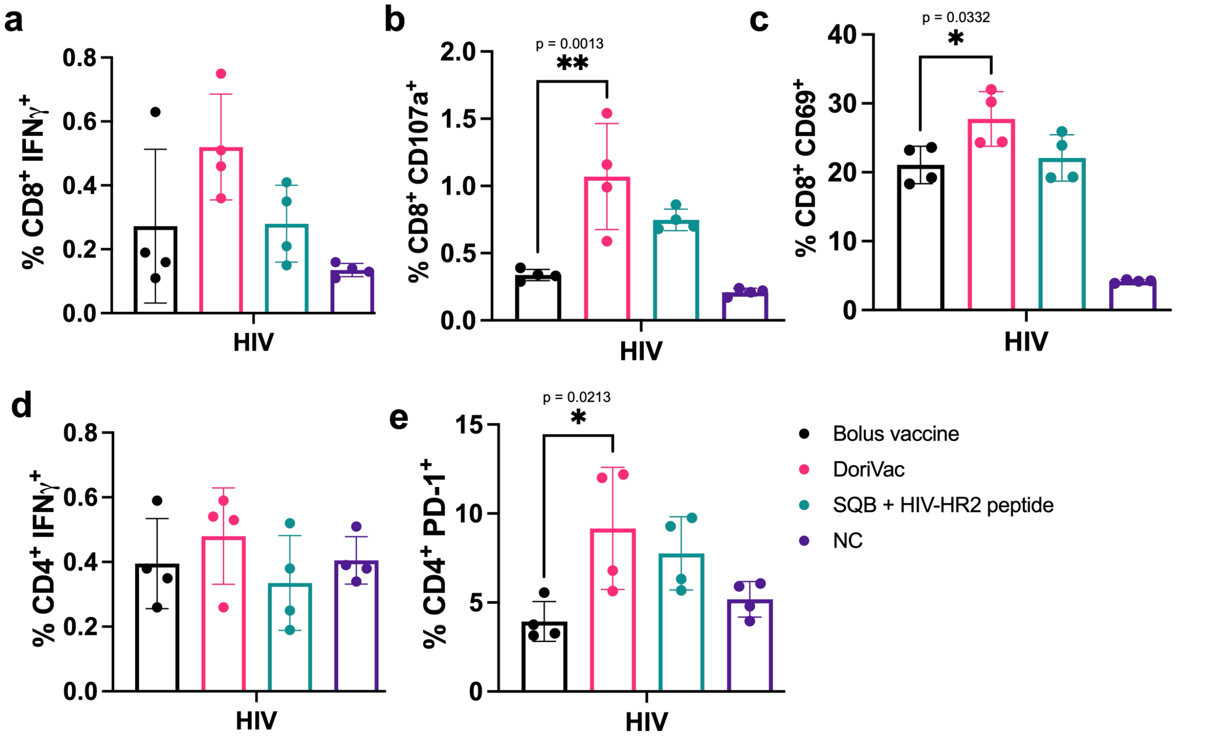


**Supplementary Fig. 10 | Co-delivery of CpG and antigen peptide is essential to robust T cell responses. a,** IFN-γ⁺ CD8⁺ T cells in the lymph node (LN) were upregulated on Day 21 in the DoriVac treatment group, compared to the bolus vaccine or the DNA origami SQB with HIV peptide groups, as determined by flow cytometry. **b,** CD107a⁺ CD8⁺ cytotoxic T cells in the LN were upregulated on Day 21 in the DoriVac treatment group, compared to the bolus vaccine or the DNA origami SQB with HIV peptide groups, as determined by flow cytometry. **c,** CD69⁺ CD8⁺ activated T cells in the LN showed upregulation on Day 21 after treatment with DoriVac, as determined by flow cytometry. **d,** IFN-γ⁺ CD4⁺ T cells in the LN were upregulated in the context of DoriVac treatment group on Day 21, compared to the DNA origami SQB with HIV peptide group, as determined by flow cytometry. **e,** PD-1⁺ CD4⁺ T cells in the LN were upregulated in the context of the DoriVac treated group on Day 21, compared to the bolus vaccine and DNA origami SQB with HIV peptide group, as determined by flow cytometry.  Data are represented as mean ± SD. The flow data was analyzed by one-way ANOVA (with correction for multiple comparisons using a Tukey's test) and significance was defined as a multiplicity-adjusted p value less than 0.05 (n=4). '*' refers to P ≤ 0.05; '**' refers to P ≤ 0.01.

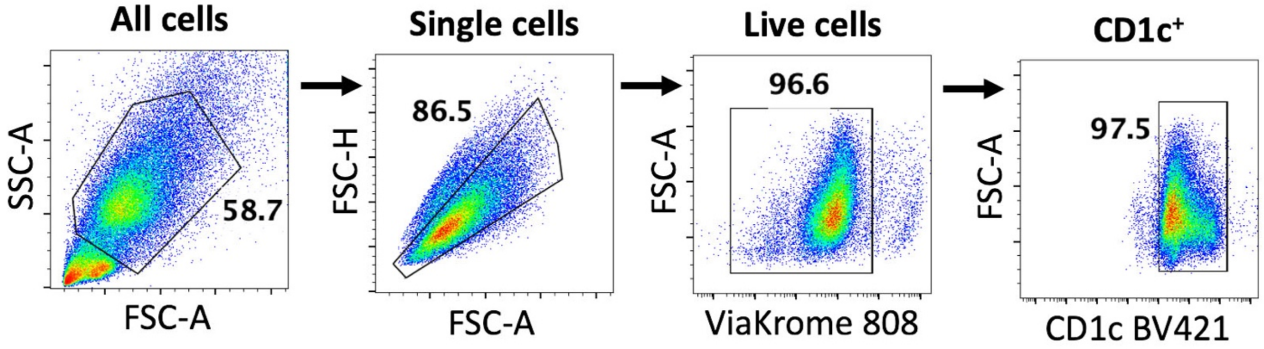


**Supplementary Fig. 11 | Activation of human monocyte-derived dendritic cells (moDCs) analysis using flow cytometry.** This figure demonstrates the gating strategy to analyse cells collected from moDCs after treatment with DNA origami vaccines.

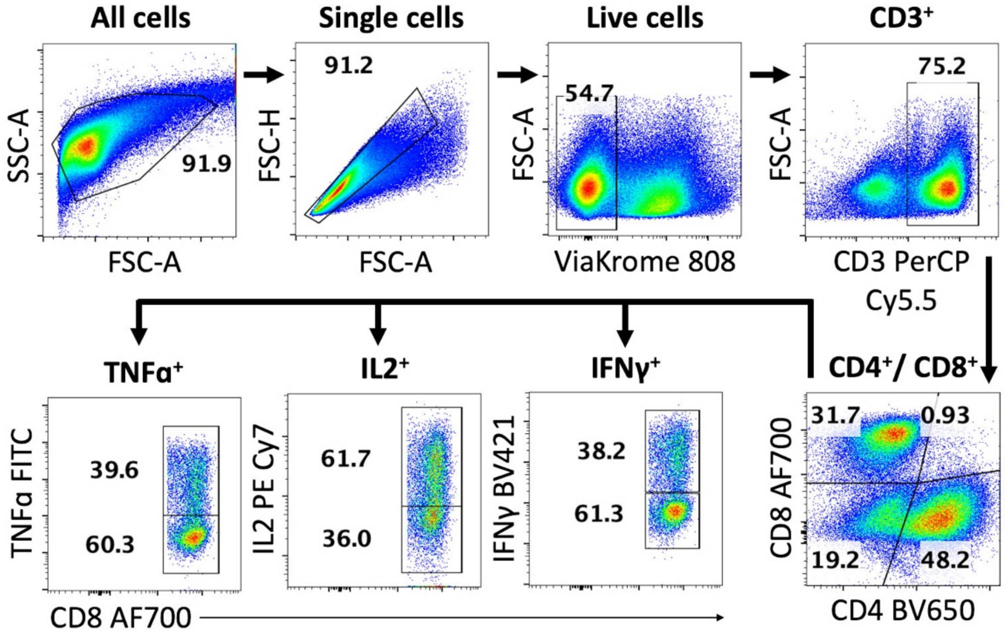


**Supplementary Fig. 12 | Intracellular cytokine staining analysis using flow cytometry for cells from LN organ-on-a-chips.** This figure demonstrates the gating strategy to analyse cells collected from LN organ-on-a-chips.

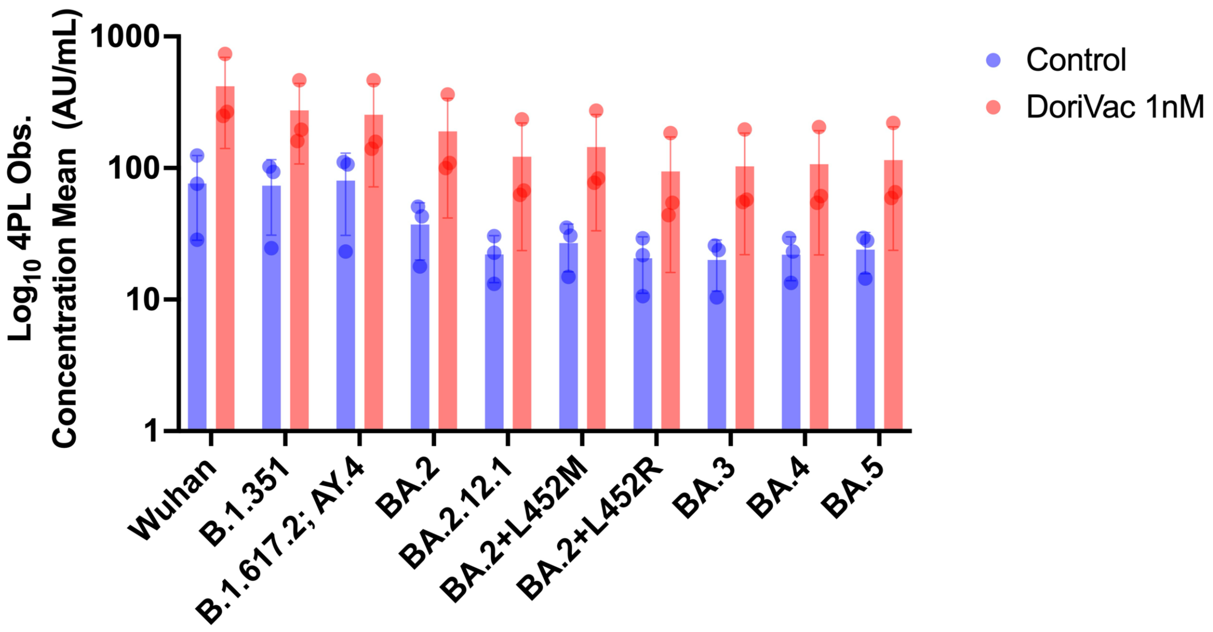


**Supplementary Fig. 13 | Broad antibody response induced by HR2-DoriVac in a human lymph-node-on-chip model against SARS-CoV-2 spike variants.** Concentration of antibodies (log10, 4PL fit observed) measured by Meso Scale Discovery (MSD) assay against SARS-CoV-2 spike protein from different variants of concern. Each bar represents the mean antibody concentration across three independent chips (n=3) in the presence of DoriVac (1 nM, red bars) and untreated control (blue bars). Variants include Wuhan, B.1.351, B.1.617.2; AY.4 Alt Seq 2, BA.2, BA.2.12.1, BA.2+L452M, BA.2+L452R, BA.3, BA.4, and BA.5. DoriVac treatment elicited higher antibody responses compared to the control across all tested variants, demonstrating a broad and robust humoral immune response.

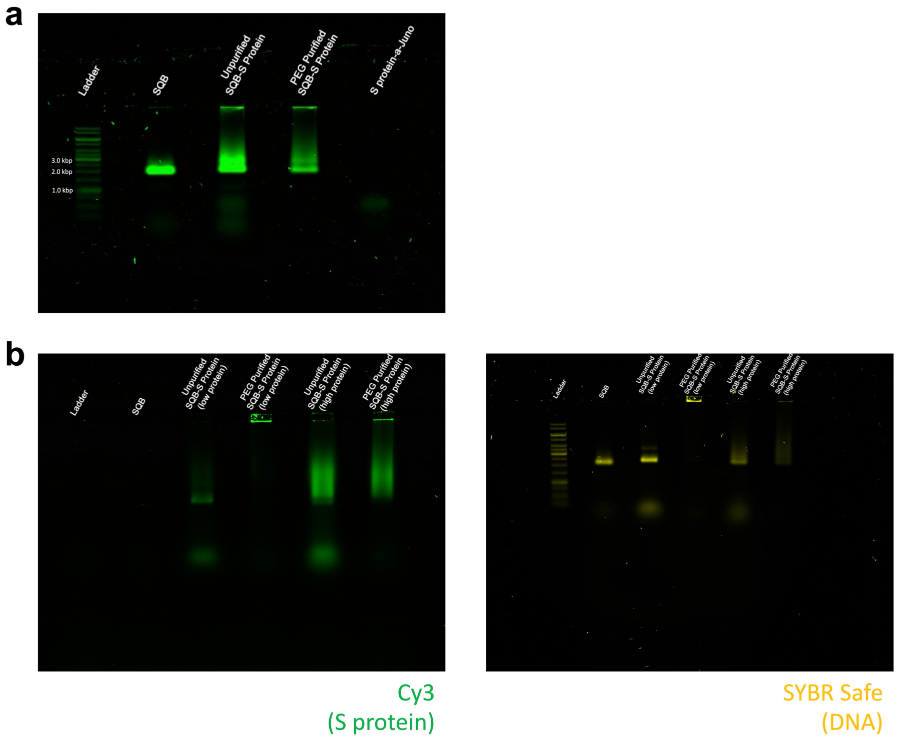


**Supplementary Fig. 14 | Conjugation of Cov-2 Spike S Protein to DoriVac SQB DNA origami. a,** Agarose gel electrophoresis demonstrating the successful attachment of oligonucleotide-conjugated S-proteins (S protein-a-Juno) to SQB origamis, resulting in upward band shifts in both the unpurified and PEG-purified DoriVac samples. PEG-purified samples were used for subsequent animal studies. **b,** Agarose gel electrophoresis demonstrating the complete removal of excess unbound oligonucleotide-conjugated Cy3-labelled Cov-2 spike S proteins from S-protein-conjugated SQBs after PEG purification (left, Cy3 green channel). The gel was subsequently post stained with SYBR safe to visualize the DNA origamis and oligonucleotides (right, yellow channel). We tested oligonucleotide-conjugated proteins added to SQBs in 40x excess of the amount of SQBs (high protein) and 11x (low protein).

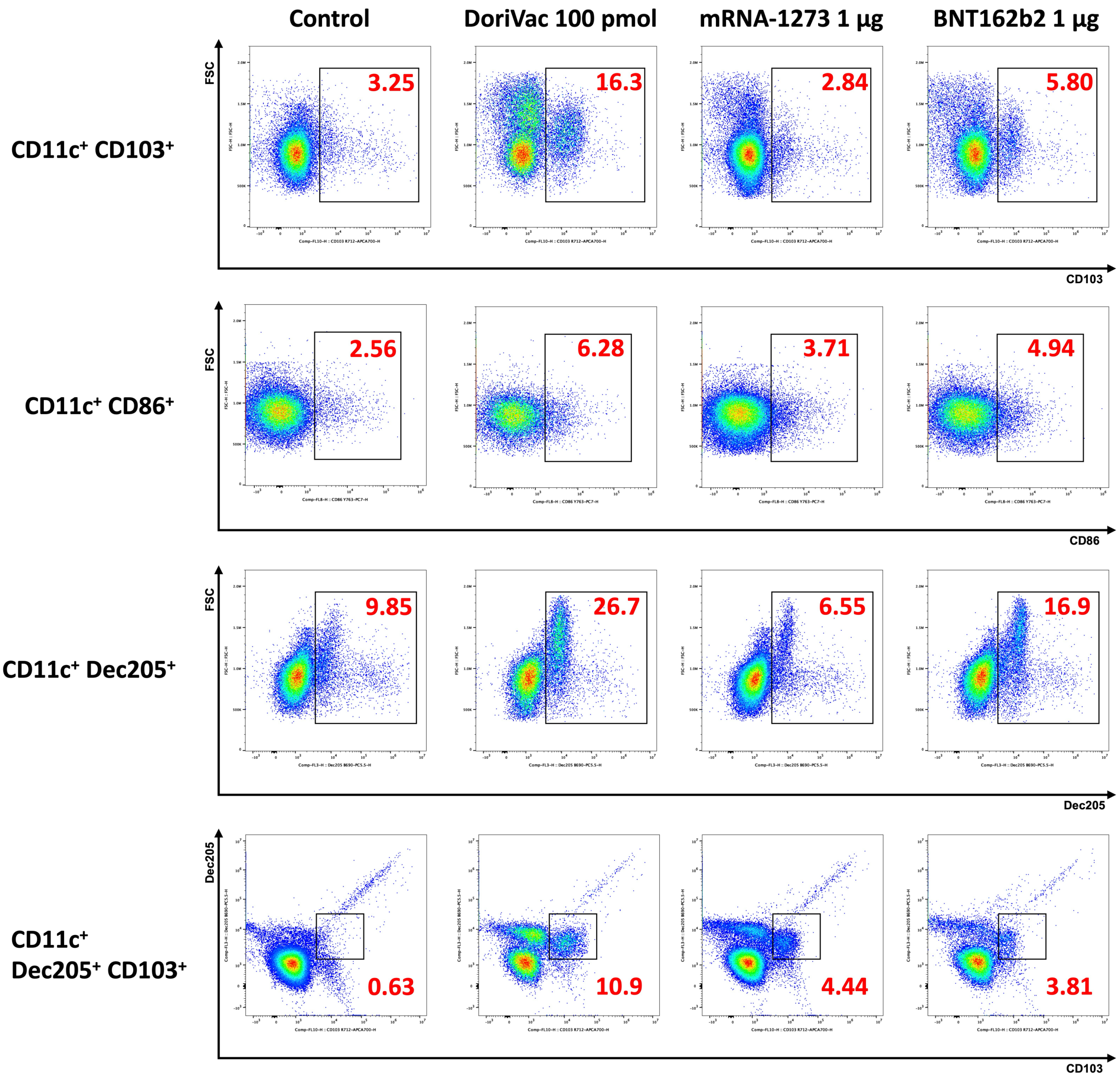


**Supplementary Fig. 15 | Immune activation in CD11c⁺ PBMCs on day 22, one day after the second dose.** Representative images from flow scatter plots and corresponding frequencies of CD11c⁺ population. Gating strategy is shown in Supplementary Fig. 5.

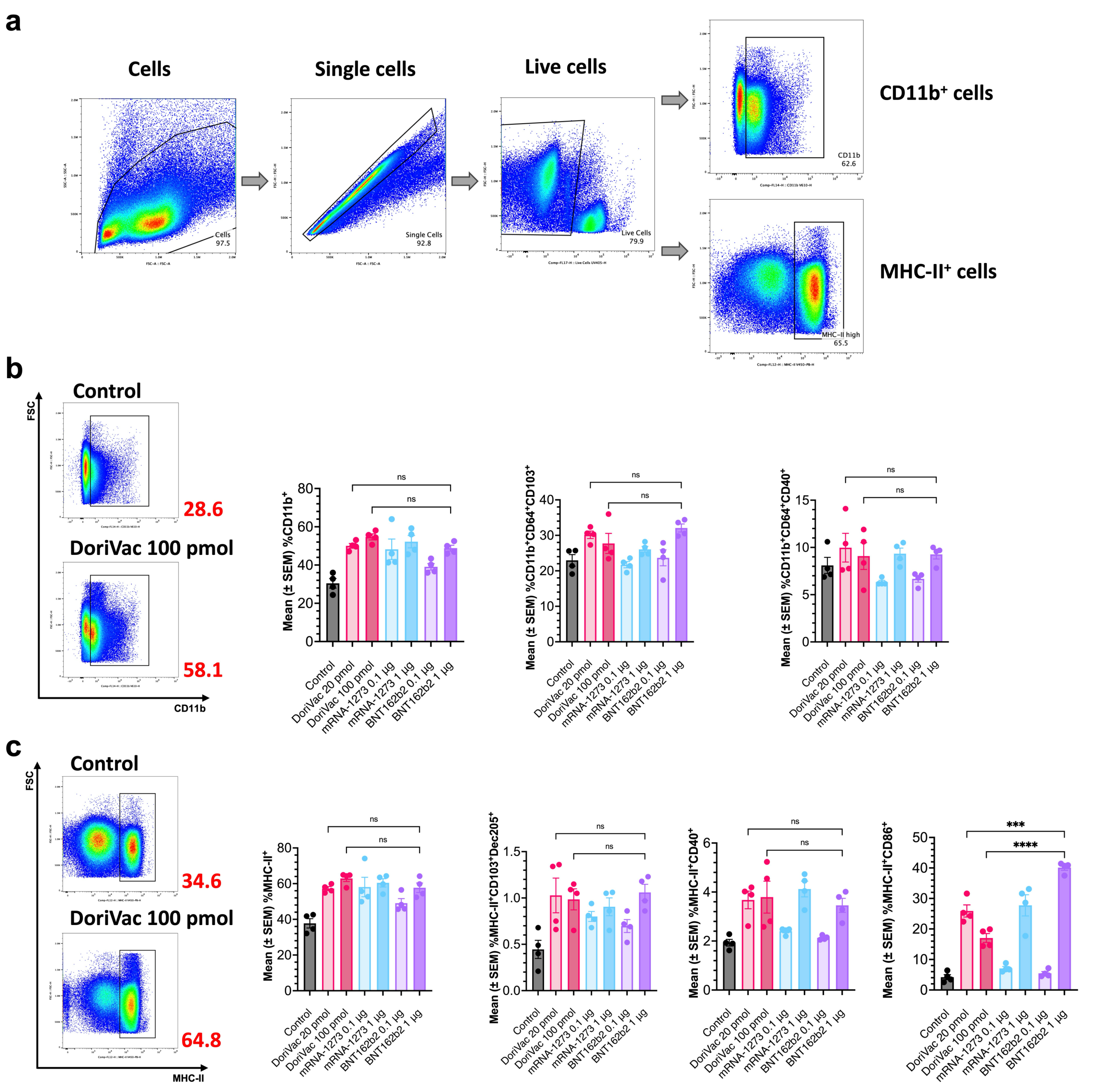


**Supplementary Fig. 16 | Immune activation in the lymphatic cell population on day 50, one day after the third dose. a,** Cell gating strategy for CD11b⁺ myeloid cells and MHC-II⁺ APCs in the lymph node (LN). **b,** CD11b⁺ myeloid cells and cross-presenting dendritic cells in the LN were upregulated on day 50 in the DoriVac treatment groups, compared to the untreated control, as determined by flow cytometry. DoriVac treatment groups showed responses comparable to the mRNA-LNP groups. Representative images from flow scatter plots and corresponding frequencies of CD11b⁺ population are shown on left. **c,** MHC-II⁺ APCs in the LN were upregulated on day 50 in the DoriVac treatment groups, compared to the untreated control, as determined by flow cytometry. DoriVac treatment groups showed responses comparable to mRNA-LNP groups for most APC activation markers except the BNT162b2 1 µg group for MHC-II⁺ CD86⁺ APCs. Representative images from flow scatter plots and corresponding frequencies of MHC-II⁺ population are shown on left. The flow data was analyzed by one-way ANOVA (with correction for multiple comparisons using a Tukey's test) and significance was defined as a multiplicity-adjusted p value less than 0.05 (n=4). 'ns' refers to P > 0.05; '***' refers to P ≤ 0.001; '****' refers to P ≤ 0.0001.

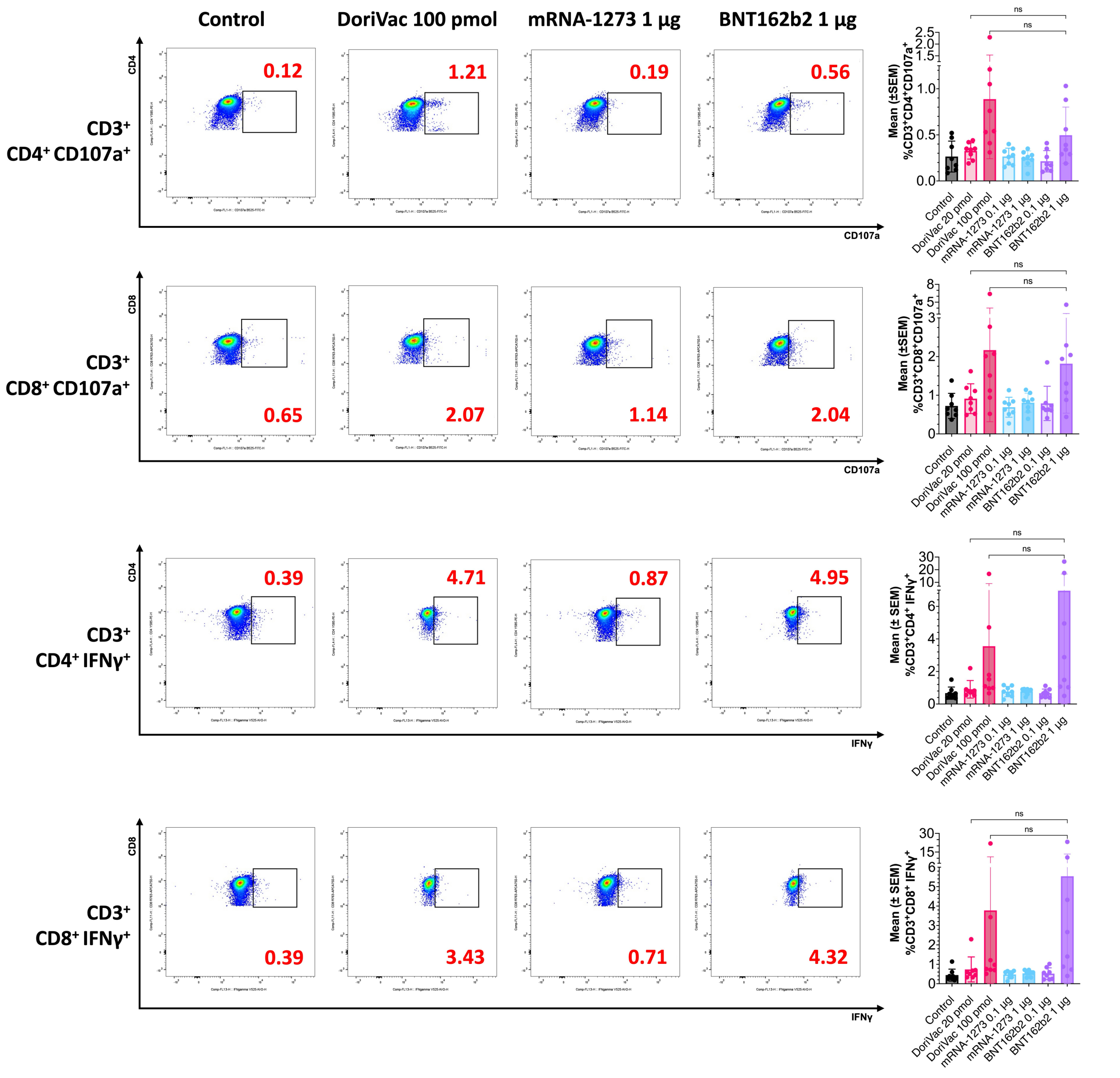


**Supplementary Fig. 17 | Percentages of CD107⁺ and IFNγ-secreting PBMCs on day 22, one day after the second dose.** Percentages of CD107⁺ PBMCs (n=8) and IFNγ secreting PBMCs (n=8) on day 22 as determined by flow cytometry. Gating strategy is illustrated in Supplementary Fig. 9. Representative images from flow scatter plots and corresponding frequencies are shown on the left. CD107a⁺ CD4⁺ and CD8⁺ PBMCs were upregulated on day 22 in the DoriVac treatment groups, compared to the untreated control. DoriVac treatment groups showed responses comparable to the mRNA-LNP groups. The flow data was analyzed by one-way ANOVA (with correction for multiple comparisons using a Tukey's test) and significance was defined as a multiplicity-adjusted p value less than 0.05 (n=8). 'ns' refers to P > 0.05.

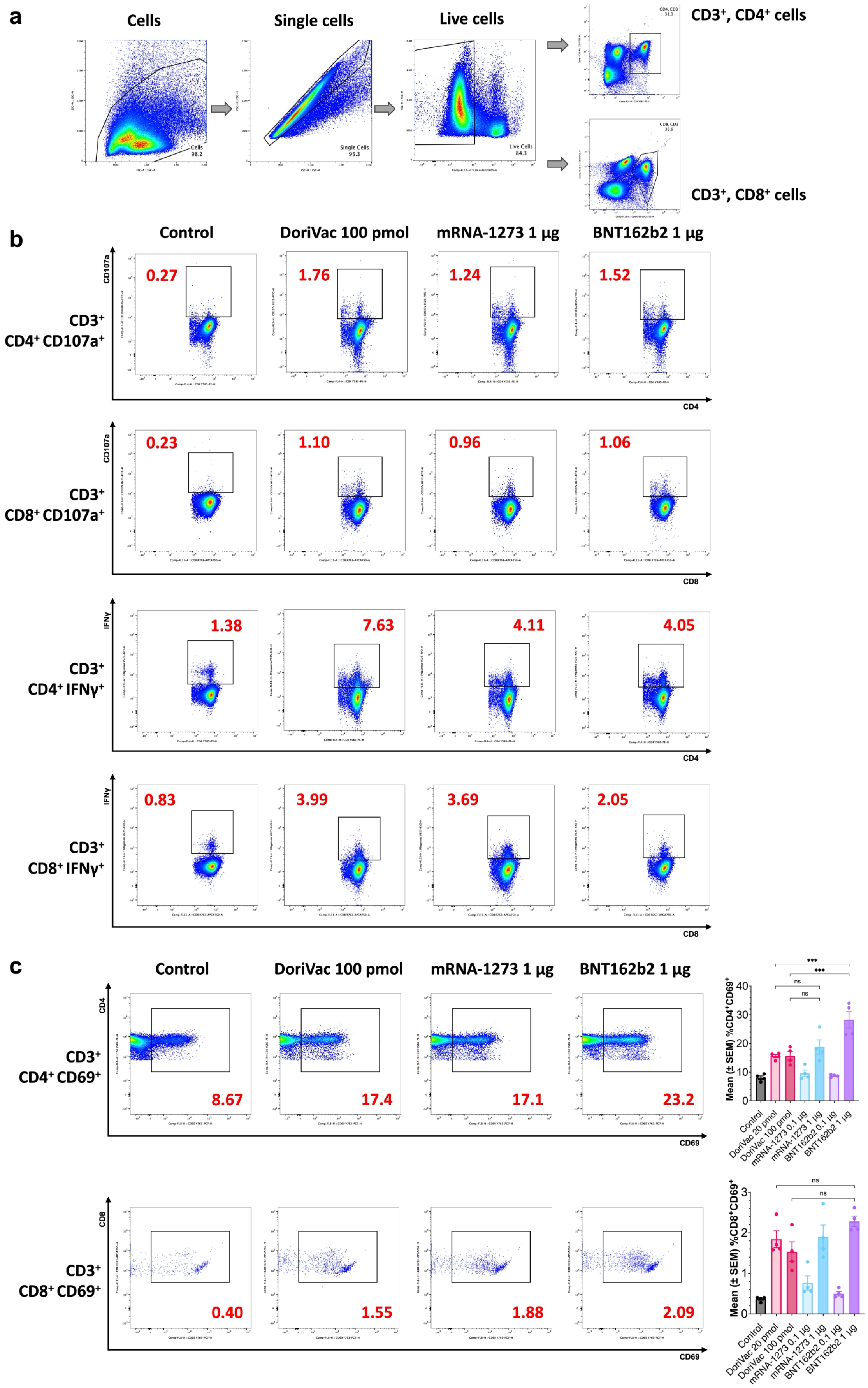


**Supplementary Fig. 18 | Immune activation in CD4⁺ and CD8⁺ effector and IFNγ-secreting T cells on day 50, one day after the third dose. a,** Gating strategy is modified from that illustrated in Supplementary Fig. 9 for the day 50 LN sample to allow for a better separation of CD4⁺ and CD8⁺ LN cells. **b,**

Representative images from flow scatter plots and corresponding frequencies of $CD4^{+}$ and $CD8^{+}$ effector and IFNγ-secreting populations. **c,** $CD69^{+}$ $CD4^{+}$ and $CD8^{+}$ cells (n=4) in the lymph node (LN) were upregulated on day 50 in the DoriVac treatment groups, compared to the untreated control, as determined by flow cytometry. DoriVac treatment groups showed responses comparable to mRNA-LNP groups except the BNT162b2 1 µg group for $CD4^{+}CD69^{+}$ T cells. Representative images from flow scatter plots and corresponding frequencies are shown on the left. The flow data was analyzed by one-way ANOVA (with correction for multiple comparisons using a Tukey's test) and significance was defined as a multiplicity-adjusted p value less than 0.05 (n=4). 'ns' refers to $P > 0.05$; '***' refers to $P \leq 0.001$.

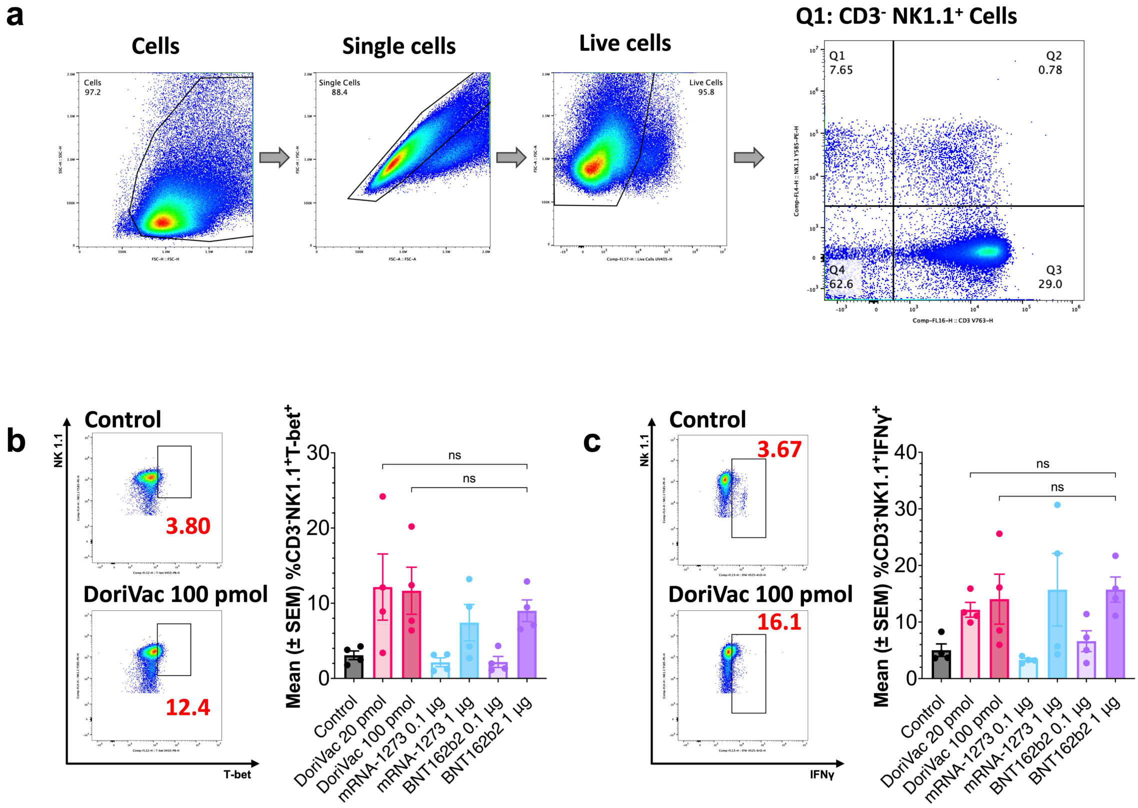


**Supplementary Fig. 19 | Immune activation in PBMC NK Cells on day 50, one day after the third dose.** **a,** Cell gating strategy for CD3$^-$NK1.1$^+$ PBMCs. **b–c,** T-bet and IFNγ expression in NK1.1$^+$ PBMC NK cells on day 50 in DoriVac-treated groups indicates enhanced NK cell activation and proinflammatory response compared to untreated control, as determined by flow cytometry (n=4). DoriVac treatment groups showed responses comparable to mRNA-LNP groups. Representative images from flow scatter plots and corresponding frequencies of population are shown on left. The flow data was analyzed by one-way ANOVA (with correction for multiple comparisons using a Tukey's test) and significance was defined as a multiplicity-adjusted p value less than 0.05 (n=4). 'ns' refers to P > 0.05.

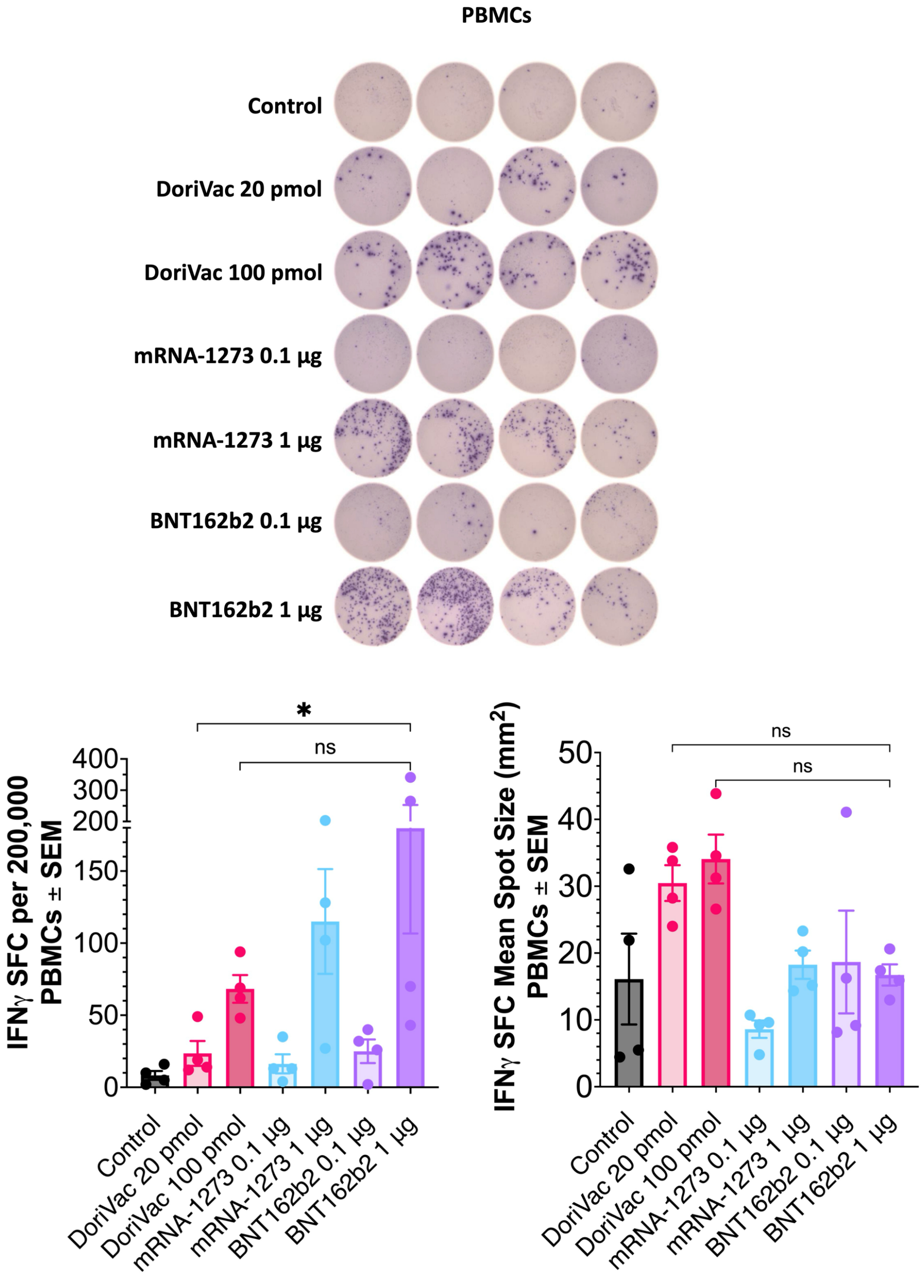


**Supplementary Fig. 20 | SARS-CoV-2-specific T cell response following DoriVac treatment as measured by IFNγ ELISpot on day 35, two weeks after the second dose.** IFNγ ELISpot demonstrating frequency of antigen-specific PBMCs (left, n=4, day 35), with accompanying quantification of IFNγ ELISpot SFUs and mean spot sizes. The results demonstrate an increase in SARS-COV-2 antigen-specific T cell frequency after treatment with DoriVac compared to the control. DoriVac treatment groups showed responses comparable to mRNA-LNP groups, only except when comparing the lower DoriVac dose to the best performing BNT162b2 1 µg group. The data was analyzed by one-way ANOVA (with correction for multiple comparisons using a Tukey's test) and significance was defined as a multiplicity-adjusted p value less than 0.05 (n=4). 'ns' refers to $P > 0.05$; '*' refers to $P \leq 0.05$.

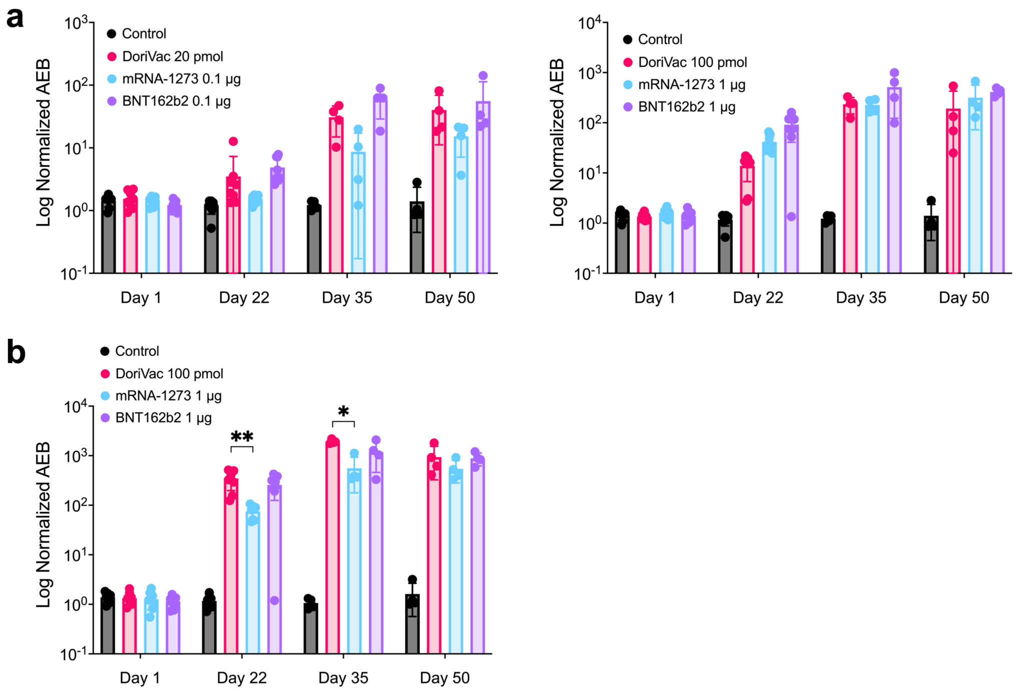


**Supplementary Fig. 21 | Anti-S1 and anti-spike IgG production in different treatment groups as measured by SiMoA. a,** Quantification of relative anti-S1 IgG levels in plasma samples using SiMoA (n=8 on day 1 and day 22, n=4 on day 35 and day 50). DoriVac treatment group exhibited increased antibody levels compared to the control across all sampling dates. **b,** Quantification of relative anti-spike IgG levels in plasma samples of control and 100 pmol/1 µg treatment groups using SiMoA (n=8 on day 1 and day 22, n=4 on day 35 and day 50). DoriVac treatment group exhibited increased antibody levels compared to the control across all sampling dates. The data was analyzed by one-way ANOVA (with correction for multiple comparisons using a Tukey's test) and significance was defined as a multiplicity-adjusted p value less than 0.05. '*' refers to $P \leq 0.05$; '**' refers to $P \leq 0.01$.

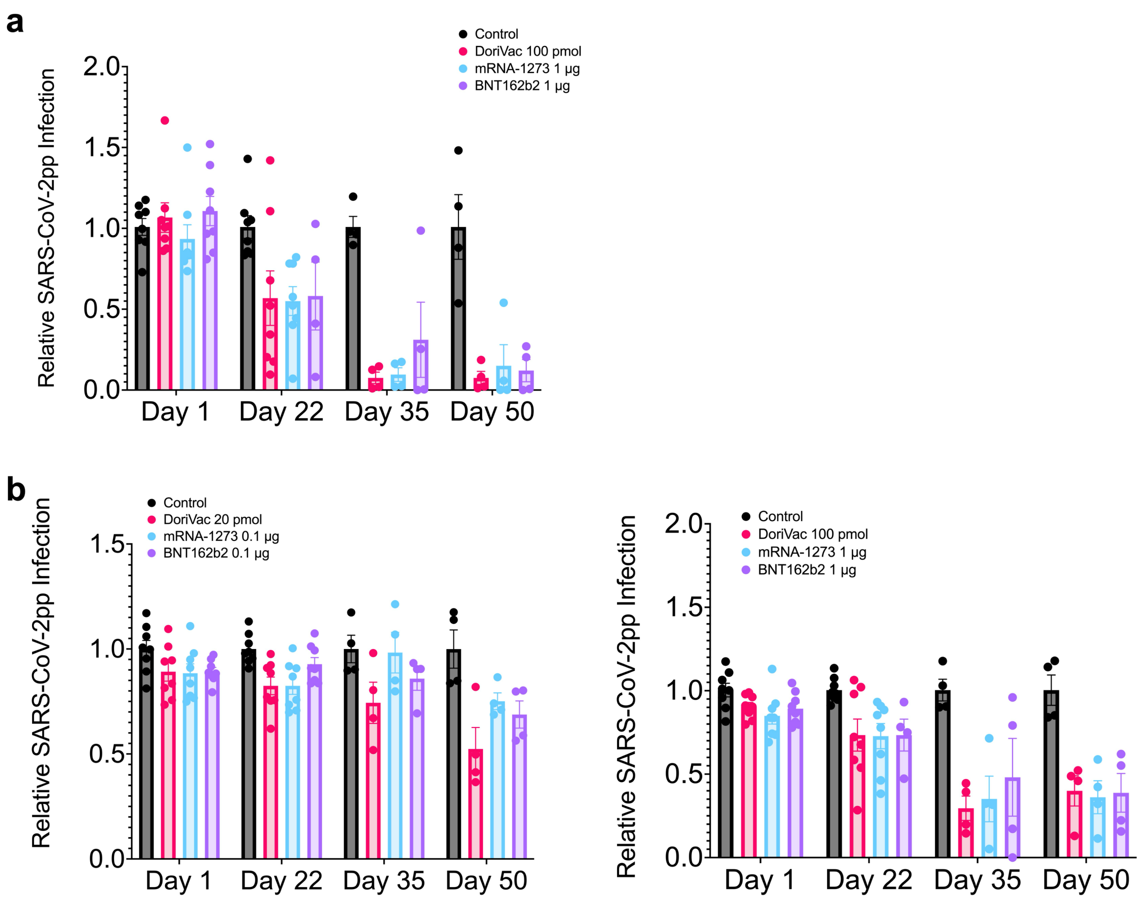


**Supplementary Fig. 22 | SARS-CoV-2 pseudovirus neutralization by different treatments in ACE2-293T cells at low and high doses. a,** SARS-CoV-2 pseudovirus (SARS-CoV-2pp) neutralization assay (1:20 dilution; n=8 on day 1 and day 22, n=4 on day 35 and day 50, except for BNT162b2 1 µg for which n=8 on Day 1 and n=4 on Day 22, 35, 50) for 100 pmol/1 µg treatment groups in model cell line ACE2-293T. DoriVac exhibited more pseudovirus neutralization compared to control starting on day 22. **b,** SARS-CoV-2 pseudovirus (SARS-CoV-2pp) neutralization assays for both 20 pmol/0.1 µg and 100 pmol/1 µg treatment groups were repeated with a 100x serum dilution for all samples. DoriVac continued to show stronger (20 pmol) or comparable (100 pmol) neutralization compared to mRNA-LNP vaccines.

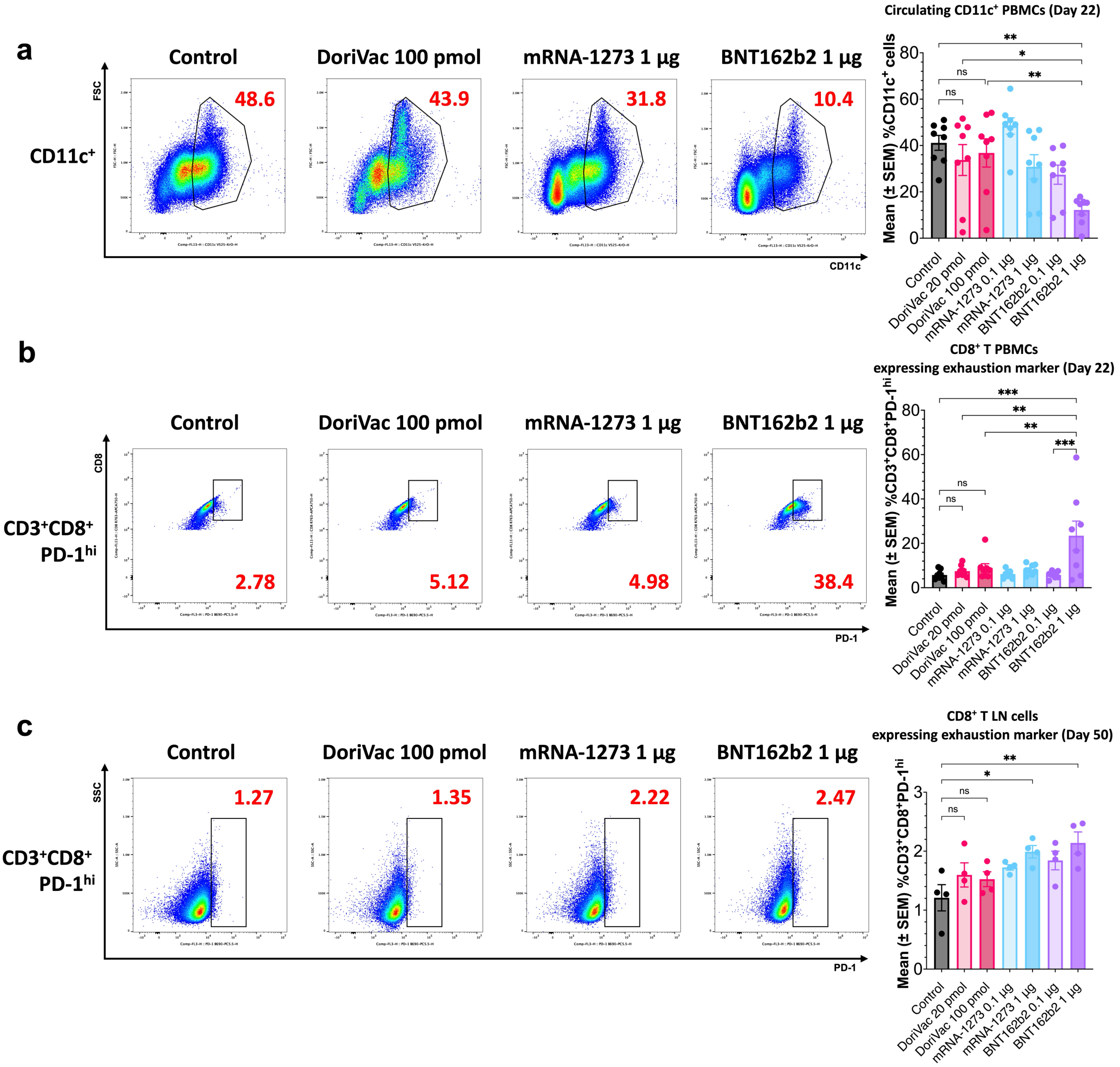


**Supplementary Fig. 23 | Differential immune exhaustion in PBMCs and LN cells across treatment groups. a,** Frequency of circulating CD11c⁺ PBMCs on day 22 across treatment groups. Representative images from flow scatter plots and corresponding frequencies of CD11c⁺ population are shown on the left. Gating strategy is shown in Supplementary Fig. 5. DoriVac treatment groups showed sustained activation, while BNT162b2 1 µg dose exhibited significant reductions compared to the control. **b,** CD8⁺ T cells in PBMCs expressing the exhaustion marker PD-1 on day 22. Representative images from flow scatter plots and corresponding frequencies are shown on the left. Gating strategy is shown in Supplementary Fig. 9. The DoriVac 100 pmol dose group demonstrated significantly lower exhaustion compared to the BNT162b2 1 µg dose group. **c,** CD8⁺ T cells from LN expressing PD-1 on day 50. Representative images from flow scatter plots and corresponding frequencies are shown on the left. Gating strategy is shown in Supplementary Fig. 18. Despite immune activation, DoriVac 100 pmol dose group did not show the same level of PD-1 upregulation as BNT162b2 1 µg dose group. The flow data was analyzed by one-way ANOVA (with correction for multiple comparisons using a Tukey's test) and significance was defined as a multiplicity-adjusted p value less than 0.05 (n=8 for day 22, n=4 for day 50). 'ns' refers to $P > 0.05$; '*' refers to $P \leq 0.05$; '**' refers to $P \leq 0.01$; '***' refers to $P \leq 0.001$.

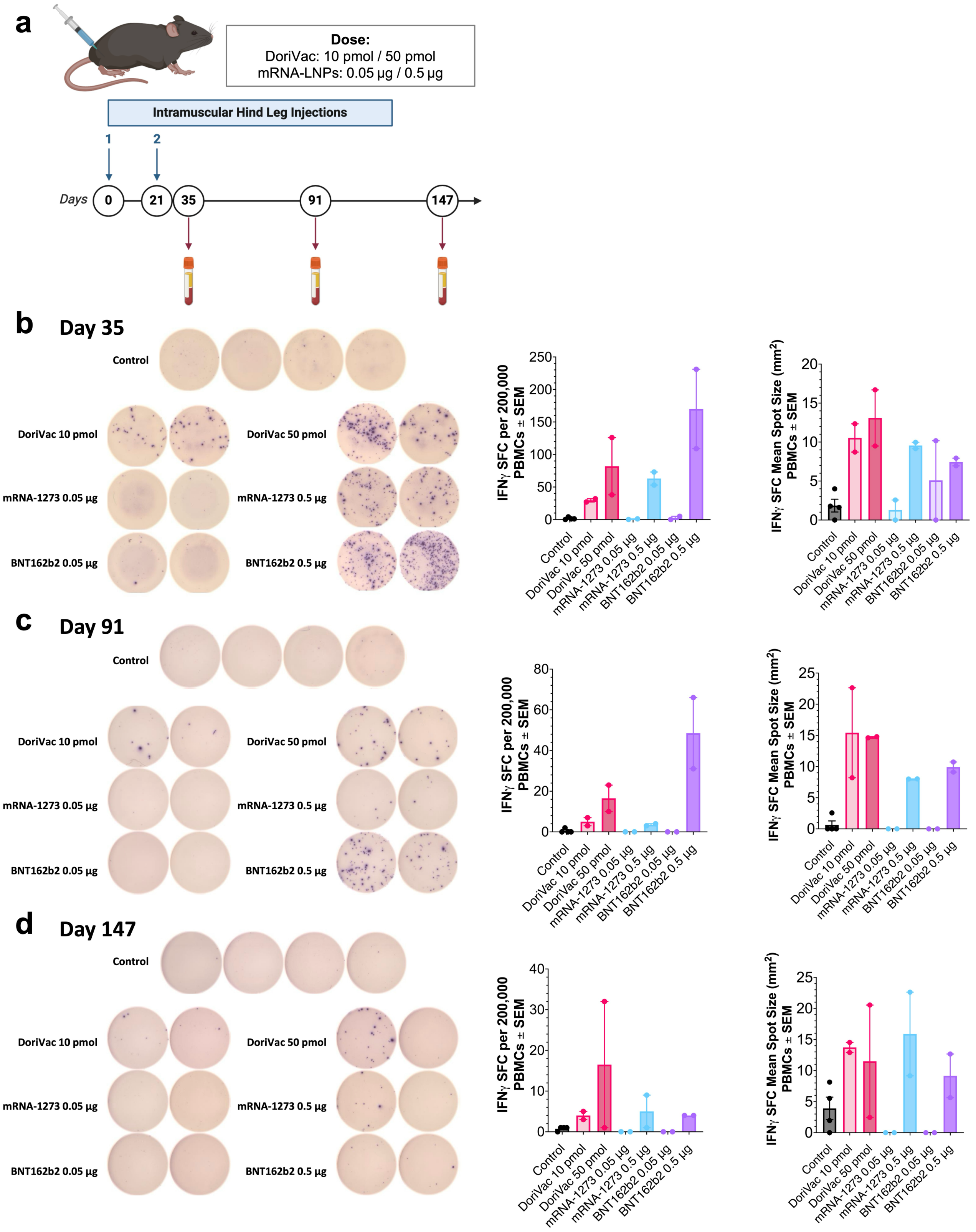


**Supplementary Fig. 24 | SARS-CoV-2-specific T cell frequency and spot size following intramuscularly administered treatments as measured by IFNγ ELISpot. a,** Schematic delineating the intramuscular vaccine administration protocol for naïve C57BL/6 mice and the data collection timeline. Blood samples were collected on day 35, day 91 and day 147. PBMCs were isolated for ELISpot assay. **b–d,** IFNγ ELISpot demonstrating frequency of antigen-specific PBMCs (n=2, except for control n=4) 35 days (2 weeks), 91 days (10 weeks) and 147 days (21 weeks) after first treatment administration, with accompanying quantification of IFNγ ELISpot SFUs and mean spot sizes. Results show an increase in SARS-COV-2 antigen-specific T cell frequency after treatment with DoriVac compared to the control.

DoriVac also produced larger spot sizes than some mRNA-LNP treated groups. The BNT162b2 0.5 µg dose treatment group experienced a large decrease in spot counts between day 91 and day 147, while DoriVac 50 pmol and mRNA-1273 0.5 µg treatment groups remained fairly consistent in counts, suggesting a more sustained T cell response.

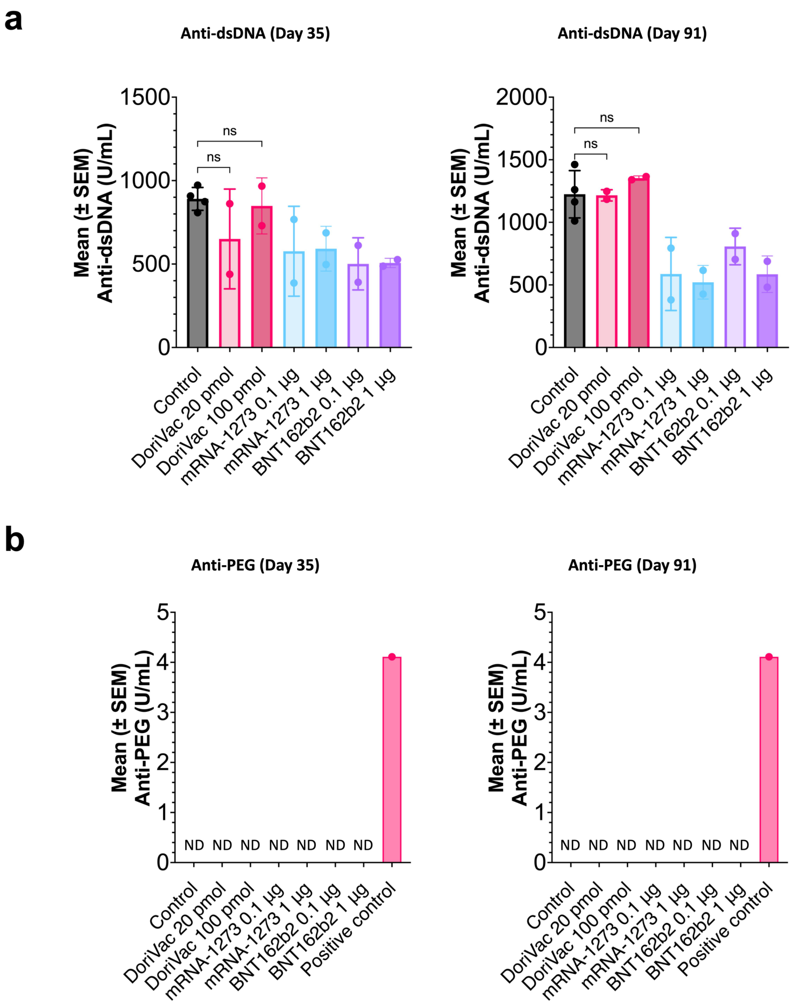


**Supplementary Fig. 25 | Anti-dsDNA and anti-PEG IgG production following intramuscularly administered treatments. a,** Quantification of relative anti-dsDNA IgG levels in serum samples collected on day 35 and day 91 (2 weeks and 10 weeks after second dose; n=2, except for control n=4; all data points are means of technical duplicates). No significant differences were observed between the control and treated groups (ns). **b,** Quantification of relative anti-PEG IgG levels in serum samples collected on day 35 and day 91 (2 weeks and 10 weeks after second dose; n=2, except for control n=4; all data points are means of technical duplicates). Only the positive control exhibited a measurable anti-PEG IgG level, and its value was in the manufacturer-stated range (within 2.8–5.2 U/mL), confirming assay validity. The “U” in U/mL refers to arbitrary activity units, which are defined by the calibrator curves provided by the manufacturer. All other groups were below the limit of detection (ND). All assays were performed with a 1:100 serum dilution, which is the lowest dilution recommended by the manufacturer (≥1:100). The data were analyzed by one-way ANOVA (with Tukey’s correction for multiple comparisons), and significance was defined as a multiplicity-adjusted p value less than 0.05. ‘ns’ indicates $p > 0.05$.

### Supplementary Methods | Determination of Relevant Doses for SARS-CoV-2-Protein-Conjugated DoriVac Compared to mRNA Vaccines

#### Rationale for Dosing Selection

To benchmark the immunogenicity of the DNA–protein hybrid DoriVac against established vaccine platforms, we compared DoriVac doses with those of commercially available SARS-CoV-2 mRNA–lipid nanoparticle (mRNA-LNP) vaccines. Our goal was to identify physiologically relevant doses suitable for mouse models, enabling meaningful comparisons of immune responses, including dendritic cell activation, T cell priming, and antibody generation.

#### Reference mRNA Vaccine Dosing Regimens in Mice

Mice have commonly been used as a preclinical model to evaluate mRNA vaccine immunogenicity and protective efficacy prior to human trials. Studies with mRNA-1273 and BNT162b2 have demonstrated robust immune responses at doses ranging from 0.1 µg to 10 µg per injection, depending on the specific aims and formulation.[1-4] For example, Li et al. reported administering 5 µg of an mRNA-based SARS-CoV-2 vaccine intramuscularly on days 0 and 21 to induce robust neutralizing antibody responses and protection in mice, with lower "human-equivalent" doses approximated at 0.2 µg.[1] Corbett et al. explored a range of doses from 0.01 µg to 10 µg for mRNA-1273, concluding that a 1 µg dose induced robust humoral and cellular responses.[2] Thus, typical murine dosing protocols for SARS-CoV-2 mRNA vaccines often center around 0.1–10 µg per dose, depending on experimental design and the desired magnitude of immune responses.[1,2,5,6]

#### Establishing an Equivalent DoriVac Dose

DoriVac nanoparticle doses were determined based on mole quantities of DNA origami and associated protein antigens. For our SARS-CoV-2 spike-protein-conjugated DoriVac, we employed two primary dose levels—20 pmol and 100 pmol of SQB (DNA origami "square block") constructs per injection. Each DoriVac particle displays multiple spike protein molecules, providing high antigen density. The 20 pmol dose corresponds to approximately $1.2 \times 10^{13}$ DNA origami particles, and the 100 pmol dose is fivefold higher. Assuming a nominal occupancy of ~5 spike proteins per SQB, the 20 pmol dose presents on the order of ~$6 \times 10^{13}$ spike protein copies. By comparison, mRNA vaccine doses rely on endogenous translation of administered mRNA. Data from mRNA expression studies suggest that 1 µg of mRNA can yield on the order of $10^{13}$ to $10^{14}$ spike protein copies per injection, though actual yields vary with mRNA stability, translational efficiency, and delivery conditions.[7-10] Notably, this calculation is an approximation and may not directly translate into equivalent immunogenicity due to differences in mRNA protein translation, antigen processing, and localization. These mRNA copy estimates are derived from publicly available SARS-CoV-2 spike-encoding mRNA vaccine sequences.[11] Using these reference sequences, approximately 4–5 × $10^{11}$ mRNA molecules are present in 1 µg of mRNA.

#### Comparisons and Justification of Selected Dose Range

While it is tempting to directly compare the number of antigen molecules delivered by DoriVac with that theoretically produced from mRNA translation, such comparisons must be interpreted with caution. mRNA-LNP vaccines incorporate proprietary structural elements and modified nucleotides to enhance translation and stability, resulting in highly efficient protein production.[1,2,12] Conversely, DoriVac directly delivers pre-formed proteins to antigen-presenting cells. Thus, differences in antigen presentation pathways, kinetics of antigen availability, and immune-processing mechanisms complicate straightforward equivalences between mRNA and protein doses.

Furthermore, while mRNA vaccine doses of 0.1–1 µg are frequently sufficient for robust immunogenicity in mice[1,2], these doses rely on the host's translational machinery and may differ substantially from direct protein delivery platforms. mRNA vaccines continuously produce antigen for as long as the mRNA remains stable and efficiently translated. In contrast, DoriVac delivers the full antigen complement at once, potentially affecting dose-response kinetics.

Previous studies estimating the number of proteins translated per mRNA molecule have proposed median translation rates and protein half-lives on the order of hours.[7-9] Using a median estimate of ~1,800 proteins generated per mRNA transcript[7], 1 µg of mRNA (comprising ~4–5 × $10^{11}$ mRNA copies for SARS-CoV-2 spike) might yield on the order of ~8–9 × $10^{14}$ proteins. Independent *in vitro* data corroborate this scale, measuring ~6 × $10^{14}$ proteins per µg of spike-encoding mRNA.[13] Such estimates provide useful ballpark figures but are not definitive benchmarks, as *in vivo* translation efficiency is influenced by numerous factors, including cell type, tissue microenvironment, and vaccine formulation.

Our chosen DoriVac doses (20 pmol and 100 pmol) reflect a balance between delivering sufficient antigen for immune priming and maintaining practical nanoparticle concentrations. In our previous study, the 20 pmol DoriVac dose elicited a robust immune response against the antigen and demonstrated favorable efficacy in the tumor model.[14] Analogous to how mRNA vaccine researchers have refined dosing regimens over time, future studies may optimize DoriVac dosing by incorporating a wider range of doses and dosing intervals, as well as evaluating alternative routes of administration.

### Conclusions and Future Directions

Determining a "human-equivalent" dose or a direct functional comparison between protein-bearing DNA origami and mRNA vaccines is inherently challenging. Due to the differences in delivery modalities, antigen processing, and immunobiology, numerical equivalences in antigen copy number should serve only as rough guides. The DoriVac doses applied here were chosen to allow a fair comparison against established mRNA-LNP vaccine data. Further exploration—such as parallel dosing studies, long-term immunogenicity assessments, and direct comparisons under identical experimental conditions—will be needed to fine-tune DoriVac dosing strategies and fully elucidate its immunogenic potential relative to current vaccine technologies.
